# Nuclear RNA concentration coordinates RNA production with cell size in human cells

**DOI:** 10.1101/2021.05.17.444432

**Authors:** Scott Berry, Micha Müller, Lucas Pelkmans

## Abstract

Unlike its DNA template, RNA abundance and synthesis rates increase with cell size, as part of a mechanism of cellular RNA concentration homeostasis. Here, we study this scaling phenomenon in human cells by combining genome-wide perturbations with quantitative single-cell measurements. Despite relative ease in perturbing RNA synthesis, we find that RNA concentrations remain highly constant. Systems-level analysis indicates that perturbations that would lead to increased nuclear mRNA abundance result in downregulation of mRNA synthesis. This is associated with reduced levels of several transcription-associated proteins and protein states that are normally coordinated with RNA production in single cells, including RNA polymerase II (Pol II) itself. Acute shut-down of nuclear RNA degradation, elevation of nuclear mRNA levels, and mathematical modelling indicate that mammalian cells achieve RNA concentration homeostasis by an mRNA-based negative feedback on transcriptional activity in the nucleus. Ultimately, this acts to robustly scale Pol II abundance with cell volume and coordinate mRNA synthesis and degradation.

## INTRODUCTION

Within a cell population, the number of mRNA transcripts of a given gene is highly variable between cells (Raj and Oudenaarden, 2008). Yet much of this variability can be explained by variation in cell size – with larger cells containing more transcripts (Battich et al., 2015; Padovan-Merhar et al., 2015). Such ‘scaling’ of mRNA abundance with cell size is highly conserved and is underpinned by increasing transcript production rates with cell size, while degradation rates remain constant (Ietswaart et al., 2017; Kempe et al., 2015; Padovan-Merhar et al., 2015; Sun et al., 2020; Zhurinsky et al., 2010). However, despite these consistent observations across eukaryotes, we still lack a mechanistic understanding of how cells control global RNA production, and how this is coordinated with cell size. The predominant mechanistic hypothesis is the ‘limiting factor model’, which supposes that cells contain a factor that is limiting for transcription and whose abundance is closely coupled to cell size (Marguerat and Bähler, 2012). If this limiting factor is localised to a cellular structure that does not scale with cell size (e.g. DNA), then its *local* concentration on that structure will be proportional to cell size, despite having constant *global* concentration (Padovan-Merhar et al., 2015). It has been suggested that RNA Polymerase II (Pol II) itself is the limiting factor (Lin and Amir, 2018; Padovan-Merhar et al., 2015; Sun et al., 2020), but causal evidence is lacking.

Control of RNA degradation has thus far not been implicated in cell size-scaling, however studies in yeast have revealed that disruption of RNA degradation tends to result in compensatory decreases in RNA synthesis rates, and vice versa, that disrupting transcription leads to decreases in RNA degradation rates (Haimovich et al., 2013; Sun et al., 2012, 2013). In yeast, this ‘buffering’ mechanism was reported to involve direct (Haimovich et al., 2013) or indirect (Sun et al., 2013) transcriptional regulation by the cytoplasmic 5’-3’ exonuclease Xrn1. In mammalian cells, similar systematic studies of mRNA buffering are lacking (reviewed in (Hartenian and Glaunsinger, 2019)), although there have been reports of mRNA stabilisation upon transcriptional inhibition (Helenius et al., 2011; Slobodin et al., 2020). In contrast to yeast, however, both acceleration (Abernathy et al., 2015; Gilbertson et al., 2018) *and* disruption of cytoplasmic RNA degradation (Lee et al., 2012; Singh et al., 2019) have been associated with transcriptional repression. The role of the cytoplasm in mRNA buffering therefore remains unclear in mammalian cells. Moreover, in both yeast and mammalian systems, studies of mRNA buffering have focused on cell population measurements and have not considered effects of cell size changes in perturbations, complicating the interpretation of their effects.

Since both buffering and size-scaling contribute to mRNA concentration homeostasis, they may be part of the same mechanism, but whether this is the case, and how this works is unknown. To study these phenomena together, we here combine genomewide genetic perturbation screening with multiplexed quantitative measurements of single cells. Crucially, this enables us to connect effects seen in perturbations with naturally varying properties of unperturbed cells. We uncover hundreds of perturbations that affect global RNA synthesis rates in single cells, including pathways not previously implicated in transcriptional control. However, in most cases we find that RNA concentration is not disrupted. Systems-level analysis of the genes involved together with detailed characterisation of molecular phenotypes at the single-cell level suggest a model in which transcription rates are negatively regulated by *nuclear* mRNA concentration. We propose that the activity, and ultimately abundance, of RNA Pol II is determined by this mRNA-based feedback, to enable robust mRNA concentration homeostasis in human cells.

## RESULTS

### Cell size perturbation leads to precise adaptation of mRNA abundance

In unperturbed human cells, mRNA abundance scales with cell size (Figure S1A) (Battich et al., 2015; Kempe et al., 2015; Padovan-Merhar et al., 2015). To investigate if mRNA concentrations are robustly maintained upon cell volume perturbation, we established a method of cell volume measurement compatible with high-throughput branched-DNA (bDNA) smFISH (Figures S1B-H, STAR methods) and measured cytoplasmic transcript abundance for 14 size-scaling genes (Battich et al., 2013) in populations of HeLa cells. Cell size was perturbed using siRNA-mediated knockdown of GRIP2 and SBF2, genes which are not known to affect transcription but whose knockdown results in smaller and larger cells, respectively (Berchtold et al., 2018) 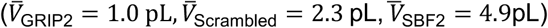.

Average mRNA abundance was dramatically reduced in smaller cells (GRIP2 RNAi) and increased in larger cells (SBF2 RNAi) (Figure 1A-B, S1I). However, mRNA abundance remained proportional to cell volume, and volume typically still accounted for the majority of transcript abundance variation (Figure S1I-L). Moreover, mean mRNA abundance in perturbations was well predicted by the change in volume (Figure S1M) and normalising spot count by volume at the single-cell level led to overlapping distributions for most genes (Figure 1B, S1N). Cell volume is therefore a dominant source of heterogeneity in mRNA abundance in cell populations, and its perturbation leads to precise adaptation of transcript abundance to maintain genespecific mRNA concentration.

**Figure 1:**
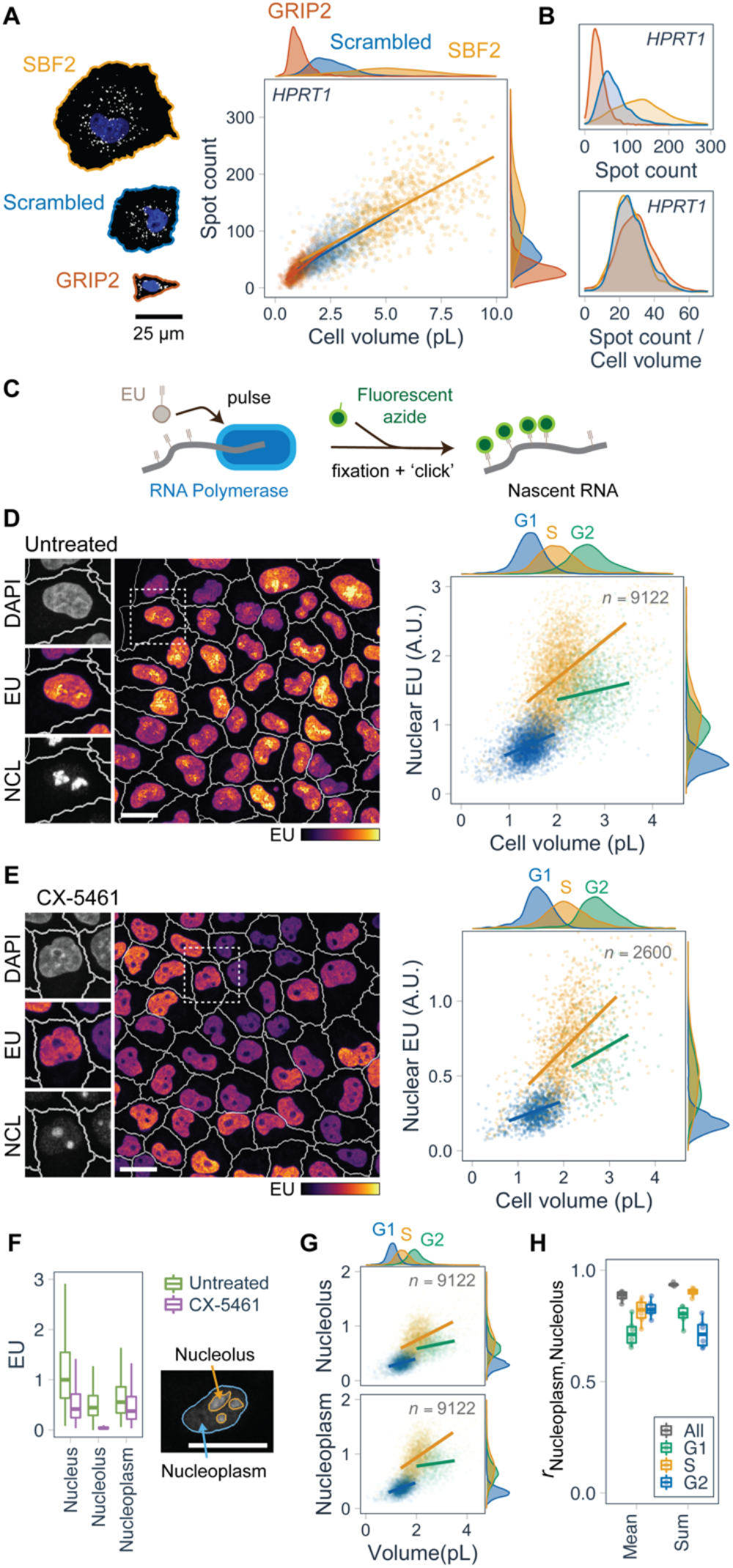
RNA abundance and production rates scale with cell size. (A) Number of cytoplasmic *HPRT1* transcripts detected by smFISH as a function of cell volume, in cells transfected with scrambled siRNA or siRNA targeting GRIP2 or SBF2. DAPI (blue) and *HPRT1* smFISH (grey) for selected example cells. (B) Single-cell spot count distribution for *HPRT1*, normalised by cell volume in lower panel. (C) Metabolic RNA labelling using EU. (D) EU incorporation in untreated cells during a 30 min pulse. Nucleolin (NCL) immunofluorescence, EU and DAPI inset. Quantification of single-cell sum nuclear EU intensity as a function of cell volume, for different cell cycle stages. (E) As in D, with 2h CX-5461 pre-treatment. (F) Sum EU intensity in subnuclear regions in CX-5461-treated and untreated cells. Boxplots summarise single-cell values with outliers omitted for clarity. Values relative to median nuclear EU intensity of untreated cells. (G) As in D, for sum nucleolar and sum nucleoplasmic EU. (H) Correlation of nucleolar and nucleoplasmic EU, for either the sum or the mean of pixel values. Boxplots summarise correlations observed in individual replicate wells. All data from HeLa cells. Scale bars 25μm. See also Figures S1-S2.

### RNA production rates are coupled to cell cycle and cell volume

RNA abundance is determined by production and degradation rates. However, inferring gene-specific mRNA production rates in single cells is technically challenging and often requires assumptions (Ietswaart et al., 2017; Padovan-Merhar et al., 2015; Sun et al., 2020). Because size-scaling is a transcriptome-wide phenomenon, we measured bulk RNA production in single cells *in situ* using metabolic pulse labelling with 5-ethynyl uridine (EU) (Jao and Salic, 2008; Padovan-Merhar et al., 2015; Shah et al., 2018) in combination with measurement of cell volume and stains to assign cells to G1/S/G2 cellcycle phases (Figure 1C-D, S2A-J). In two cell lines (HeLa, 184A1) and in primary human keratinocytes, EU incorporation increased with cell volume (Figure S2I), in agreement with previous results (Padovan-Merhar et al., 2015). However, including cell-cycle information revealed that G2 and S-phase cells showed higher EU incorporation than G1 cells of the same volume (Figure 1D, S2J). Furthermore, HeLa cells showed an additional increase during S-phase (Pfeiffer and Tolmach, 1968) – above the level seen in G2. While these cell cycle effects are interesting and suggest differences compared to yeast (Voichek et al., 2016), we here focus on cell-size scaling of EU incorporation – using precise cell cycle information to exclude changes to DNA template abundance.

EU is incorporated into all major RNA species by RNA polymerases I, II, and III (Jao and Salic, 2008). To evaluate the contribution of ribosomal RNA (rRNA) to EU incorporation, we treated cells with the RNA Pol I inhibitor CX-5461 (Drygin et al., 2011), which resulted in elimination of EU incorporation in the nucleolus – the site of rRNA transcription (Figures 1E, S2G-H).

To quantify this, we segmented the nucleolus and nucleoplasm (non-nucleolus) and summed EU intensity in each region separately (Figure S2L). CX5461 reduced nucleolar EU by 89-92% while the reduction in total nuclear EU was more modest (46-56%) (Figures 1F, S2M). Moreover, nucleolar segmentation in untreated cells revealed that the nucleolus contributes 45% of nuclear EU incorporation. Taken together, this suggests that rRNA contributes approximately half of the EU incorporation in nascent RNA. This is less than the contribution of rRNA to total RNA abundance (80% (Wolf and Schlessinger, 1977)) and is consistent with much greater stability of rRNA than mRNA (estimated average half-life 3-8d (Gillery et al., 1995; Halle et al., 1997) and 3.5h, respectively (Herzog et al., 2017; Tani et al., 2012)). Cell-volume and cellcycle dependence of total nuclear EU incorporation was similar between CX-5461-treated and untreated cells (Figures 1E, S2K). Moreover, in untreated cells, EU intensities in the nucleolus and nucleoplasm were highly correlated and showed similar cell-volume and cell-cycle dependence (Figure 1G-H).

To investigate how increases in cell size affect RNA production, we also treated HeLa cells with the CDK inhibitor roscovitine for 48h (Cadart et al., 2018). This led to an approximate doubling of cell volume in each cell cycle phase, without affecting the cell-cycle distribution (Figure S2N-O). EU incorporation increased approximately in proportion to cell volume changes. This demonstrates that perturbing cell volume results in coordinated adaptation of bulk RNA production rates in human cells, in agreement with studies at the cell-population level in yeast (Fraser and Nurse, 1979; Zhurinsky et al., 2010).

### Genetic perturbation screen reveals large changes in RNA production rates

To investigate the genetic control of RNA production rates and their coordination with cell size, we next applied this RNA metabolic labelling assay in the context of an arrayed genome-wide siRNA perturbation screen (Figure 2A) (Müller et al., 2021). The screen comprises spatially resolved multivariate measurements of ~80 million HeLa cells across 21,823 perturbations, providing a comprehensive resource to perform systems-level analyses of the regulation of RNA production as a function of cell size and cell-cycle stage. Because cellular protein content and nuclear area are both proportional to cell volume (Cantwell and Nurse, 2019; Kafri et al., 2013) (Figure S1G), we used these as measures of cell size in the screen.

**Figure 2:**
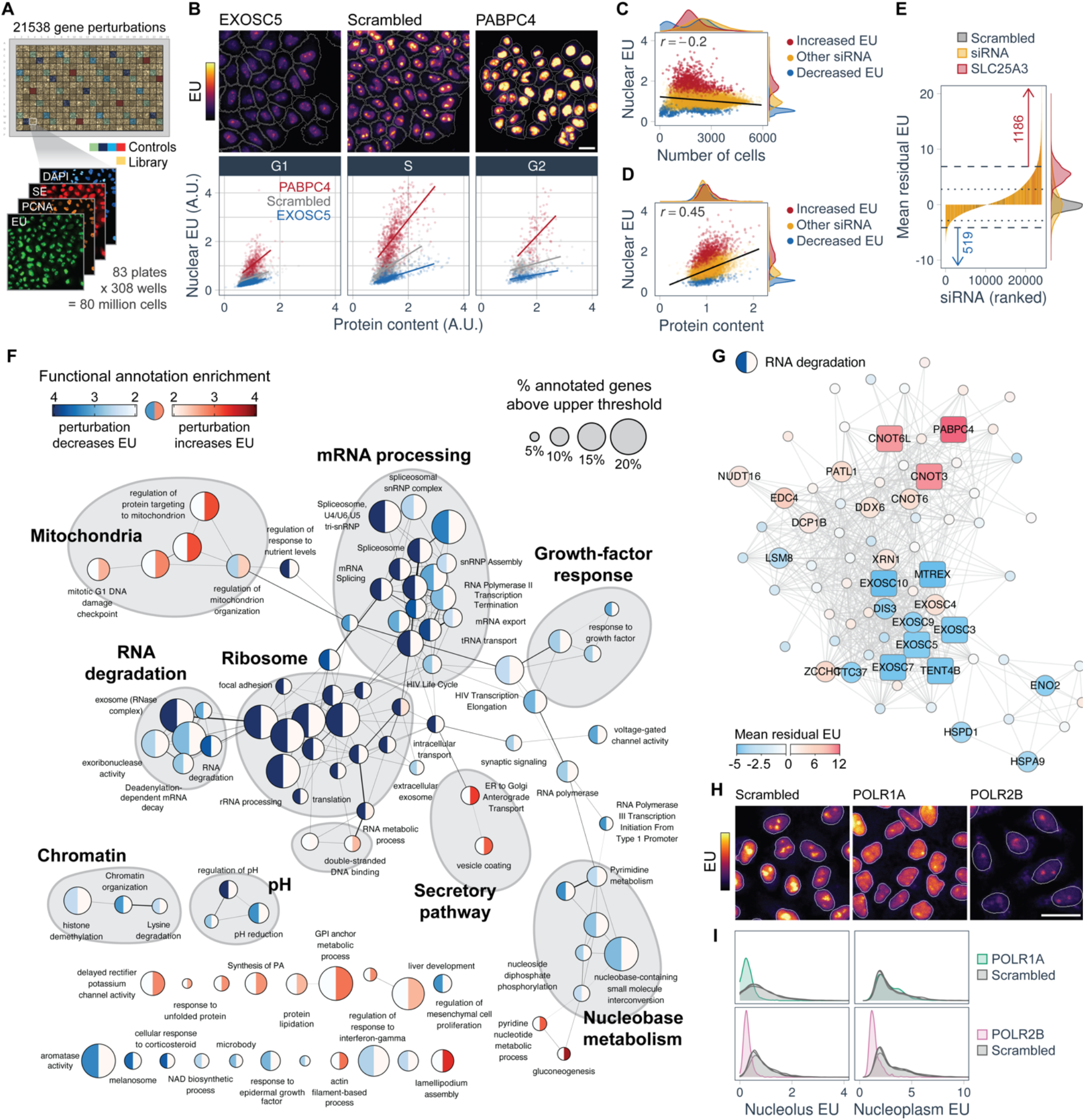
Genetic screen identifies regulators of RNA production. (A) Arrayed genome-wide siRNA screen. (B) Upper: example images of EU metabolic labelling from the screen. Lower: sum nuclear EU intensity as a function of cellular protein content, for each cell cycle phase. (C) Perturbation-averaged sum nuclear EU as a function of number of interphase cells in the screen. (D) Perturbation-averaged sum nuclear EU as a function of cellular protein content (G1 cells only). (E) Mean residual EU across the screen for scrambled siRNA (n=1,826), SLC25A3 (n=664) and library siRNAs (n=21,538). Gene perturbations ranked by mean residual EU. Dotted and dashed lines indicate lower and upper hit thresholds (*p*_posterior_ = 0.5,0.85), respectively (Müller et al., 2021). (F) Network of functional annotations enriched in perturbations with increased or decreased mean residual EU (STAR methods). (G) STRING protein-protein association network of genes with RNA degradation annotation. Conditions with low cell number omitted. Labelled circles for *p*_posterior_ > 0.5 and squares for *p*_posterior_ > 0.85. (H) Images of EU incorporation in the genomewide screen for POLR1A or POLR2B knockdown. (I) Single-cell distributions of sum nucleoplasmic and nucleolar EU for perturbations in H. Three scrambled siRNA wells from the same plates shown for comparison. All scale bars 25μm. See also Figures S3-S5.

A large number of perturbations led to increases or decreases in EU incorporation (Figure 2B), with changes often of a large magnitude (mean EU foldchange percentiles *P*_1_ = 0.56 and *P*_99_ = 2.2) (Figure S3A). While some perturbations were associated with reduced viability, the general trend was weak (Figure 2C). Perturbation-averaged EU incorporation correlated with cellular protein content across the screen (Figure 2D), further demonstrating that altered cell size typically leads to a coordinated change in RNA production. Furthermore, EU incorporation increased with cellular protein content at the single-cell level in all perturbations (Figure S3E-G). To systematically identify perturbations of EU incorporation, we therefore derived a measure of EU intensity that is corrected for cell size and cellcycle stage at the single-cell level (STAR Methods).

We refer to this as ‘mean residual EU’. It is negative for perturbations with reduced EU incorporation (n=519 hits, 413 of which have >500 cells) and positive for those with increased EU incorporation (n=1186 hits, 1183 of which have >500 cells) (Figure 2E). Mean residual EU hits are not associated with characteristic cell-cycle distribution or cell-size changes (Figure S3C).

To reveal the functions of genes whose perturbation underlies altered EU incorporation, we performed rank-based enrichment analysis of functional annotations using mean residual EU (STAR methods). Enrichment scores were higher for the ‘down’ hits than ‘up’ hits, suggesting that reduction in EU incorporation occurs through disruption of a more focused (or better annotated) set of pathways. As expected, ribosomal biogenesis (Pol I), as well as Pol II- and Pol III-dependent transcription were strongly associated with reduced EU incorporation (Figure 2F). However, nuclear RNA processing and splicing, chromatin organisation, nuclear RNA export, and RNA degradation annotations were also strongly enriched for reduced EU incorporation. Closer examination of the genes underlying these annotations (Figures 2G, S3H, S4) revealed, for example, that enrichment of the term ‘RNA degradation’ for reduced EU incorporation was driven mostly by the *nuclear* rather than *cytoplasmic* RNA decay factors, especially the RNA exosome (a multicomponent 3’ to 5’ ribonuclease (Schmid and Jensen, 2018)), and nuclear exosome targeting factors such as MTREX, ZFC3H1 and TENT4B (Figure S5A). Annotations enriched for *increased* EU incorporation include the secretory pathway – specifically vesicle coating and ER to Golgi anterograde transport (for example RAB1A, SEC16B, TRAPPC1), and also protein lipidation, especially genes involved in GPI-anchor biosynthesis (for example PIGP, DPM3) (Figures S3H, S4). These novel phenotypes suggest intriguing links between cell surface homeostasis and transcriptional regulation. EU incorporation increases were also observed upon perturbation of many sequencespecific DNA binding proteins, for example NFX1/NFXL1, and ELK1 and also histone demethylases such as KDM2B (H3K4/K36-demethylase) (Figures S3H, S4), indicating that individual transcription factors and histone modifiers can also have strong repressive effects on overall RNA synthesis.

Nucleolar and nucleoplasmic RNA production are highly coordinated in single cells (see Figure 1H). Across the genome-wide screen, we also observed a high correlation between perturbation-averaged mean EU intensities in the nucleolus and nucleoplasm (*r* = 0.97), with sum EU intensities slightly less well correlated (*r* = 0.86) (Figure S5B), due to changes in nucleolus size (Boulon et al., 2010). This suggests that perturbations typically have similar effects on ribosomal and non-ribosomal transcription. Despite this general correspondence, however, we did identify a subset of hits which specifically reduced nucleolar but not nucleoplasmic EU, many of which were related to Pol I-dependent transcription (Figures S5C-J, STAR Methods). At face value this may be unsurprising, however we did *not* observe a converse nucleoplasm-specific EU reduction when targeting genes associated with RNA Pol II transcription (Figure S5K). For example, POLR1A RNAi specifically affected nucleolar EU while POLR2B RNAi affected both the nucleolus and the nucleoplasm (Figure 2H-I). This is consistent the role of Pol II in promoting Pol I-dependent transcription (Abraham et al., 2020; Burger et al., 2013; Caudron-Herger et al., 2016) and suggests that size-scaling of Pol I-dependent transcription may occur as a consequence of size-scaling of Pol II-dependent transcription.

### RNA concentration is stable in perturbations with altered RNA production

To determine how RNA abundance is affected in conditions in which synthesis rates are perturbed, we measured mRNA abundance using fluorescence *in situ* hybridisation against polyadenylated RNA (poly(A) FISH) and total RNA abundance using RNA StrandBrite – a fluorescent stain specific for RNA. Both stains were visible in the nucleus and cytoplasm, with poly(A) FISH enriched in nuclear speckles and RNA StrandBrite enriched the nucleolus (Figure 3A, S6A). Both were strongly correlated with cellular protein content (Figure S6D).

**Figure 3:**
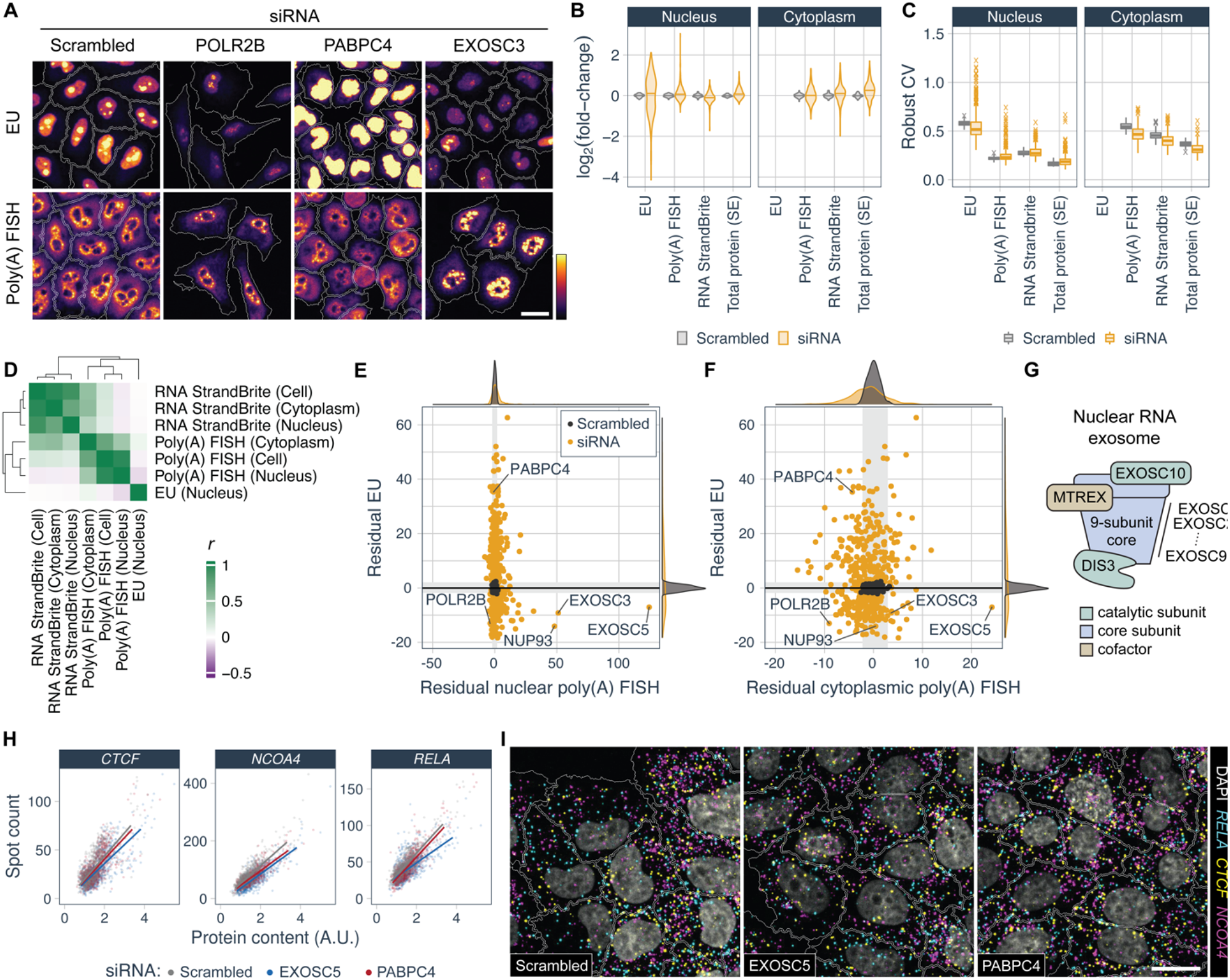
RNA concentration is stable in conditions with altered RNA production. (A) EU metabolic labelling and poly(A) FISH images for selected siRNA perturbations. (B) Fold-change in perturbation-averaged mean nuclear and cytoplasmic intensities relative to scrambled siRNA controls. Scrambled siRNA (n=80), library siRNA (n=415). Low cell number conditions omitted. (C) Cell-to-cell variability (robust coefficient of variation) of mean nuclear and cytoplasmic intensities within each well, for data in B. (D) Correlations of mean residual RNA abundance measurements and EU (n=379). (E) Mean residual EU versus mean residual nuclear poly(A) FISH. Mean of two replicates shown. Grey boxes indicate 1st/99th percentiles of scrambled siRNA controls. (F) As in E, for cytoplasmic poly(A) FISH. (G) Nuclear RNA exosome schematic. (H) Cytoplasmic transcript abundance measured by smFISH for *CTCF*, *NCOA4* and *RELA* transcripts as a function of cellular protein content. Example siRNA perturbations shown together with scrambled siRNA controls. (I) Example smFISH images. All scale bars 25μm. See also Figures S6-S7.

RNA abundance measurements were performed on a set of 436 gene perturbations, which were chosen from enriched annotations in the genome-wide screen to maintain both functional and phenotypic diversity (Figure S6B) (Müller et al., 2021). Across this set of perturbations, we observed that mRNA and total RNA intensities were much less strongly perturbed than EU intensities (Figures 3B, S6E), implying that RNA concentration homeostasis is not generally disrupted. For example, POLR2B and PABPC4 knockdown resulted in a 5.7-fold reduction or 2.6-fold increase, respectively, in EU incorporation relative to scrambled siRNA controls, but both showed a 1.1- or 1.2-fold reduction in mRNA abundance in the nucleus and cytoplasm, respectively (Figure 3A). Moreover, the variability *within* cell populations (robust coefficient of variation) was also greater for EU incorporation than protein, mRNA and total RNA abundance (Figure 3C).

To systematically identify changes in RNA abundance, we first corrected for cell size and cell cycle changes using linear regression, which led to narrower distributions of perturbation-average poly(A) FISH and RNA StrandBrite intensities, and greater overlap with scrambled siRNA controls (Figure S6E-F). This indicates that cell size and cell cycle changes explain some of the differences that we see in RNA abundance (Figure S6G). We then compared these RNA abundance measurements with EU incorporation across perturbations (Figures 3D, S6H). Average total RNA and mRNA abundances were moderately correlated with one another across perturbations, however there was a conspicuous lack of strong correlation of either with EU (Figures 3D-F, S6H-K). This implies that in human cells, RNA degradation rates generally change in concert with production rates, similar to yeast (Haimovich et al., 2013; Sun et al., 2012). In the case of nuclear mRNA abundance, we even observed a weak but significant anti-correlation (*r* = −0.08, *p* = 0.02) driven by a small group of perturbations that show reduced EU incorporation, and strong overaccumulation of mRNA in the nucleus (Figures 3E, S6G). This disruption of mRNA concentration homeostasis occurs when targeting nuclear pore component NUP93 or core components of the RNA exosome: EXOSC3, EXOSC5 (Fan et al., 2018; Silla et al., 2018), revealing the critical roles played by nuclear RNA degradation and export. In addition to nine core subunits (EXOSC1-EXOSC9), the nuclear RNA exosome (Figure 3G) contains two catalytic subunits: DIS3, which targets products of Pol II transcription, and EXOSC10 which plays a role in nucleolar RNA processing (Davidson et al., 2019; Schmid and Jensen, 2018). Although both EXOSC10 and DIS3 showed reduced EU incorporation in the genome-wide screen (Figure 2G), their knockdown did not lead to increased nuclear mRNA levels (Figure S6G), likely because of partial functional redundancy between them (Davidson et al., 2019; Fan et al., 2018; Tomecki et al., 2010).

To measure gene-specific cytoplasmic mRNA abundance, we applied bDNA smFISH to detect transcripts for nine genes across 50 genetic perturbations (Figures 3H-I, S7A-F). Perturbations included RNA exosome and nuclear pore components, transcription machinery and splicing factors, as well as diverse perturbations with increased EU incorporation. Although genetic perturbations often led to changes in gene-specific transcript abundance, changes were not in a consistent direction for a given perturbation (Figure S7E, S7G), and transcript abundance of specific genes was typically not correlated with EU incorporation across perturbations (Figure S7H-I). Moreover, RNA abundance remained highly coordinated with cellular protein content at the singlecell level (Figure 3H, S7E), indicating maintenance of gene-specific mRNA concentration.

Together, these data reveal that RNA concentrations are generally maintained in conditions with altered EU incorporation, pointing to a generic coordination of RNA synthesis and degradation in human cells.

From the hundreds of conditions studied, we identified only a few exceptions in which mRNA concentration homeostasis was strongly disrupted – specifically in the nucleus. These perturbations involve core components of the nuclear RNA exosome and nuclear pore (EXOSC3, EXOSC5, NUP93), and all show strong accumulation of nuclear poly(A) FISH signal and reduced EU incorporation. Since perturbation of nuclear mRNA processing and export are also associated with reduced EU incorporation in the genome-wide screen, this points to a mechanism by which cells down-regulate RNA synthesis in response to an inability to export or degrade nuclear mRNA.

### Characterization of molecular changes underpinning perturbation of RNA production

The diverse set of genetic perturbations identified that affect EU incorporation provide an opportunity to investigate how cells globally regulate bulk RNA production. To determine how the abundance of Pol II and other proteins involved in RNA production and processing are altered when RNA production rates change, we quantified EU incorporation together with the abundance and localisation of 18 proteins or post-translational modifications (PTMs) in the same cells, using iterative indirect immunofluorescence imaging (4i) (Figure 4A-B) (Gut et al., 2018). The antibody panel includes markers of active promoters (H3K4me3) and heterochromatin (H3K9me3), as well as RNA Pol II (POLR2A, also known as RPB1) and its phosphorylated forms: POLR2A-S5P, POLR2A-S2P, which are markers of transcription initiation (Glover-Cutter et al., 2009) and elongation (Peterlin and Price, 2006), respectively. We also measured nuclear speckles (SON) and speckle-associated proteins involved in splicing (SNRPB2), RNA stability (PABPN1), and export (ALYREF), as well as the nucleolus (NCL) and RNA Pol I (POLR1A).

**Figure 4:**
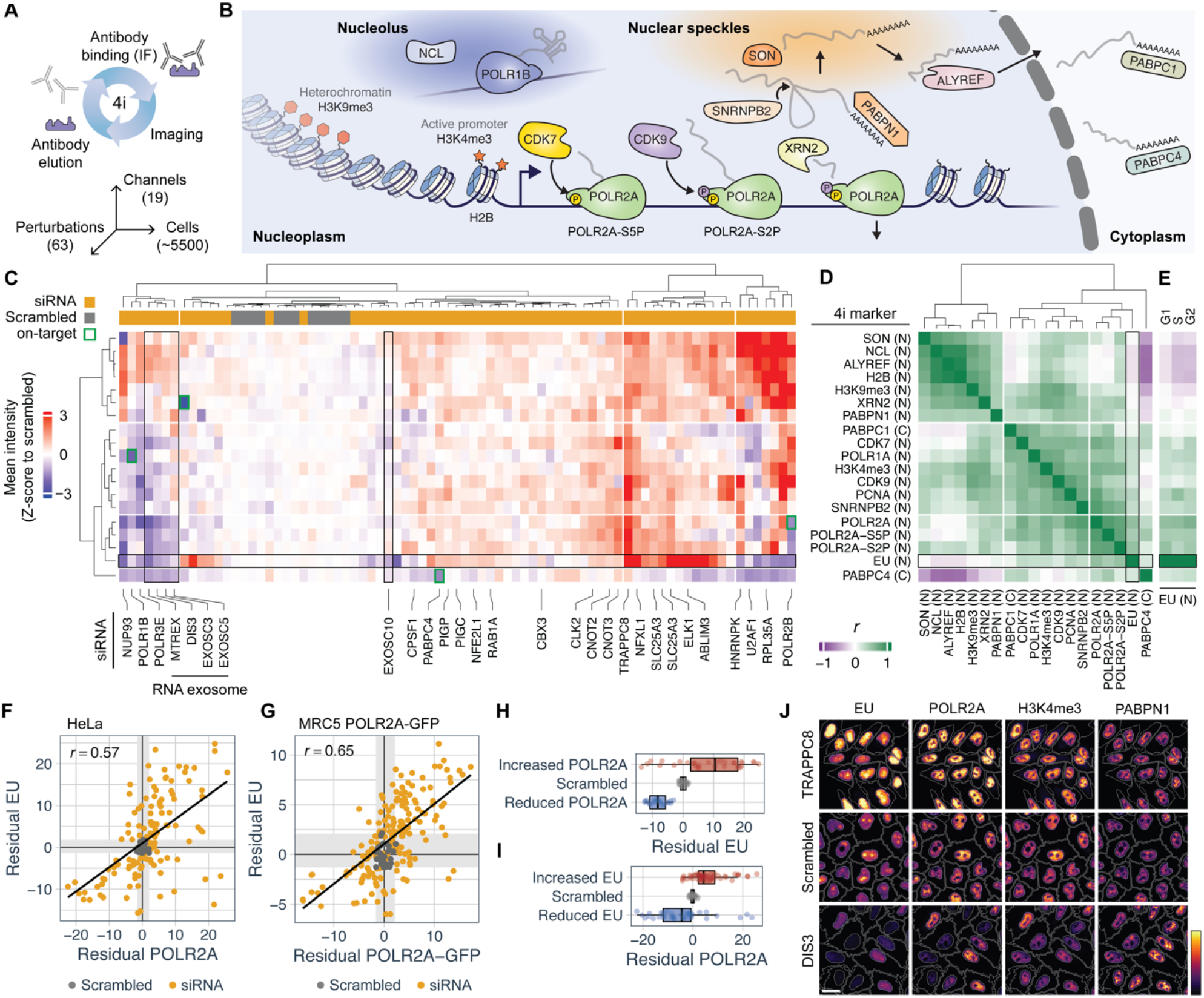
Highly multiplexed profiling of cellular phenotypes associated with altered RNA production. (A) 4i schematic (B) Diagram of proteins and PTMs measured by 4i and their roles in RNA metabolism. (C) Hierarchical clustering of 4i markers across 63 siRNA perturbations. Each marker standardised to nonperturbed (scrambled siRNA) cells using the robust z-score before averaging replicates. G1 cells only. On-target indicates that siRNAs directly target a gene in the antibody panel. Row labels in D. N indicates nuclear mean intensity while C indicates cytoplasmic mean intensity. (D) Pairwise correlations between perturbation-averaged 4i marker intensities (the rows of C). (E) Correlations of perturbation-averaged 4i marker intensities with EU by cell-cycle phase. (F) Residual mean POLR2A versus residual mean EU for each well in the 4i experiment. Pearson’s correlation for non-controls inset. Grey shaded boxes represent 1st/99th percentile of scrambled siRNA controls. (G) As in F, for MRC5 POLR2A-GFP cells with POLR2A quantified using GFP intensity, across the same 63 perturbations (n=3 per perturbation). (H) 4i experiment: residual mean EU for perturbations in which POLR2A is increased/decreased compared to scrambled siRNA controls (Benjamini-Hochberg adjusted p<0.05). (I) As in F, with POLR2A and EU reversed. (J) Example 4i images. See also Figure S8.

We selected 63 perturbations from the genome-wide screen, focusing on gene perturbations in which mRNA concentration homeostasis is perturbed (EXOSC3/5, NUP93) as well as other pathways enriched in the screen, including RNA processing (U2AF1, HNRNPK), transcription factors (NFXL1, ELK1), chromatin modification (SETD1A, CBX3), the endomembrane system (RAB1A, TRAPPC8) and GPI-anchor biosynthesis (PIGC, PIGP). After validating 4i-based quantification of protein abundance in single cells (STAR Methods, Figures S8A-B), we performed duplicate 4i experiments in which we analysed an average of 5,500 +/-2,500 cells (mean +/-s.d) per perturbation, together with over 80,000 control cells (Figures S8C-F).

To obtain an overview of how different proteins and PTM levels change in perturbations, we used hierarchical clustering of perturbation-averaged mean intensities, focusing on G1 cells (Figure 4C). This revealed diverse molecular phenotype profiles, which clustered into groups with characteristic EU incorporation. Notably, perturbation of four of the five RNA exosome components (DIS3, EXOSC3, EXOSC5, MTREX but not EXOSC10) clustered together, indicating a common cellular phenotype. This ‘DIS3 phenotype’ was characterised by reduction in abundance of Pol II and other components associated with active transcription (e.g. CDK7, H3K4me3), together with increases in mean ALYREF and H2B intensity. The DIS3 phenotype differed from other conditions with reduced EU incorporation, such as the disruption of components of pre-mRNA processing machinery, (e.g. HNRNPK, U2AF1, SNRPF), which showed greater reductions in cell viability, and more extreme changes to cell morphology.

To determine which markers show concordant changes across perturbations, we took perturbation-average intensity values of each marker and calculated pairwise correlations between them (Figure 4D). Clustering this correlation matrix revealed 4 groups of markers. Foremost, we observed that EU clustered with all three markers of RNA Pol II, revealing that changes in EU incorporation induced by genetic perturbations are most closely related to changes in RNA Pol II abundance (Figure 4E). A second group contained markers associated with “active transcription” such as CDK7, CDK9, H3K4me3, which also correlated well with EU. A third group containing H2B, H3K9me3, and ALYREF did not show a close relationship with EU and was anti-correlated with PABPC4, which showed surprisingly distinct changes in abundance compared to all other markers, even PABPC1 (Figure 4D). To further investigate the link between RNA Pol II abundance and EU incorporation, we corrected EU and POLR2A intensities for cell-cycle and cell-size effects (STAR Methods). Residual EU and residual POLR2A were well correlated across perturbations (r_EU, POLR2A_ = 0.57, Figure 4F, S8H-I), and significant changes to one were associated with changes to the other (Figure 4H-I). To validate this finding, we made use of MRC5 cells in which both copies of POLR2A are tagged with GFP (Steurer et al., 2018). Direct imaging of POLR2A-GFP and EU across the same set of 63 perturbations revealed a similar positive correlation (*r*_EU, POLR2A-GFP_ = 0.65, Figures 4G, S8J-K), confirming that RNA Pol II abundance typically changes together with transcriptional activity in perturbations.

To visualise the multidimensional character of the measured phenotypes in single cells, we embedded all 434,575 cells in a 2D Uniform Manifold Approximation and Projection (UMAP) (McInnes et al., 2018), using intensity and texture features derived from 4i together with morphology and cell crowding features (Figure 5A-B). Perturbations typically localised to specific regions of the UMAP, however almost all contained a subpopulation that cannot be readily distinguished from scrambled siRNA controls (Figure 5C, S9A), likely corresponding to non-perturbed cells. Although EU was omitted when constructing the UMAP, other features in the data led to a non-random pattern of EU intensity, which is distinct from both cell cycle and nuclear area (Figures 5B, S9B). We observed two major axes of variability in the main body of the UMAP: from bottom to top, nuclei become larger and cell cycle progresses, while from left to right, POLR2A intensity increases together with EU intensity, and other “active transcription” markers. Perturbations with increased EU incorporation, such as TRAPPC8 and NFXL1, typically occupy this region with increased intensity of active transcription markers (Figure 5C). As expected, targeting one of the proteins in the antibody panel whose intensities are used to create the UMAP (POLR1A, PABPC4, XRN2, POLR2B), led to perturbations in which cells localised away from the main group of cells (Figure 5A). Notably, RNA exosome perturbations localised to two distinct clusters (Figure 5C), with DIS3 and MTREX primarily occupying the lower cluster, and EXOSC3 and EXOSC5 occupying both. The EXOSC3/5-specific cluster is distinguished from the other by increased PABPN1 levels (nuclear poly(A) binding protein) and a less pronounced reduction in POLR2A (Figure S9B-C), which may also be related to the increases in nuclear mRNA that we saw using poly(A) FISH for EXOSC3/5 but not MTREX/DIS3 (Figure 3A, S6G).

**Figure 5:**
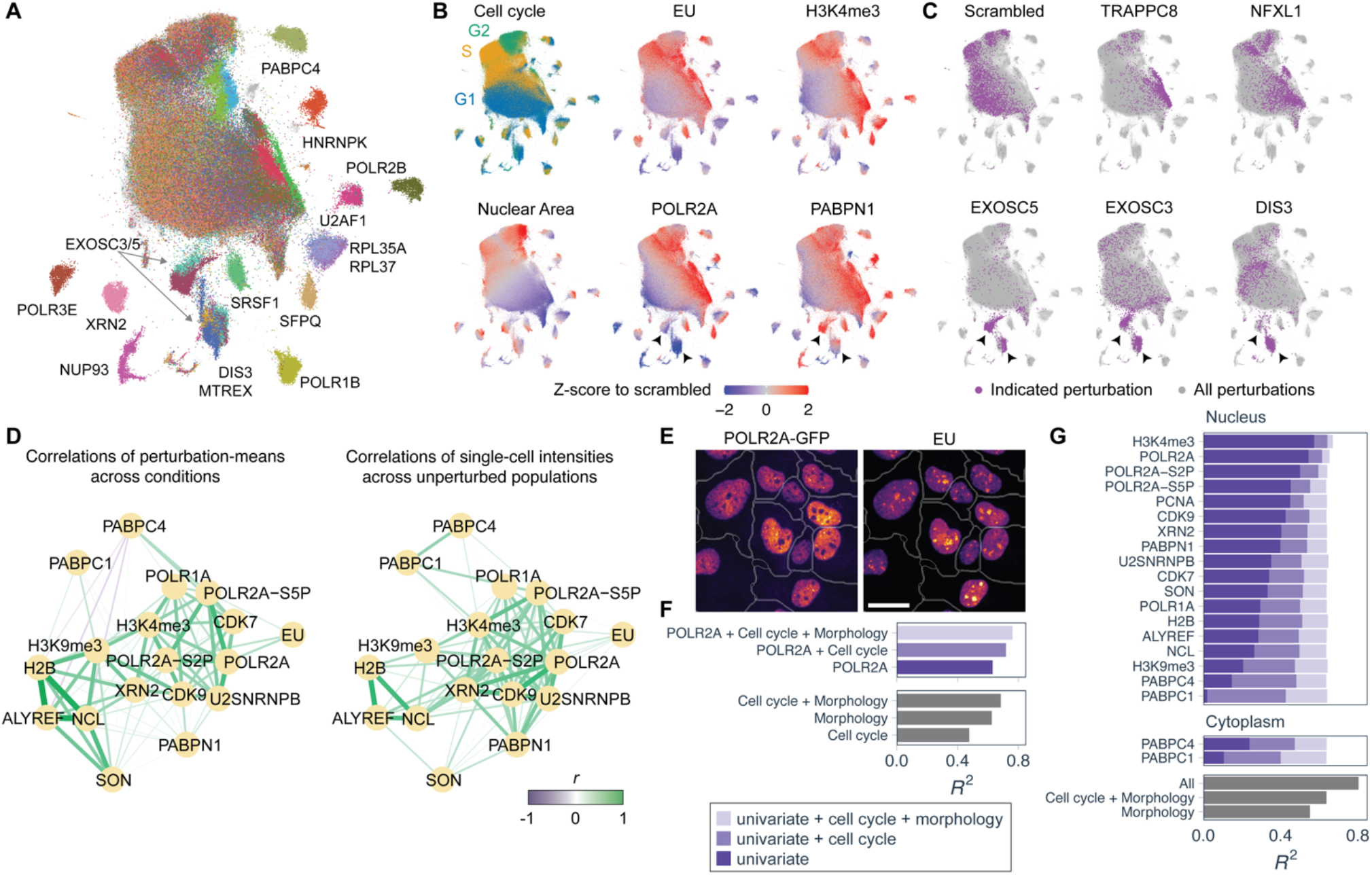
Coordination of transcription machinery with RNA production at the single-cell level. (A) UMAP of 434,575 HeLa cells from 4i experiment. Selected perturbations coloured, with remaining cells grey. Labels indicate predominant perturbations in each cluster. (B) UMAP coloured by cell cycle or nuclear mean intensity of indicated marker. (C) Distribution of cells on UMAP for selected perturbations. (D) Pairwise correlations between 4i markers represented as a network, either calculated across perturbations or across unperturbed cell populations. G1 cells only. (E) GFP fluorescence and EU metabolic labelling in MRC5 POLR2A-GFP cells. (F) Coefficient of determination (*R*^2^) of linear regression predicting sum nuclear EU in MRC5 POLR2A-GFP cells at the single-cell level. Predictors indicated on axis. (G) *R*^2^ of linear regression predicting sum nuclear EU in HeLa cells, with 4i markers as predictors, as indicated. Univariate models have a single predictor, while ‘+ Cell cycle’, ‘+ Morphology’, and ‘All’ indicate the additional predictors included (STAR Methods). See also Figures S9-S10.

We noticed that the distribution of unperturbed (scrambled siRNA) cells on the UMAP extended into regions typically occupied by genetically perturbed cells (Figure S9A). This indicates that there is substantial cell-to-cell variability in levels of measured proteins and PTMs even within unperturbed cell populations. To understand how this heterogeneity is related to that of RNA production rates, we calculated correlations between mean intensities of 4i markers and EU at the single-cell level in unperturbed controls. Several markers, including POLR2A, CDK7, and H3K4me3, were positively correlated with EU (Figure S10A), indicating that cellular levels of RNA Pol II and other active transcription markers in *unperturbed* populations increase with RNA production rates. Interestingly, 4i markers that covary with EU in unperturbed populations are typically those that also showed concordant changes with EU when the latter is perturbed. That is, correlations 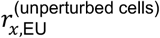 and 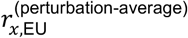 are themselves positively correlated across 4i markers (*r* = 0.65, p <10^-7^, Figure S10B-C). Moreover, network representations of these pairwise correlations calculated at the perturbation-scale and the single-cell scale were highly similar (Figure 5D). At both levels, we observed a module of “active transcription” markers including CDK7, H3K4me3, POLR2A, and SNRPB2 that are mutually coordinated and positively related to EU incorporation, and a second set of factors including H2B, ALYREF, and NCL are also mutually coordinated but are unrelated to EU incorporation.

To confirm the close relationship of Pol II abundance and RNA production in unperturbed cells, we performed EU metabolic labelling with cell volume and cell cycle measurement in unperturbed MRC5 POLR2A-GFP cells (Figure S10D). Strikingly, POLR2A and EU were highly correlated at the singlecell level, and POLR2A-GFP showed a similar relationship as EU with both cell volume and cell cycle (Figures 5E, S10D-G). Furthermore, using linear regression, we found that POLR2A-GFP intensity alone explained a similar fraction of variance in EU incorporation as a ‘cell size’ model with cell volume and protein content as predictors (mean *R*^2^ = 0.61 for both models, Figure 5F). In the HeLa 4i dataset, regression models predicting EU at the single-cell level from each 4i marker individually had *R*^2^ ranging from 0.02 for nuclear PABPC1 to 0.55 and 0.58 for POLR2A and H3K4me3, respectively, with latter values similar to those of a morphology-only model (*R*^2^ = 0.55, Figure 5G). When combined with morphology and cell cycle information, POLR2A and H3K4me3 were the only two markers which increased *R*^2^ compared to models with morphology and cell cycle features alone. Overall, this analysis reveals that cells coordinate the cellular abundance of Pol II and other markers of active transcription with RNA production – both in unperturbed cells and when RNA production is perturbed.

### Repression of transcription by increased nuclear RNA abundance

Our results thus far reveal that mRNA concentrations are generally maintained despite large changes to RNA production rates. However, we identified knockdown of the nuclear RNA exosome as one of few conditions in which RNA concentration homeostasis is disrupted (see Figure 3E-F). In this case, nuclear mRNA accumulation is associated with reduced EU incorporation and reduced levels of Pol II and several other markers of active transcription that are normally coordinated with RNA production rates in single cells (see Figure 4C). From our genome-wide screen, we found that perturbations that would lead to nuclear mRNA accumulation (for example genes involved in mRNA processing, nuclear RNA export, or nuclear RNA degradation) are also associated with reduced RNA production. This suggests that nuclear mRNA abundance may negatively feed-back to regulate transcription. Such a model is appealing because it could potentially underlie the coordination of transcription with cell size: for example, because nuclear and cell size are coupled (Cantwell and Nurse, 2019), transcription rates would scale with nuclear size (and therefore cell size) to maintain a constant nuclear mRNA concentration. To further pursue this hypothesis, we turned to the auxin-inducible degradation (AID) system (Nishimura et al., 2009) (Figure 6A) to examine the effects of rapid disruption of nuclear RNA degradation.

**Figure 6:**
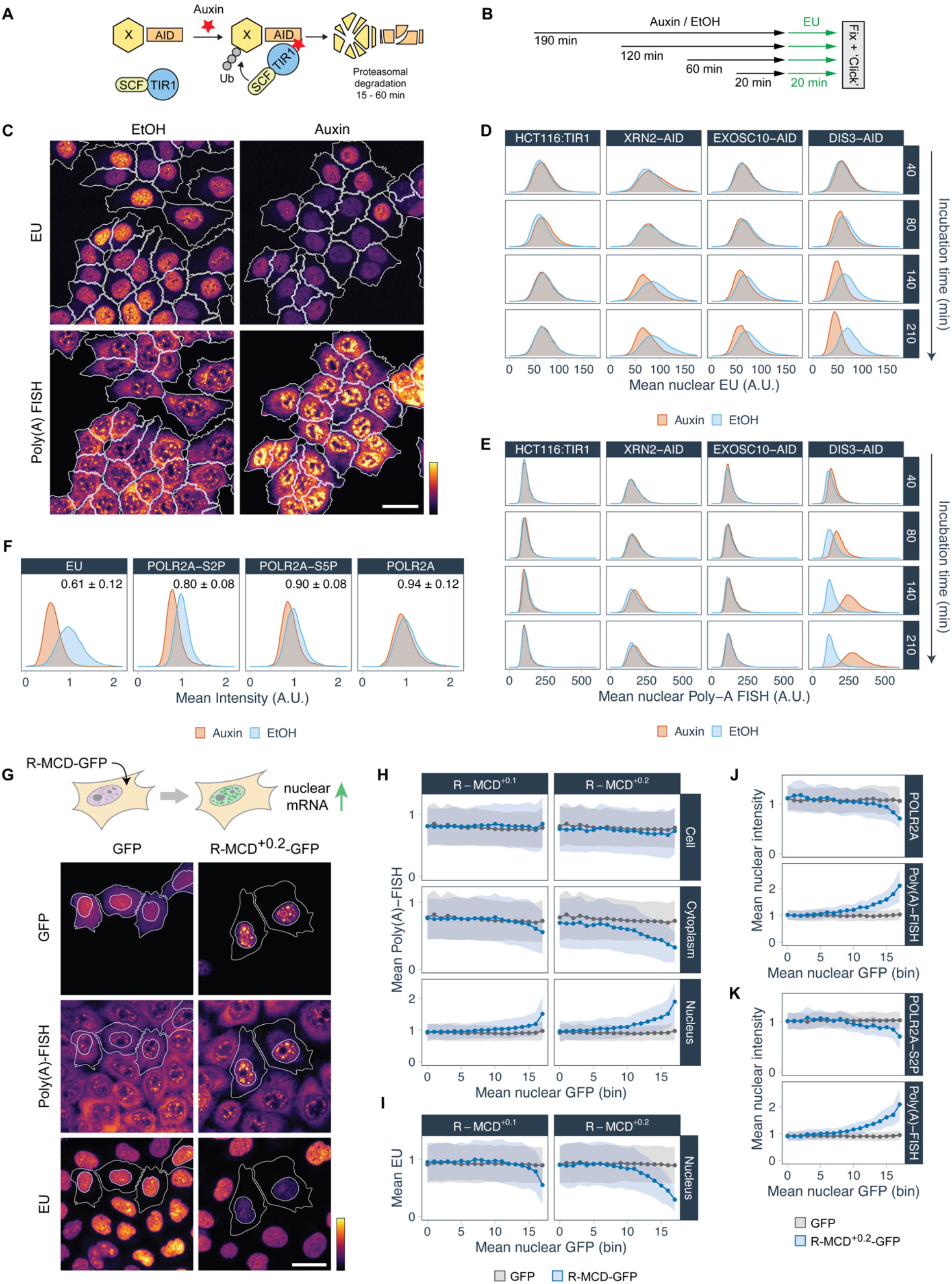
Repression of transcription upon increased nuclear mRNA abundance. (A) Auxin-inducible degradation (AID). (B) AID time-course experiment. (C) EU incorporation and poly(A) FISH in DIS3-AID cells after 140 min Auxin or EtOH. (D) Mean nuclear EU intensity distributions in parental HCT116:TIR1 and AID cells in time-course experiment. (E) As in D, for nuclear poly(A) FISH. (F) Mean nuclear intensity distributions of EU and POLR2A (immunofluorescence), in DIS3-AID cells treated with Auxin or EtOH for 3.5h. Inset: population median for Auxin relative to EtOH controls (mean ± s.d., n=6 (IF) or n=18 (EU) wells). (G) Poly(A) FISH and EU incorporation in cells expressing R-MCD^+0.2^-GFP or GFP-control. GFP-expressing cells outlined. (H) Mean poly(A) FISH intensity for the cell, cytoplasm or nucleus, as a function of binned nuclear GFP intensity (GFP: n=23,927; >715 per bin; R-MCD^+0.^-GFP: n=14,179; >388 per bin; R-MCD^+0.2^-GFP: n=6,296; >290 per bin). Points and shaded regions show mean ± s.d. (I) As in H, for mean nuclear EU (same cells as H) (J) As in H, for mean nuclear POLR2A (GFP: n=6,442; >175 per bin; R-MCD^+0.2^-GFP: n=1,225; >51 per bin) (K) As in H, for mean nuclear POLR2A-S2P (GFP: n=5,167; >123 per bin; R-MCD^+0.2^-GFP: n=1,346; >61 per bin). See also Figure S11.

HCT116 cells that contain AID-tagged versions of either DIS3 or EXOSC10 (catalytic exosome subunits) or XRN2 (a nuclear 5’ to 3’ exonuclease involved in transcription termination) allow depletion of target proteins within an hour (Davidson et al., 2019; Eaton et al., 2018). Using these cells, we performed time-course experiments treating with auxin or EtOH (vehicle) for 20-190 min before pulse-labelling with EU for the final 20 minutes before fixation (Figure 6B). Beginning at 60-80 minutes after auxin addition, we saw reduced EU incorporation in DIS3-AID, EXOSC10-AID and XRN2-AID cells compared to EtOH-treated controls (Figure 6C-D, S11A-B). The effect was strongest in DIS3-AID cells, with EU incorporation reduced by 35 ± 5 % (mean ± s.d., *p* < 10^-9^) after 3.5h. This is likely an underestimate of the actual decrease in RNA synthesis because acute perturbation of RNA degradation is expected to stabilise nascent RNA, and therefore increase rather than decrease EU intensity. Long-term depletion of exosome core components EXOSC3 and EXOSC5 by siRNA results in accumulation of nuclear mRNA (Figures 3A,3E). We therefore also performed time-course experiments in these AID cells using poly(A) FISH to visualise mRNA. XRN2-AID and EXOSC10-AID did not show increased poly(A) FISH intensity – consistent with the RNA that accumulates in these cases being non-polyadenylated (Eaton et al., 2018). In DIS3-AID however, we observed strong auxin-dependent accumulation of nuclear mRNA, which occurred over the same time-scale as the reduction in RNA synthesis (Figure 6C,E). Moreover, transcriptional reduction and mRNA accumulation occurred in the same cells (Figure S11C). This differs from long-term depletion of DIS3 by siRNA, which does not result in nuclear mRNA accumulation (see Figure S6G) due to partial functional redundancy with EXOSC10 (Fan et al., 2018). The effects of such redundancy are substantially minimised using rapid auxin-mediated DIS3-depletion (Davidson et al., 2019).

DIS3 performs post-transcriptional nuclear RNA degradation, suggesting that accumulation of nuclear RNA may be the primary event that occurs upon DIS3 depletion, with transcriptional reduction occurring as a consequence of this – perhaps as a result of the hypothesized feedback from mRNA on transcription. To determine whether rapid DIS3 depletion is associated with changes in Pol II, we measured POLR2A, POLR2A-S5P, and POLR2A-S2P by immunofluorescence (Figure 6F, S12D). After 3.5h of auxin treatment, we observed a ~40% reduction in EU incorporation, which was accompanied by a ~20% reduction in POLR2A-S2P, a ~10% reduction in POLR2A-S5P and a ~6% reduction in total POLR2A. This reveals that DIS3 depletion initially leads to reduced Pol II *activity*, with reduction in Pol II *abundance* occurring over a longer timescale (Figure 4C).

To test the hypothesis of a negative feedback of nuclear RNA concentration on transcription in an orthogonal manner, we made use of artificial arginine-enriched mixed-charge domain (R-MCD) proteins, which were recently found to drive nuclear mRNA retention (Greig et al., 2020). When expressed in cells, positively charged R-MCD^+0.1^-mGFP and R-MCD^+0.2^-mGFP localised to nuclear speckles, leading to dose- and charge-dependent nuclear accumulation and cytoplasmic depletion of mRNA (Figure 6G-H, S11E-F), in line with previous data. Total cellular mRNA abundance was not affected. In agreement with our hypothesis, this nuclear mRNA retention was accompanied by reduced EU incorporation (Figure 6I). Moreover, EU reduction and nuclear mRNA accumulation showed similar quantitative dependence on GFP intensity. In addition, nuclear mRNA retention driven by R-MCD^+0.2^-eGFP expression was associated with reductions in both POLR2A and POLR2A-S2P – most prominently at the highest GFP intensities (Figures 6J-K, S11G-H).

Elevating nuclear mRNA abundance by either acutely disrupting nuclear RNA degradation or forcing nuclear mRNA retention therefore leads to reduced RNA synthesis. Because exosome depletion initially affects transcriptional activity, with Pol II abundance changes occurring over longer timescales, this suggests that Pol II abundance is *dictated by* transcriptional activity, rather than determining it – in contrast to the limiting factor model. Furthermore, since R-MCD expression results in cytoplasmic depletion of mRNA, *nuclear* rather than *cytoplasmic* RNA abundance is relevant for the reduction of RNA synthesis.

### A minimal mathematical model of transcription with negative feedback from mRNA

To explore how nuclear mRNA could regulate transcription and ultimately coordinate RNA Pol II levels with transcription rates and cell size, we constructed a simplified mathematical model incorporating Pol II-mediated transcription and nuclear mRNA.

It is well known that inhibiting POLR2A with α-amanitin (Lee et al., 2002; Mitsui and Sharp, 1999; Nguyen et al., 1996), as well as blocking transcription initiation with triptolide (Alekseev et al., 2017;Bensaude, 2011; Steurer et al., 2018) both lead to Pol II degradation. Yet when these compounds are combined with proteasome-inhibition, Pol II detaches from chromatin but remains stable (Steurer et al., 2018). This is consistent with a model in which Pol II is targeted for degradation primarily when it is not chromatin-associated, which could provide a generic mechanism to coordinate Pol II abundance with transcriptional activity, as we observed experimentally. To formulate this mathematically, we adapted a kinetic model of Pol II transcription derived from fluorescence-recovery after photobleaching (FRAP) (Steurer et al., 2018), adding the hypothesis that only ‘unbound’ Pol II can be degraded (Figure 7A, S12A). We then measured POLR2A and POLR2A-S2P levels using immunofluorescence, after treating cells with triptolide or CDK9 inhibitor AZD4573 (which prevents Ser2-phosphorylation of Pol II CTD) (Cidado et al., 2020). Identifying the elongating state as POLR2A-S2P (Heidemann et al., 2013), we simulated triptolide and AZD4573 treatments and optimised a single free parameter (Pol II synthesis rate) to fit the data. The model quantitatively reproduced triptolide-induced loss of POLR2A, and AZD4573-induced reductions of POLR2A-S2P and POLR2A (Figures 7B). In contrast, an alternative model in which all Pol II species were subject to degradation did not show POLR2A abundance changes (Figure 7B, S12B).

**Figure 7:**
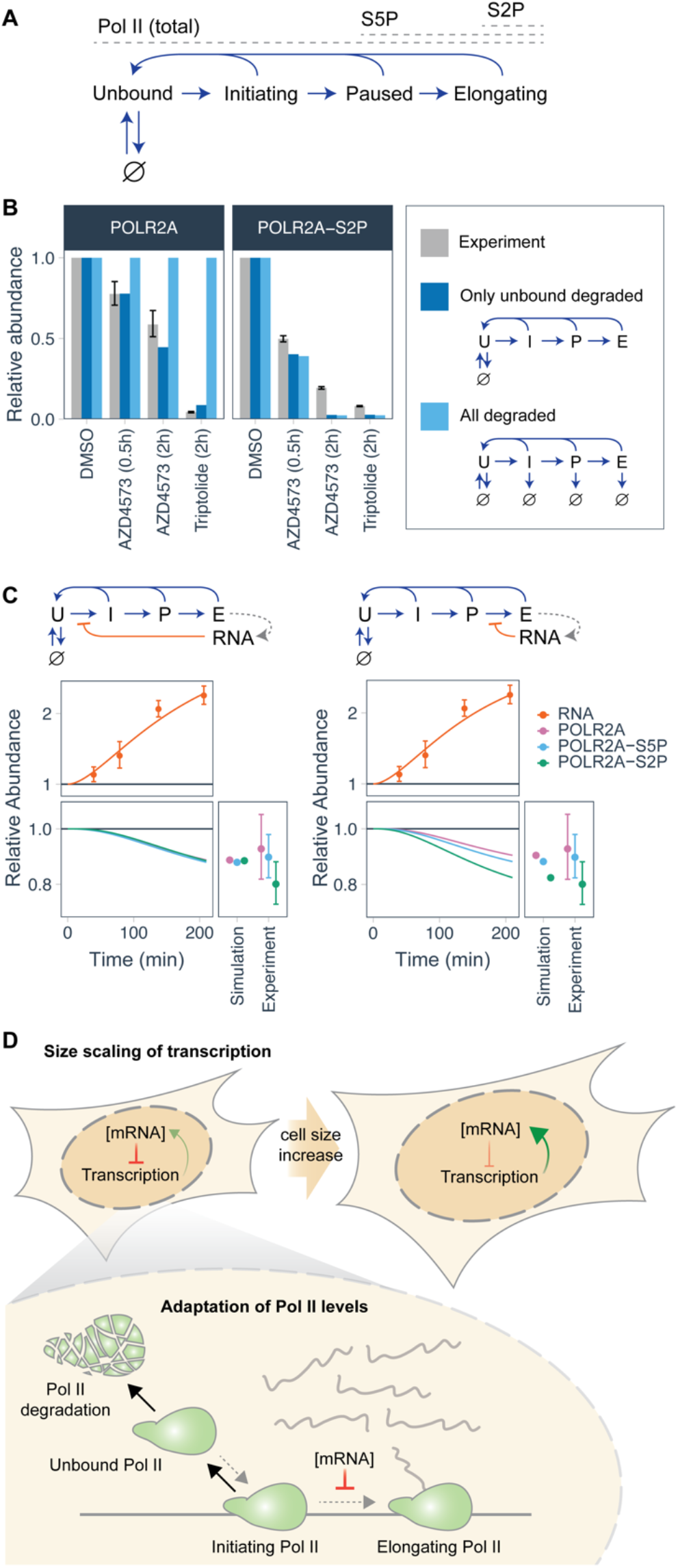
Mathematical model of transcription with negative feedback from mRNA. (A) Schematic of Pol II states and transitions in the model. Arrows to/from ∅ indicate Pol II degradation/synthesis. (B) Nuclear POLR2A and POLR2A-S2P levels relative to DMSO controls, measured by immunofluorescence upon chemical perturbation of transcription. Best-fit models shown for comparison. (C) Simulation of DIS3-AID depletion experiment for models including negative feedback from mRNA, as indicated in schematics above. Best-fit models shown with poly(A) FISH data from Figure 6E, and Pol II immunofluorescence data from Figure 6F. All error bars represent mean ± s.d. for all pairwise comparisons of treated and control wells. (D) Schematic of the proposed mechanism. See also Figure S12.

We next investigated how mRNA abundance could affect Pol II-dependent transcription by considering the different points at which the feedback may act: either by stimulating Pol II degradation or inhibiting Pol II initiation, pausing, or pause release. Upon simulation of auxin-induced depletion of DIS3, all models reproduced the experimentally observed increase in mRNA abundance (Figure 7C, S12C), however the predictions for Pol II were slightly different: models where mRNA activates Pol II degradation or represses transcription initiation or pausing predict simultaneous loss of phosphorylated and total POLR2A. Conversely, when mRNA represses the transition from pausing to elongation, the model predicts greater loss of POLR2A-S2P than POLR2A-S5P and total POLR2A (Figure 7C). Of the models considered, the latter is the most similar to our experimental observations, suggesting that feedback of nuclear mRNA on transcription may act through inhibition of transitions from paused to elongating Pol II.

In summary, the model reveals that preferential degradation of unbound Pol II is sufficient to make overall Pol II abundance dependent on transcriptional activity. When combined with mRNA-based feedback, transcriptional activity and ultimately Pol II levels are then also sensitive to changes in nuclear mRNA degradation and export rates, through their effects on nuclear mRNA concentration. This is consistent with the effects we see across perturbations wherein transcription rates and Pol II levels respond to perturbation of RNA processing downstream of transcription. Because the model is formulated in terms of concentrations, steady states correspond to mRNA synthesis rates and Pol II abundance that scale with cell volume (Figure 7D). Negative feedback from mRNA makes steady-state concentrations of Pol II and nuclear mRNA less sensitive to changes in parameters such as Pol II synthesis and degradation (Figure S12D-E), and also allows faster restoration of Pol II and mRNA levels upon perturbation of cell volume (Figure S12F-G) – increasing the robustness of mRNA concentration homeostasis in fluctuating environments.

## DISCUSSION

Here we reveal the remarkable robustness of mRNA concentration homeostasis in human cells, to changes in either cell volume or RNA production rates. Scaling of transcript abundance with cell volume is underpinned by increasing transcription rates with cell size, a process that we reveal involves coordinated control of Pol II abundance and other hallmarks of RNA production including H3K4me3, CDK7 and SNRPB2. Acute perturbation experiments however show that this coordination is not because Pol II abundance directly determines transcription rates, but because Pol II abundance is adapted to transcriptional activity. Rather, our experiments and modelling point to nuclear mRNA concentration as being the quantity under strict homeostatic regulation and reveal that this negatively impinges on transcriptional activity to ultimately determine the abundance of the transcription machinery. This mechanism has a strong parallel with classic ‘feedback inhibition’ – employed pervasively throughout metabolic networks to coordinate activities of biosynthetic enzymes with cellular requirements (Pardee and Reddy, 2003). Feedback inhibition through allosteric effects is more robust than control of enzyme abundance (Sander et al., 2019), which is analogous to mRNA acting primarily on transcriptional activity rather than on Pol II abundance, as indicated by fitting the model to the DIS3-AID experiments.

Such a mechanism in mammalian cells contrasts the buffering model proposed for budding yeast, based on the finding that deletion of cytoplasmic RNA decay factors, especially Xrn1, results in reduced transcription (Haimovich et al., 2013; Sun et al., 2013). While we also find that cytoplasmic RNA concentration is buffered upon genetic perturbation of transcription, multiple observations indicate a crucial role for the nucleus in this process: First, the genome-wide screen reveals that perturbation of nuclear rather that cytoplasmic RNA degradation factors feed back on transcription (Figures 2G, S5A). Second, only upon perturbation of nuclear RNA degradation and export is homeostasis of mRNA concentration broken (Figure 3E). Third, nuclear RNA retention experiments indicate that nuclear rather than cytoplasmic RNA is relevant for the homeostatic feedback on transcription (Figure 6H-I). This is also consistent with reports that accelerated *cytoplasmic* mRNA degradation does not result in a compensatory increase in transcription in human cells (Abernathy et al., 2015; Gilbertson et al., 2018). Interestingly, recent work in fission yeast also suggests that nuclear size rather than cell size may be the quantitative determinant of mRNA size-scaling (Sun et al., 2020).

The detailed mechanism by which mRNA concentration regulates RNA production remains to be characterized. A direct effect is possible. Indeed, in vitro transcription is inhibited by exogeneous RNA. This has been proposed to occur by RNA directly interfering with Pol II binding to the DNA template (Pai et al., 2014) and could involve interference with the phase separation of transcriptional condensates on chromatin (Henninger et al., 2021; Portz and Shorter, 2020). An indirect effect is also possible: with mRNA modulating the activity or localisation of a limiting transcriptional regulator. Nuclear RNA concentration has been proposed to regulate condensation of nuclear RNA-binding proteins (Maharana et al., 2018) and, interestingly, nuclear speckles are enriched for transcription elongation factors, rather than transcription initiation factors (Galganski et al., 2017). Increased mRNA levels in nuclear speckles, for example as observed upon R-MCD overexpression, could result in retention of elongation factors and thereby suppress efficient Pol II elongation.

The minimal model developed here explains several experimental observations and *buffers* mRNA concentration. However, in its current form, it does not achieve robust ‘set-point’ homeostasis (Briat et al., 2016; Reed et al., 2017). Further elaboration of the model requires more details on the molecular nature of the transcriptional feedback, and on how mRNA degradation and export rates are affected by mRNA abundance. For example, nuclear mRNA export in human cells appears to depend on ongoing transcription (Tokunaga et al., 2006), which could also contribute to the maintenance of nuclear mRNA levels that we observed in perturbations with strong transcriptional repression, such as POLR2B knockdown. In addition, work in yeast has revealed that blocking nuclear export can lead to increased decay of newly synthesized mRNA (Tudek et al., 2018). Such mechanisms could both play key roles in set-point homeostasis of nuclear mRNA concentration, but it is currently unclear how they are inter-connected. The rich perturbation datasets and simple model that emerge from our work, linking activity and abundance of the transcription machinery to nuclear mRNA concentration, provide a starting point to explore links between these global cellular controls of mRNA metabolism and how they relate to the volume of mammalian cells.

## ACKNOWLEDGMENTS

We thank Larisa Venkova and Matthieu Piel for FXm training and resources; Merve Avar, Daniel Heinzer, Marc Emmenegger and Adriano Aguzzi for printing siRNA libraries; David Spector for the anti-SNRPB2 antibody; and all members of the Pelkmans Lab for discussions and manuscript comments. AID cells were kindly provided by Steven West. POLR2A-GFP cells were kindly provided by Jurgen Marteijn. S.B. was supported by an EMBO long-term fellowship (ALTF1175-2016), and an HFSP long-term fellowship (LT000238/2017-L). L.P. is supported by the European Research Council (ERC-2019-AdG-885579), the Swiss National Science Foundation (SNSF grant 310030_192622), and the University of Zurich.

## AUTHOR CONTRIBUTIONS

Conceptualization, S.B. and L.P.; Methodology, S.B.; Investigation, S.B. and M.M.; Formal Analysis, S.B. and M.M.; Writing–Original Draft, S.B.; Writing – Review & Editing, S.B., M.M. and L.P.; Funding Acquisition, S.B. and L.P.

## STAR METHODS

### RESOURCE AVAILABILITY

#### Lead contact

Further information and requests for resources and reagents should be directed to and will be fulfilled by the Lead Contact, Lucas Pelkmans.

#### Materials Availability

This study did not generate new unique reagents.

#### Data and Code Availability

Images, single-cell level features and summary data for the genome-wide screen have been deposited at the Image Data Resource (IDR) under accession number idr00093, and are described in an accompanying manuscript (Müller et al., 2021). Single-cell-level features and summaries for other genetic perturbation experiments are provided at the locations listed in Key Resources. Raw image datasets for other experiments are available upon reasonable request.

Image analysis was performed using TissueMAPS, an open-source project for high-throughput image analysis developed in our group. All analysis modules, including code written for this paper, is packaged together with the main repository at https://github.com/pelkmanslab/TissueMAPS. An example analysis pipeline description with module files containing parameter settings for the genomewide screen is provided at IDR (idr00093). This example is a typical TissueMAPS workflow, however the exact modules used and parameter values differ depending on experiments. These can be provided on request. Code for calculating population-context features was written in python based on previous MATLAB code from our group (Snijder et al., 2012) and is available at https://github.com/scottberry/popcon. R code for the analysis of single-cell data derived from images was developed on a per-experiment basis and is available on request.

### EXPERIMENTAL MODEL AND SUBJECT DETAILS

#### Cell lines and culture conditions

HeLa Kyoto (female) cell populations were derived from a single-cell clone and were tested for identity by karyotyping (Battich et al., 2015). HeLa cells were cultured in high glucose DMEM supplemented with 10% fetal bovine serum (FBS) and 1% GlutaMAX. Cells with low passage number (2-6) were used for all experiments.

184A1 (human female breast epithelial) cell populations were derived from a single-cell clone, and were used at low passage number (2-6) for all experiments. 184A1 cells were cultured in DMEM/F12 media supplemented with 5% horse serum, 20ng/ml epidermal growth factor, 10μg/ml insulin, 0.5μg/ml hydrocortisone, 10ng/ml cholera toxin.

Keratinocytes were donated by a healthy 2.5-year-old male (Biedermann et al., 2010), isolated and kindly provided by E. Reichmann and L. Pontiggia (University of Zurich, Zurich). Keratinocytes were cultured in CnT-57 medium (CELLnTEC) supplemented at 1:100 (v/v) with Penicillin/Streptomycin. For propagation and experiments, plastic cell culture plates were coated with rat-tail collagen I, according to manufacturer’s instructions. Experimental plates were prepared 5 days after thawing cells (passage 1).

Parental MRC-5 and derived POLR2A-GFP (RPB1-eGFP) cells (Steurer et al., 2018) were cultured in DMEM/F12 supplemented with 10% FBS and 1:100 (v/v) Penicillin/Streptomycin.

Parental HCT116:TIR1 and derived EXOSC10-AID, DIS3-AID, XRN2-AID cells (Davidson et al., 2019; Eaton et al., 2018) were cultured in DMEM supplemented with 10% FCS and 1:100 (v/v) Penicillin/Streptomycin. All cells were tested for absence of mycoplasma before use and grown at 37°C, 95% humidity and 5% CO_2_.

Unless otherwise specified, cells were grown and imaged in uncoated Greiner μClear plastic-bottom 96- or 384-well plates.

### METHOD DETAILS

#### siRNA transfection

Transfection with siRNA was performed as previously described (Berchtold et al., 2018). Briefly, 900 HeLa or 1500 MRC5 POLR2A-GFP cells were plated per well in 384-well plates for reverse transfection onto a mixture of pooled siRNAs (5 nM final concentration) and Lipofectamine RNAiMAX (0.08μl per well in OptiMEM) according to manufacturer’s specifications. Cells were subsequently grown for 72 hours at 37°C in a final volume of 50μL growth media, to establish efficient knockdown of the targeted genes. For the genome-wide and secondary screens, siRNAs were dispensed in a much lower volume using an acoustic dispenser, and assays were performed in a final volume of 40μL. A detailed protocol was described previously (Müller et al., 2021).

The genome-wide screen was performed using a pool of 3 siRNAs (Ambion) targeting each gene. 21538 siRNA pools were assayed across 83 384-well plates, as previously described (Müller et al., 2021). The secondary library (EU metabolic labelling, poly(A) FISH, RNA StrandBrite) employed the same siRNA pools for 436 gene perturbations. Secondary assays were performed in duplicate. The tertiary library (4i, smFISH) consisted of a subset of 63 perturbations from the secondary library. Gene lists and other metadata are provided together with the experimental results at the locations listed in Key Resources.

#### Plasmid transfection

2700 HeLa cells were plated in each well of a 384-well plate. 12 hours after seeding, 50ng plasmid was transfected per well using Lipofectamine 2000, according to manufacturer’s protocol. Cells were then incubated for 2 hours at 37°C and then washed three times into fresh media to remove unbound transfection reagent/plasmid complexes. Cells were subsequently grown for 48 hours at 37°C before fixation, staining and imaging.

#### Chemical treatments

RNA polymerase I inhibitor, CX-5461 (Drygin et al., 2011) was dissolved in 5mN HCl at a concentration of 5mM and used at 2μM. XPB (TFIIH) inhibitor triptolide (Titov et al., 2011) was dissolved in DMSO at a concentration of 10mM and used at 2μM. CDK9 inhibitor AZD4573 (Cidado et al., 2020) was dissolved in DMSO at a concentration of 10mM and used at 0.1μM. CDK inhibitor roscovitine was dissolved in DMSO to a final concentration of 10mM and used at 20μM. Auxin was dissolved in ethanol at 200mM and used at 0.5mM. Durations of chemical treatments vary, and are noted throughout the text and figure legends.

#### Image acquisition

Unless otherwise specified, all imaging was performed an automated spinning-disk microscope (CellVoyager 7000, Yokogawa), which is equipped with four excitation lasers (405, 488, 568, 647nm) and two Neo sCMOS cameras (Andor). For cell volume measurement and smFISH experiments, a 40X/NA0.95 air objective was used. For other experiments such as the genome-wide screen, secondary screen, poly(A) FISH, RNA StrandBrite, 4i, and R-MCD-GFP experiments, a 20X/NA0.75 objective was used. Certain examples images (e.g. Figure 1D) were acquired with a 60X/NA1.27 objective. With the exceptions of cell volume experiments and 3D smFISH, images were maximum-projected during acquisition. Images presented in the same figure for the same stain were always identically rescaled.

#### DNA and protein staining

Nuclear DNA was stained using 4’,6-diamidino-2-phenylindole, dihydrochloride (DAPI) for 5-10 minutes at a final concentration of 0.4μg/mL in phosphate buffered saline (PBS). Total protein was stained using Alexa Fluor 488 NHS Ester or Alexa Fluor 647 NHS Ester (succinimidyl ester) for 10 minutes at a final concentration of 0.2μg/mL in 50mM carbonate-bicarbonate buffer pH 9.2.

#### High-throughput cell volume measurement

To measure cell volume in fixed cell populations, we developed a novel approach which involves attaching fluorescent beads to the upper surface of cells, and also the slide surface (Figure S1B). We then determine the three-dimensional positions of these beads using spinning-disk confocal microscopy and use these to generate a computational reconstruction of the cell.

All staining and washing steps were performed either manually, or on a semi-automated liquid handling platform (BioTek) using the following method: cells were fixed with 4% PFA for 30 min and washed four times with phosphate-buffered saline (PBS). To avoid beads entering inside cells, we performed biotinylation and bead attachment before cell permeabilization. To biotinylate cell surface, EZ-Link Sulfo-NHS-LC-Biotin was freshly dissolved in PBS and added to cells at a final concentration of 0.25mg/mL for 5 min. Cells were then washed four times with PBS. To prepare beads for attachment, we diluted 40nm-diameter streptavidin-coated fluorescent beads (Fluospheres) into a buffer containing 0.5x PBS and 0.01% Triton X-100, to a concentration of 0.005% solids (1/200 from 1% stock). Beads were then dispersed by sonication in 1mL aliquots for 3 x 30 s in a Bioruptor waterbath sonicator (Agilent), and were then added to cells by adding an equal volume of bead suspension to the residual PBS in multiwell plates (final bead concentration 0.0025% solids). Bead suspensions were mixed in the wells by pipetting up and down, or brief vigorous shaking of plates and were then incubated at room temperature for 10 min before washing off unbound beads with PBS. Cells were then permeabilised before proceeding with other stains.

After additional staining, beads were imaged as a separate acquisition (excitation laser: 568nm, emission filter 590/20 nm) using a 40X/NA0.95 (air) or 60X/NA1.27 (water immersion) objective on a spinning-disk microscope (CellVoyager 7000, Yokogawa), equipped with sCMOS cameras (0.1625μm or 0.10833 μm pixel dimensions, for 40X/60X objectives, respectively). Approximately 60 confocal Z-slices were obtained per imaging site with step sizes of 0.25μm or 0.33μm. Calculation of cell volume from bead images is described in “Quantification and Statistical Analysis”.

#### Fluorescence exclusion method (FXm)

FXm was performed as previously described (Cadart et al., 2017). Briefly, a PDMS chamber with height 21.8μm was prepared using an epoxy mould provided by L. Venkova and M. Piel (Institut Curie, Paris, France). The chamber was then attached to a 35mm FluoroDish using plasma treatment, and internal surfaces were coated with 50ug/mL human fibronectin for 1 hour at 37°C. Chambers were then incubated in cell growth media overnight, and washed once more with growth media. Cells were then trypsinised and loaded as a suspension into the FXm chamber. After allowing cells to adhere for 4 hours, media was replaced with fresh media containing 1 mg/mL Dextran-Alexa Fluor 488 (10,000 MW). FXm measurements were then performed on live cells at 37°C, 5% CO_2_ using an epifluorescence microscope (VisiTIRF, Visitron) with a 20X/0.4NA objective lens (Nikon). Fluorescently labelled dextran was excited with 450-490nm (SOLA lightengine) and emission collected from 500-550nm. Data analysis was performed using custom software as previously described (Cadart et al., 2017). For comparison with cell volume reconstruction method, cells were fixed in the chamber using 4% PFA and bead attachment was performed manually, as described above. Chamber-level overviews were derived by computationally stitching images from both methods, and the same single cells were identified in both datasets by aligning these overviews.

#### Single-molecule RNA fluorescent in situ hybridisation (smFISH)

Single-molecule FISH was performed as previously described (Battich et al., 2013), using ViewRNA high-content screening assay kit (Affymetrix) with Type 1 (Figure 1) or Type 1, 6, and 10 (Figure 3) primary probes and signal amplification kits (Affymetrix). Samples were imaged on a spinning-disk microscope using a 40X/NA0.95 air objective with step-size of 1.0μm. Computational identification of spots was performed in 3D (Figure 1, with cell volume measurement), or in 2D from maximum-projected images (Figure 3, genetic perturbation screen), as previously described (Stoeger et al., 2015).

For smFISH experiments in the (tertiary) genetic perturbation screen (Figures 3H-I, S7), we focused on nine genes (*RHEB*, *STX6*, *CTCF*, *GLS*, *RELA*, *HPRT1*, *CSPG4*, *NCOA4*, *TERF2IP*). These genes are expressed throughout the cell cycle, encompass diverse biological functions, and have a range of estimated mRNA half-lives (2.3h to >24h (Tani et al., 2012)). They were also previously shown to have cytoplasmic mRNA abundance that scales with cell size (Figures 1A, S1I) (Battich et al., 2015). The genes targeted by siRNA in this experiment are shown in Figure S7E. Only one of the nine genes (*HPRT1*) had mean mRNA abundance that correlated with EU incorporation across perturbations (Figure S7H-I). *HPRT1* was typically down-regulated in perturbation conditions with reduced EU and up-regulated in perturbations with increased EU – a trend consistent with its biological function in nucleotide metabolism.

#### In situ metabolic labelling of nascent RNA

Nascent RNA was visualised using metabolic labelling as previously described (Jao and Salic, 2008), with modifications. Briefly, adherent cells were cultured in complete media at 37°C, 5% CO_2_ for 2-3 days. 5-ethynyl uridine (EU) was then dissolved to a concentration of 2mM in pre-warmed complete media. EU was added to cells by partially aspirating growth media and dispensing an equal volume of 2mM EU using a BioTek washer-dispenser (e.g. 30μL 2mM EU added to 30μL residual for 384-well plates: final EU concentration = 1mM). Cells were then incubated for 20 or 30 min at 37°C, 5% CO_2_, before fixation with 4% PFA at room temperature for 20-30 min. After fixation, cells were permeabilised with 0.5% Triton X-100 and washed 3 times with TBS (50mM Tris pH 8.0, 150mM NaCl).

To render nascent RNA fluorescent, we prepared sufficient volume of click reaction master mix for all wells on a plate at 1.5x concentration, as follows: 75μM Alexa Fluor 488 azide or Alexa Fluor 647 azide, 3mM CuSO4, 150mM Sodium ascorbate, in TBS. Click reaction was dispensed using a BioTek washer-dispenser and incubated for 30 min at room temperature before washing cells 3x into PBS. The click reaction is detrimental to the intensity of several fluorophores including the fluorescent beads used for cell volume measurement and also GFP. To combine these with EU metabolic RNA labelling, we fixed, permeabilised and imaged either GFP or beads before click reaction. Images were subsequently aligned using the DAPI channel, which was included in both imaging rounds, using the computational procedure described previously for 4i (Gut et al., 2018).

#### Immunofluorescence and iterative indirect immunofluorescence imaging (4i)

For PCNA staining in non-4i immunofluorescence experiments, and for the genome-wide screen, cells were blocked with 1% BSA in PBS for 1 hour and then incubated for 2 hours with rabbit anti-PCNA antibody dissolved in 1% BSA in PBS. Pol II immunofluorescence in non-4i immunofluorescence experiments was performed using Intercept blocking buffer (LI-COR Biosciences) for blocking and antibody incubations.

Immunofluorescence-based quantification in combination with *in situ* RNA metabolic labelling across multiple imaging cycles was validated using MRC5 POLR2A-GFP cells (Steurer et al., 2018). We first pulsed cells with EU, then fixed and imaged GFP and DAPI. Subsequently, we performed click reaction and POLR2A immunofluorescence and then re-imaged. Images from the two rounds were aligned to the DAPI channel, using the computational procedure described previously for 4i (Gut et al., 2018). We then quantitatively compared POLR2A immunofluorescence with GFP intensity (Figure S8A-B): IF and GFP were highly correlated (*r* = 0.92) and both showed similar partial correlation with EU (*r*_*x*,EU|nucl. area_ of 0.49 and 0.46, respectively).

4i was performed as previously described (Gut et al., 2018) with two modifications: Intercept blocking buffer (LI-COR Biosciences) was used for all blocking, primary and secondary antibody incubations, and 50mM HEPES was included in imaging buffer – which was adjusted to a pH of 7.4. Before 4i experiments, all antibodies were tested for compatibility with elution buffer using the following criteria: similar staining on normal and elution-buffer treated cells, minimal residual signal after elution and restaining with secondary antibody. After 4i, we also validated that agreement between replicates was high (Figure S8E) and that the technical variability between control (scrambled siRNA) wells was much less than the differences induced by the perturbations (Figure S8F). To ensure successful antibody elution in each cycle, we included elution controls in each imaging cycle. This consists of reprobing a test well with a secondary antibody in the imaging cycle after it was stained with primary and secondary antibody and imaged. Efficient elution was verified in all cases.

#### Poly(A) RNA fluorescent in situ hybridisation (FISH)

Poly(A) FISH was adapted from published smFISH protocols (Raj et al., 2008) for high-throughput liquid handling. Briefly, cells were fixed for 15 min in 4% paraformaldehyde, permeabilised in 0.2% Triton X-100 for 15 min (secondary screen) or 70% EtOH for 1 hour (HCT116 experiments). If required, click reaction was then performed before applying FISH probes. Cells were then washed into 2X saline sodium citrate buffer (SSC) containing 10% (v/v) formamide. Before hybridisation, samples were transferred to hybridisation buffer (2X SSC, 10% (v/v) formamide, 100mg/mL dextran sulfate), and pre-incubated for 1 hour at 37°C. Fluorescently labelled DNA oligonucleotide probes, dT(30)-Atto488 or dT(30)-Cy5, were purchased from Microsynth as labelled and HPLC-purified. These were diluted into the hybridisation buffer and applied to cells at a final concentration of 400nM. After 16 hours at 37°C in a rotating incubator, cells were washed into 2X SSC, 10% formamide, incubated again at 37°C for 1 hour, before washing into 2X SSC. For combining with subsequent immunofluorescence, poly(A) FISH was imaged before immunostaining. Images were aligned using the DAPI channel, which was included in both imaging rounds, using the computational procedure described previously for 4i (Gut et al., 2018).

#### StrandBrite total RNA staining

RNA StrandBrite was used according to manufacturer’s instructions. We confirmed that RNA StrandBrite staining is specific for RNA by incubating cells with 0.5mg/mL RNAse A at 37°C for 30 min (Figure S6A). RNA StrandBrite green is spectrally distinct from poly(A) FISH when using Cy5-labelled oligo(dT) probes, so the two assays were combined. However, during protocol optimisation, we found that RNA StrandBrite staining was not stable during the poly(A) FISH protocol, showing an altered localisation pattern and reduced signal intensity after FISH. However, poly(A) FISH was not affected by prior RNA StrandBrite staining. To combine the two assays, we therefore performed RNA StrandBrite staining and imaging before poly(A) FISH. Images were aligned using the DAPI channel, which was included in both imaging rounds, using the computational procedure described previously for 4i (Gut et al., 2018).

### QUANTIFICATION AND STATISTICAL ANALYSIS

#### Image processing

The majority of image analysis was performed in TissueMAPS, using previously developed methods (Stoeger et al., 2015). All images were corrected for illumination artefacts as previously described (Stoeger et al., 2015). In some cases, we also developed novel algorithms, or used pixel classification (Sommer et al., 2011) or neural networks (Stringer et al., 2021) to aid in segmentation tasks. This is indicated where applicable.

##### Nuclear and cell segmentation

Nuclei were segmented by adaptive thresholding of the DAPI or H2B signal. After separating clumped nuclei and excluding small objects, cells were then segmented using total protein stain (succinimidyl ester), using these nuclei as seeds for a watershed, as described previously (Stoeger et al., 2015). The analysis pipeline for the genome-wide screen is described in (Müller et al., 2021). For MRC5 cells and the 4i experiment, we used the cellpose generalist nuclear segmentation neural network (Stringer et al., 2021) without additional training because it yielded superior results for separating touching nuclei. However, we found that nuclei identified by cellpose were not always completely contained within the DAPI-positive regions of the image, or did not completely fill the nucleus. Cellpose nuclei were therefore further refined using adaptive thresholding and/or watershed expansion using the DAPI signal.

##### Nucleolar segmentation

The nucleolus was segmented from DAPI and Alexa Fluor 647-succinimidyl ester images using pixel classification in Ilastik (Sommer et al., 2011), as depicted in Figure S2L and described previously (Müller et al., 2021).

##### Cell volume measurement

To perform computational 3D cell reconstruction, cells were first segmented in two dimensions using procedures described above. We then localised beads in 3D, using an approach previously developed for smFISH (Raj et al., 2008; Stoeger et al., 2015). The brightest voxel of each segmented bead in the original image was taken as the bead centre. Next, the slide surface was estimated by fitting a plane in 3D through all beads that were detected outside of cells. The 3D positions of beads are then recalibrated relative to the slide surface, so that the list of (x,y,z) coordinates for each cell represents the height of the upper cell surface above the slide. To reconstruct the cell surface, outlier beads were excluded by fitting a 3D alpha shape (Da et al., 2020) to the list of coordinates for each cell, as well as a set of points at the 2D cell periphery, which are computationally anchored to the slide surface. The upper cell surface was then linearly interpolated using the remaining points. Code is written in python, making use of numpy (Harris et al., 2020) and scipy (Virtanen et al., 2020), and was integrated into TissueMAPS. We directly compared cell volumes estimated with this method with those obtained by fluorescence exclusion method (FXm) (Cadart et al., 2017) finding good agreement at the single-cell level (Figures S1B-E). We also found that cell volumes correlated well with other measures of cell size, such as nuclear and cell area, and total protein levels, visualised with succinimidyl ester staining (Figure S1F). Correlations of volume with nuclear area were slightly higher for HeLa than for “flatter” 184A1 cells and keratinocytes (Figure S1H), which often form long projections. Both protein content and nuclear area were approximately proportional to cell volume in HeLa cells (Figure S1G) and are therefore excellent proxies for cell volume in unperturbed cells.

##### Chromatic aberration correction – 4i

Chromatic aberrations were evident when comparing different imaging channels in the 4i experiment. To correct these, we imaged multicolor fluorescent beads and used these to fit parameters of an affine transformation aligning each channel to the 568nm channel. These channel-specific transformations were then applied to all images for the 405nm, 488nm and 647nm channels, using a custom procedure developed in python making use of scipy (Virtanen et al., 2020), numpy (Harris et al., 2020) and scikit-image (Walt et al., 2014).

#### Data cleaning

After cell segmentation, we trained supervised machine-learning models (support-vector machines) to exclude certain acquisition and segmentation artefacts using the TissueMAPS framework. This interactive procedure is similar in nature to previous software developed in the lab, CellClassifier (Rämö et al., 2009). Briefly, it involves manually selecting cells with certain properties as training data for a classifier, visualising the results of the classification, and selecting additional examples to refine the classification (Figure S2A). Throughout this work, we excluded border cells (those with pixels touching the image boundary), mitotic/apoptotic cells (identified using DAPI texture features), cells with cytoplasmic DAPI, polynucleated cells, and all cells from sites that were not in focus. In the genome-wide screen, we also trained a classifier to exclude mis-segmented cells (Müller et al., 2021).

##### Additional data cleaning – cell volume measurement

Cell volume measurement involves computationally fitting a plane in 3D to define the slide surface. When cells were too densely packed, this cannot be achieved accurately, preventing accurate determination of cell volume. We therefore did not compute cell volumes for these sites, which were automatically identified based on the proportion of the image surface identified as unoccupied. We also excluded cells with insufficient bead density, or those which were not fully captured in the confocal volume. These were identified and removed using manually defined thresholds during exploratory data analysis.

##### Additional data cleaning – smFISH

smFISH probes were dispensed into 384-well plates using automated liquid handling, as previously described (Battich et al., 2013). Inaccuracies in handling of small volumes led to a ‘quadrant effect’ in which the upper left well in each group of 4 wells contained smFISH spots with weaker intensity. Spots in these wells could not be unambiguously distinguished from background, so these were excluded from further analysis. This reduced the number of perturbations studied from the target of 63 to 50.

##### Additional data cleaning – 4i

The 4i perturbation experiment contained 160 wells and 18 antibodies (2880 antibody/well combinations). To identify outlier wells, we compared mean intensity measurements for each well and each antibody. Where replicates differed in any stain by more than 30%, the quantitative data and underlying images were manually examined. In most cases, the discrepancy was due to cell density differences between replicate wells, and all data was retained. In nine cases, we identified that staining was uniformly lower in one replicate, likely due to errors in automated liquid-handling. These antibody/well combinations (0.3% of 2880) were excluded. For multivariate analysis (e.g. UMAP), affected wells were entirely excluded, resulting in a single replicate being used for 6 of the 66 siRNA conditions studied: CBX3, KDM4D, POLR3E, PGAP3, RAB5C, SNRPF.

##### Additional data cleaning – plasmid transfection experiments

In agreement with previous work, we found that plasmid DNA could be detected in the cytoplasm of transfected cells using poly(A) FISH (Greenberg et al., 2019). This was prominent at 12h and 24h after transfection, but was reduced after 48h, likely due to asymmetric partitioning during cell division (Wang et al., 2016). To identify cytoplasmic DNA foci, we used Ilastik-based pixel classification, with poly(A) FISH and DAPI as a two-channel input. We excluded all cells containing these foci.

#### Background subtraction

Quantitative intensity values for stains such as DAPI, succinimidyl ester, RNA StrandBrite, and poly(A) FISH were background subtracted using a constant value measured in a region outside the objects. For immunofluorescence measurements, background values were taken from a region within the cell for a well in which the primary antibody was omitted. For EU, background values were taken from a well that was not incubated with EU but was subject to click reaction. Since click reaction staining in the absence of EU showed a similar staining pattern to succinimidyl ester, we used a linear modelling approach to predict the expected background based on succinimidyl ester intensity. This slightly outperformed a constant-value background subtraction approach.

#### Data normalisation

In some cases, we observed variation in staining intensity for different rows of the plate. These inconsistencies arise because of the small differences in reagent volumes delivered by the automated liquid handling system, which operates row-wise. This was assessed routinely during data analysis by manual inspection of intensity distributions after background subtraction. When necessary, we multiplied all background-subtracted intensity values by a correction factor to equalize the medians of the rows. Where possible, in genetic perturbation experiments, only negative control (scrambled siRNA) wells were used for normalisation. The same correction strategy was used to correct for plate-to-plate variation in multi-plate experiments.

##### Additional normalisation – Poly(A) FISH

DNA probes for poly(A) FISH and also StrandBrite reagents were dispensed using a 96-well pipette head into 384-well plates. This resulted in a quadrant-specific artefact in intensity that was corrected in the same way as described for row- and plate-corrections described above.

##### Additional normalisation - 4i

Incubation in 4i imaging buffer (Gut et al., 2018) is detrimental to signal intensity for some antibodies. This is observed as a reduction in intensity over the time of image acquisition (12-16 hours per imaging cycle). We corrected for this artefact by fitting an aympototic regression model (SSasymp function in R) to the median signal intensity of all scrambled siRNA control cells as a function of acquisition time (Figure S8C-D). Fitting was achieved using the nlsLM function from the minpack.lm package (Elzhov et al., 2016), after removing outlier control wells. We then multiplied all values by a correction factor derived from this curve and used it to multiplicatively scale all data based on acquisition time. This resulted in excellent agreement between intensity values of replicate (non-control) wells, which were typically imaged 4-5 hours apart (Figure S8E).

We also observed smaller well-specific staining biases that could not be explained by time-dependent intensity reduction, but were also not clear technical outliers. These represent technical variation inherent to the experiment. Because these can lead to biases when computing correlations among populations of single cells from different wells, it was sometimes necessary to correct these. This was achieved by multiplicative centering of well medians between experimental duplicates. To ensure that this did not bias our results, we verified that correlation coefficients across the combined populations were similar to those seen within individual wells (Figure S10C). Importantly, this correction was only applied to allow pooling cells from multiple wells and was therefore not performed whenever values from individual wells were summarized and considered as replicates (for example for fold-change calculations, or correlations between well-averages).

#### Cell cycle classification

Mitotic cells do not incorporate EU, and were therefore not of interest for this work. We first identified and removed these as described in ‘Data cleanup’ (Figure S2A). G1 and G2 cells can be distinguished based on DNA content measured from DAPI staining intensity. However, identifying S-phase cells requires more information. The gold standard for identifying cells undergoing DNA replication is to use 5-ethynyl-2’-deoxyuridine (EdU) metabolic labelling, which is similar to EU labelling except that cells are pulsed with EdU instead of EU to label newly synthesized DNA.

Because EU and EdU detection rely on the same click chemistry, they cannot be combined on the same sample. We therefore included several wells of EdU in our RNA metabolic labelling experiments, and used PCNA immunofluorescence as an S-phase marker. PCNA is localised to DNA replication foci and has a characteristic punctate localisation in S-phase cells. By performing PCNA immunofluorescence on all wells, we could identify S-phase cells using EdU in the EdU-treated wells and train a classifier based on PCNA to identify S-phase cells also in the EU-treated wells (Figure S2B).

More specifically, after background subtraction and feature normalisation as described above, we trained a random forest classifier using the randomForest package in R (Liaw and Wiener, 2002), to classify S-phase cells using PCNA and DAPI texture features. We used thresholded EdU intensity as a ground truth for cells that are undergoing DNA replication (Figure S2C). Classifiers were typically 95-98% accurate (Müller et al., 2021) (Figure S2D). Remaining non-S-phase cells were classified as G1 or G2 based on DNA content (Figure S2E-F). In experiments with genetic perturbations, we included siRNA knockdown of GRIP2, SBF2, NUDT4 and scrambled siRNA controls in the EdU training data to ensure robustness to cell size and morphology perturbations, as described and validated in the accompanying manuscript (Müller et al., 2021).

#### Quantitative data analysis

After extracting quantitative features from images, quantitative analysis for individual experiments was done in R (Team, 2014), relying heavily on tidyverse packages (Wickham et al., 2019).

##### smFISH

Linear regression to predict spot count from cell volume was performed in R using the lm function with model *n*_spots_ = *a* + *bV*. Model fits were quantified using the coefficient of determination (*R*^2^), calculated using *R*^2^ = 1 – RSS/TSS, where the residual sum of squares, 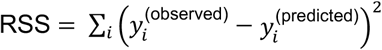, and the total sum of squares, 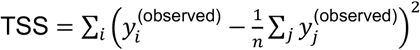 for *n* observations.

Following (Padovan-Merhar et al., 2015) we quantified the fraction of predicted mRNA abundance that is volume independent (*a*/(*a* + *bV*)) or volume-dependent (*bV*/(*a* + *bV*)) for all genes measured. This is only interpretable for *a*,*b* ≥ 0, so we omitted genes where *a* was significantly less than zero, and set *a* = 0 for intercepts that were slightly negative but not significantly (*p* > 0.05) different from zero.

##### Fold-change estimates

To compute fold-changes in intensity values observed between conditions, we took the median across all cells in each well (or a subset of cells such as a certain cell-cycle stage, as specified). We then compared these well-medians of a particular experimental condition with the well-medians of the appropriate control wells by computing all pairwise ratios (for example *n* replicate experimental wells with *k* control wells gives *n×k* fold-change estimates). We either summarized all *n* × *k* ratios as a boxplot or reported summary statistics for these ratios such as their mean and standard deviation.

##### Partial correlations

Pairwise partial correlation coefficients (Figure S10G) were calculated using the pcor function in the ppcor package in R (Kim, 2015). Such partial correlations represent correlations in which the effects of all other variables on the two under consideration are first taken into account. In cases where we account for the effects of a single additional variable before calculating the correlation (e.g. nuclear area (*r*_*x*,EU|nucl. area_)), partial correlations were calculated using the pcor.test function.

##### Multiple linear regression

Linear regression was performed with sum nuclear EU intensity as the response variable and different combinations of predictors. In cases where there were more than three predictors, we standardised all variables, performed principal component analysis (using the prcomp function in R), and kept enough principal components to capture 99.9% of the variance in the data. For 4i data, we omitted outlier antibody/well combinations, identified as described above. Multiple linear regression was performed using the ‘lasso’ method with 10-fold cross-validation using the glmnet R package (cv.glmnet) (Friedman et al., 2010). To fit models, we omitted each replicate well from the training data, trained a model on the remaining data, and then predicted values for the non-training well. This procedure was repeated for each replicate well, to generate predicted values for every cell. The procedure avoids testing models on data that was used to fit the model, but does not arbitrarily assign a training and test set.

##### ‘Residual’-based corrected intensities

Changes in EU incorporation or RNA abundance in genetically perturbed cells are confounded with changes in cell size, cell cycle and/or population-context. To correct for these effects, we used regression to linearly account for any confounding variables at the single cell level, and then used the residuals of this regression as a ‘corrected’ intensity measurement. In all cases, regression models were trained on control cells from the same experiment, so the residual measurement represents the deviation of a particular cell from that expected for a control cell. Variables that were corrected in this way are described throughout the main text, including mean nuclear EU, sum nuclear EU, sum nucleolar EU and sum nucleoplasmic EU in metabolic RNA labelling experiments; RNA StrandBrite and poly(A) FISH in the RNA abundance screen; smFISH transcript abundance counts; POLR2A and EU intensities in 4i dataset, and POLR2A-GFP in MRC5 POLR2A-GFP cells.

The particular confounding variables considered differed between experiments, depending on which additional information was available from the other cellular stains, and also the type of analysis being considered. The variables whose effects are included in the correction are identified in the text. Typically, these included some or all of the following: nuclear and cell area, cell cycle stage, total protein content (sum cell succinimidyl ester), and local cell density. We now describe in detail how these corrections are performed for a single response variable.

We first identified if there were any outlier control wells. These are important to remove before using as training data for the regression models. To identify these, we trained a model on all replicate control wells except one, and then predicted the response variable in the well omitted from model training. Quantifying this model fit using using *R*^2^, we then used a one-sided boxplot rule on *R*^2^ to classify wells as outliers if *R*^2^ < *Q*_1_(*R*^2^) – 1.5 × IQR(*R*^2^). This specifically identifies wells that are poorly predicted by models trained on all other replicate wells. Such ‘outlier’ wells are retained in the dataset but were omitted for training the regression model used to correct non-control wells to ‘residual’-type variables.

Regression models were trained in several different ways, depending on the number of predictors. For example, residual mean EU is derived from a model containing a single categorical predictor: cell cycle stage, so the ‘residual’ simply corresponds to subtracting a cell-cycle specific value from each measurement. In contrast residual sum EU for the nucleus, nucleolus and nucleoplasm contains total protein content a continuous predictor as well as cell cycle as a discrete predictor, so simple linear regression was used. For more than two predictors, we used cross-validated ‘lasso’ multiple linear regression using the glmnet R package (cv.glmnet) (Friedman et al., 2010). In cases with several correlated predictors we also included a PCA-based dimensionality reduction of the predictor variables (maintaining 99.5% of variance) before regression. Typically, these values were averaged on a per-well basis by taking the mean. These “mean residual” measurements were finally standardised to the mean and variance of scrambled siRNA control wells using the Z-score, to give a measurement in units of standard deviation of scrambled siRNA controls.

##### Genome-wide and secondary screen analysis

Quantitative data analysis of single-cell intensity measurements from the genome-wide screen and secondary screen were described previously (Müller et al., 2021). Lower and upper hit thresholds correspond to posterior probabilities of 0.5 and 0.85 for perturbations being reproducibly observed outside the 1st and 99th percentile of scrambled siRNA controls. Calculation of these thresholds was also described previously (Müller et al., 2021). ‘Low cell number’ in the screen corresponds to 500 interphase cells imaged and segmented after data cleanup. There were 258 conditions with less than 500 cells in the genome-wide screen (approximately 1% of gene perturbations). These were included in certain overviews (e.g. Figures 2C–2D) but were typically excluded, as noted throughout, for example in hit scoring and functional annotation enrichment.

Hit scoring made use of two residuals-based models: either predicting ‘sum EU’ (with cellular protein content and cell-cycle stage as predictors) or ‘mean EU’ (EU intensity averaged over nuclear area, with only cell cycle stage as a predictor). The ‘sum EU’ model explicitly accounts for cell size, while ‘mean EU’ implicitly takes cell size changes into account because nuclear size also scales with cell size (Cantwell and Nurse, 2019). In unperturbed cells, both nuclear area and protein content are proportional to cell volume (Figure S1G). These two residual-based measurements were highly correlated (*r* = 0.94), but ‘mean EU’ was slightly less variable between wells and screens and was therefore more sensitive, possibly because it avoids technical variability in measurement of cellular protein content (Figure S3B, S3D) (Müller et al., 2021). Mean residual sum EU for the nucleolus and nucleoplasm were calculated as described for the sum EU model. Comparison of mean residual measurements between primary and secondary screens was used to define hit thresholds in all cases, as described previously (Müller et al., 2021).

The selection of 436 siRNA perturbations for the secondary screen (RNA metabolic labelling and poly(A) FISH / RNA StrandBrite) was done in an automated manner designed to preserve both functional and phenotypic diversity of the panel, as described previously (Müller et al., 2021). Selection of genes for the final panel of 63 perturbations for detailed characterization by 4i was done manually based on enriched functional annotations and results of the secondary screen. Gene lists and other metadata are provided together with the experimental results at the locations listed in Key Resources.

##### Estimation of transcriptional scaling in genome-wide screen

Linear regression to determine slope and intercept values for the relationship between sum nuclear EU and cellular protein content was performed for all perturbations in the genome-wide screen with at least 500 cells. This was done in R using the lmrob function from the robustbase package (Todorov and Filzmoser, 2009) with lmrob.control(“KS2014”). If there were less than 30 cells in a particular cellcycle stage, those cells were omitted. This analysis was complicated by the presence of strong overall changes to EU incorporation and incomplete phenotypic penetrance of perturbations, which together resulted in cell populations with large cell-to-cell variability in EU incorporation. Across perturbations, slopes were correlated with mean EU intensity, however they were highly variable even between replicates (Figure S3F).

##### Robust coefficient of variation (RCV)

Despite data clean-up, single cell data can contain spurious outlier observations, particularly in genetic perturbation experiments. To estimate the population variability, we therefore used a robust analogue of the coefficient of variation given by *RCV_M_* = 1.4826 × MAD / median, where MAD is the median absolute deviation, given by *MAD*(*x*) = median(|*x* - *m*|) where *m* = median (*x*) (Arachchige et al., 2020).

##### 4i data analysis – UMAP

After background subtraction and data normalisation, we combined mean nuclear intensities and textures for all markers except EU, together with mean cytoplasmic intensities and textures of PABPC1 and PABPC4, nuclear and cell morphology measurements, and measurements of local cell density (Snijder et al., 2012) into a matrix of 436,593 cells by 718 features. We then standardised all features to scrambled siRNA control cells using the robust z-score: *x* → (*x* — median(*x*_scrambled_))/(1.4826 × MAD(*x*_scrambled_)] and performed dimensionality reduction via principal component analysis (PCA) using the R function prcomp, keeping 95% of the variance (first 108 principal components). We then randomly sampled 1400 cells from each siRNA perturbation condition and used this PCA-transformed data as input for umap using the R uwot package (Melville, 2020) with parameters a = 0.25 and b = 0.9. Finally, we used the umap_transform function to embed the full dataset in UMAP space.

#### Systems-level analysis

##### Functional annotation enrichment scoring

We retrieved GO biological process, KEGG pathways, and Reactome annotations for all genes assayed using the GeneSets.Homo.sapiens R package (Simillion, 2020), and removed any genes with zero annotations. Genes were then ranked using a quantitative phenotype (e.g. mean residual EU) for both ‘up’ and ‘down’ hit classes separately. These ranked lists were used to search for overrepresented annotations, adapting a previously published method (Green and Pelkmans, 2016). Briefly, for each annotation, we count the number times, *k* that a given annotation is observed between rank 1 and *n*, and compute the probability that this occurs by chance, given the number of assayed genes, *N*, of which *K* have the corresponding annotation. This probability is given by the hypergeometric distribution function:

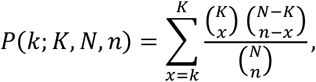

which is implemented in R as phyper(k-1,K,N-K,n,lower.tail = FALSE). This calculation is repeated for all ranks from *n* = 1,…,*n_t_* where *n_t_* is the rank at which the probability threshold exceeds 0.5 (50/50 whether a hit is reproducible). The functional annotation enrichment score (FAES) is then calculated using the minimum value of this probability, FAES = −log_10_(*min_n<nt_P*(*k*;*K*,*N*,*n*)). We also note the rank *n* = *n*_min_ at which this minimum occurs.

To represent these enriched annotations as a network, we selected up and down-enriched annotations with FAES > 2, removing any annotations with less than 20 or more than 3500 genes. We then measured the pairwise similarities between annotations, using the Cohen’s kappa statistic, κ, (Cohen, 1960) provided by the Kappa function in the vcd R package (Meyer et al., 2020). Annotations for which κ > 0.85, were merged into a group, keeping the highest FAES score for an annotation in that group. We then built a graph of these enriched annotations, where edges between nodes represent annotations with some overlap in the gene set (κ > 0.15). The network was finally visualised in Cytoscape (Shannon et al., 2003) using spring-embedded layout based on the edge weights (κ), also making use of enhancedGraphics (Morris et al., 2014).

To calculate FAES for the nucleolar-specificity of annotations that were enriched for reduced EU incorporation, we calculated “nucleolar preference” of a perturbation, which we defined as the residual of “residual nucleolus sum EU” after regressing out “residual nucleoplasm sum EU” (Figure S5G). Nucleolar preference gives a quantitative measurement of the reduction in nucleolar EU compared to that expected based on nucleoplasmic EU. Nucleolar preference is positive for perturbations with stronger nucleolar EU reduction than nucleoplasmic EU reduction. We then performed rank-based enrichment analysis for nucleolar preference using all genes with *p*_posterior_ > 0.5 for either residual nucleoplasm sum EU or residual nucleolus sum EU (Figure S5I). Finally, we compared this to FAES for reduced residual nucleolus sum EU, and identified annotations that were enriched in both analyses (Figure S5J).

##### Gene interaction networks

After grouping enriched annotations using Cohen’s κ, as described above, we obtained proteinprotein association networks from the STRING database (Szklarczyk et al., 2018) for all genes in a certain annotation group. We then constructed networks in which genes are connected if their aggregated interaction score is > 0.7. Networks were visualised in Cytoscape (Shannon et al., 2003) using spring-embedded layout based on the edge weights (association score).

##### Hierarchical clustering

Hierarchical clustering was performed in R using the seriation package (Hahsler et al., 2008) using Ward’s algorithm with optimal leaf ordering. Heatmaps were visualised with the ComplexHeatmap package (Gu et al., 2016).

#### Mathematical modelling

Ordinary differential equation (ODE) modelling was performed in R (Team, 2014) using the deSolve (Soetaert et al., 2010) package. Parameter optimisation was done using the optimx (Nash and Varadhan, 2011) package, using least-squares minimisation on log-transformed data and predictions.

##### Minimal model without RNA-based feedback

(Steurer et al., 2018) developed a model of the RNA Pol II transcription cycle in which Pol II can exist in one of four states, which we here refer to as “Unbound” “Initiating”, “Paused” and “Elongating”. Their model was parameterised and validated using Pol II fluorescence recovery after photobleaching (FRAP) of MRC5 POLR2A-GFP cells, using several chemical inhibitors with known mechanism of action. To adapt this model to an ordinary differential equation framework, we modelled the Pol II transcription cycle using the reaction network depicted in Figure S12A. Using mass-action kinetics, this gives rise to the following system of ODEs in terms of concentration,

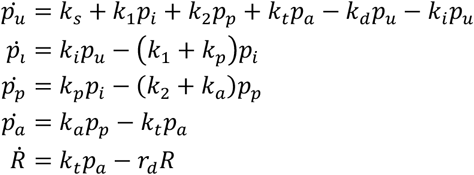

where *p_u_*,*P_i_*,*p_p_*,*p_a_* are unbound, initiating, paused, active (elongating) Pol II, is RNA, and 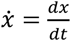 denotes the time-derivative of *x*. Note that RNA is created at a rate *k_t_p_a_*, the same overall rate at which Pol II terminates elongation, meaning that we do not consider premature termination of transcription (other than transcription aborting after pausing). We set *k_t_* using the FRAP recovery timescale of the elongating state (1370 seconds), and then optimised the remaining Pol II transition rate parameters (*k_i_*,*k_p_*,*k_a_*,*k*_1_,*k*_2_) to obtain the observed fractions of Pol II states at steady-state (7% Free, 10% Initiating, 23% Paused, 60% Elongating). A further constraint from (Steurer et al., 2018) relates to the finding that only 12.7% of polymerases that attempt initiation proceed to promoter pausing, and only 7.6% of promoter-paused polymerases continue to productive elongation. This allowed us to specify *r*_1_ and *k*_2_ in terms of *k_p_* and *k_a_*, respectively (*k*_1_ = 6.87 × *k_p_*, *r*_1_ = 12.16 × *k*_a_).

After setting the total concentration of Pol II in the nucleus at 10^5^ molecules pL^-1^ (Steurer et al., 2018), there remains a single free parameter (*k*_s_), which sets the timescale of Pol II synthesis / degradation. This parameter has units of molecules pL^-1^s^-1^, so a constant value corresponds to an absolute (molecular) rate of Pol II synthesis that scales with cell volume. To extract *k_s_* from our data, we simulated triptolide-based transcriptional inhibition by setting *k_i_* = 0, and AZD4573-based CDK9 inhibition by setting *k_a_* = 0, keeping all other parameters unchanged. We then optimised *k_s_* to fit the combined data from these two experiments, with POLR2A-S2P identified as elongating Pol II (Heidemann et al., 2013) (POLR2A-S2P = *p_a_*). Parameters and their final fitted values are listed in Table S1. To investigate alternative models in which multiple Pol II states were subject to degradation (Figures 6B, S12B), we added extra –*k_d_p_x_* (for *x ∈ i,p,a*) terms to the relevant ODEs, and refit *k_s_*, as described above.

##### Adding RNA-based feedback

Having parameterised the Pol II model, we then included RNA-based feedback as either a Hill-type activation of Pol II degradation *k_d_* → *k_d_*(*R*/*K* + *R*), or a Hill-type repression of *k_i_*,*k_p_*,*k_a_* (with Hill coefficient *n* = 1), *k_x_* → *k_x_*(*K*/*K* + *R*) for *x ∈ i*,*p*,*a*. In all cases, this change required re-fitting of some of the parameters from the base model to achieve the correct fractions of Pol II in each state and the same total Pol II concentration (Table S1).

To fit the RNA degradation rate and feedback parameters, *r_d_*, and *K*, we simulated auxin-induced degradation of DIS3 by setting,

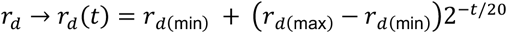

which simulates the exponential decrease in DIS3 abundance observed experimentally (Davidson et al., 2019). *r_d_* does not decrease to zero because this parameter represents all nuclear mRNA degradation and export, as well as dilution due to nuclear growth. We used *r*_*d*(min)_ = *r*_*d*(max)_/4, which assumes that DIS3 is responsible for ~3/4 of nuclear mRNA degradation, however the actual value does not qualitatively affect the differences seen between models. Assuming POLR2A = *p_u_* + *p_i_* + *p_p_* + *p_a_*, POLR2A-S5P = *p_p_* + *p_a_*, POLR2A-S2P = *p_a_* and mRNA = *R*, we optimised *r_d_*, and *K* to fit mRNA (poly(A) FISH) and Pol II (POLR2A, POLR2A-S5P, POLR2A-S2P immunofluorescence) measurements from DIS3-AID experiments. The magnitude of Pol II changes in the model are governed mostly by *K* (representing the concentration of RNA with half-maximal effect on *k_x_*). However, the precise value of *K* does not affect the qualitative differences between models in terms of the relative effects on POLR2A, POLR2A-S5P, and POLR2A-S2P.

To analyse how steady-state RNA levels are affected by changes to model parameters (parameter sensitivity analysis) in Figure S12D-E, we focused on the model which was most consistent with the experimental data (Pol II feedback on “paused” to “elongating” transitions: *k_a_* → *k_a_*(*K*/*K* + *R*)). We varied either *k_s_* (to simulate over- or under-production of Pol II) or *r_d_* (to simulate changes to nuclear RNA degradation, export or growth rate), and then re-calculated the steady-state values of POLR2A or RNA for this new parameter set (represented relative to steady state levels obtained from the best-fit parameters). In Figure S12E, we further considered how non-linear feedback of RNA affects buffering capacity by *k_a_* → *k_a_*(*K^n^*/*K^n^* + *R^n^*) for *n* ∈ 1,2,4. In Figure S12F-G, we considered how models respond dynamically to perturbation of cell volume. For example, a 20% cell volume increase was modelled by dividing each of the steady-state concentrations by 1.2. We then simulated the return to steady-state concentrations using numerical integration.

---

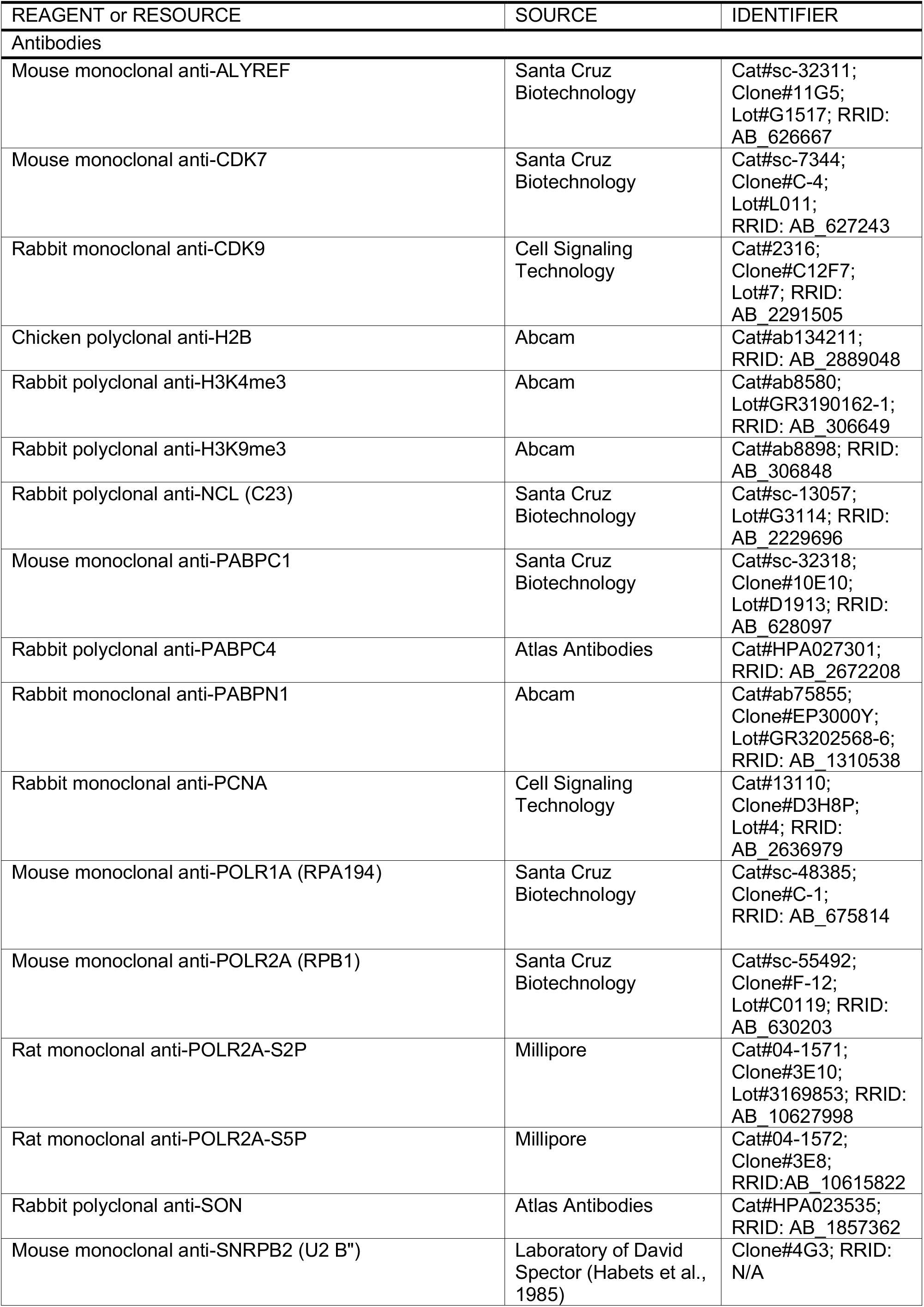
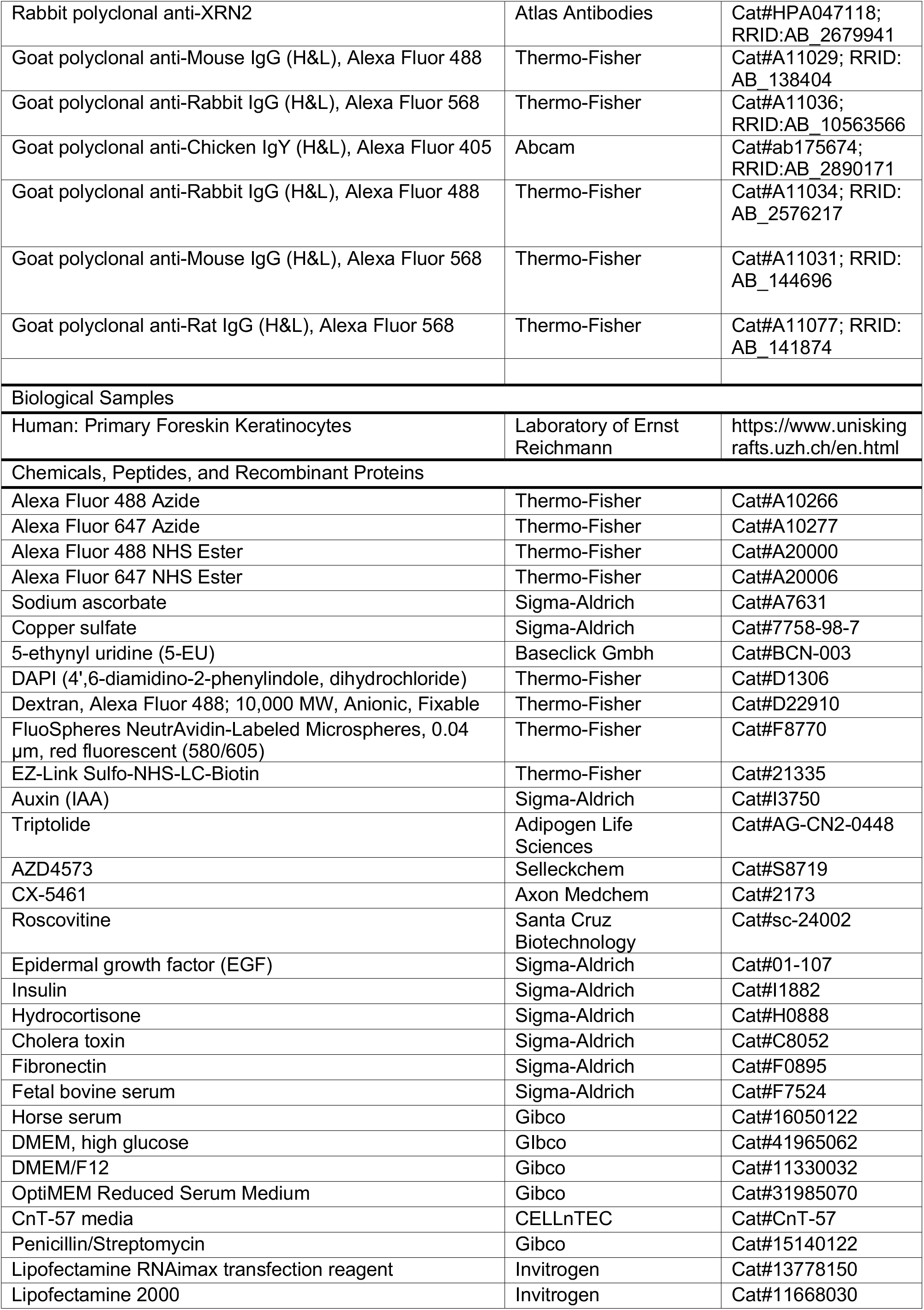
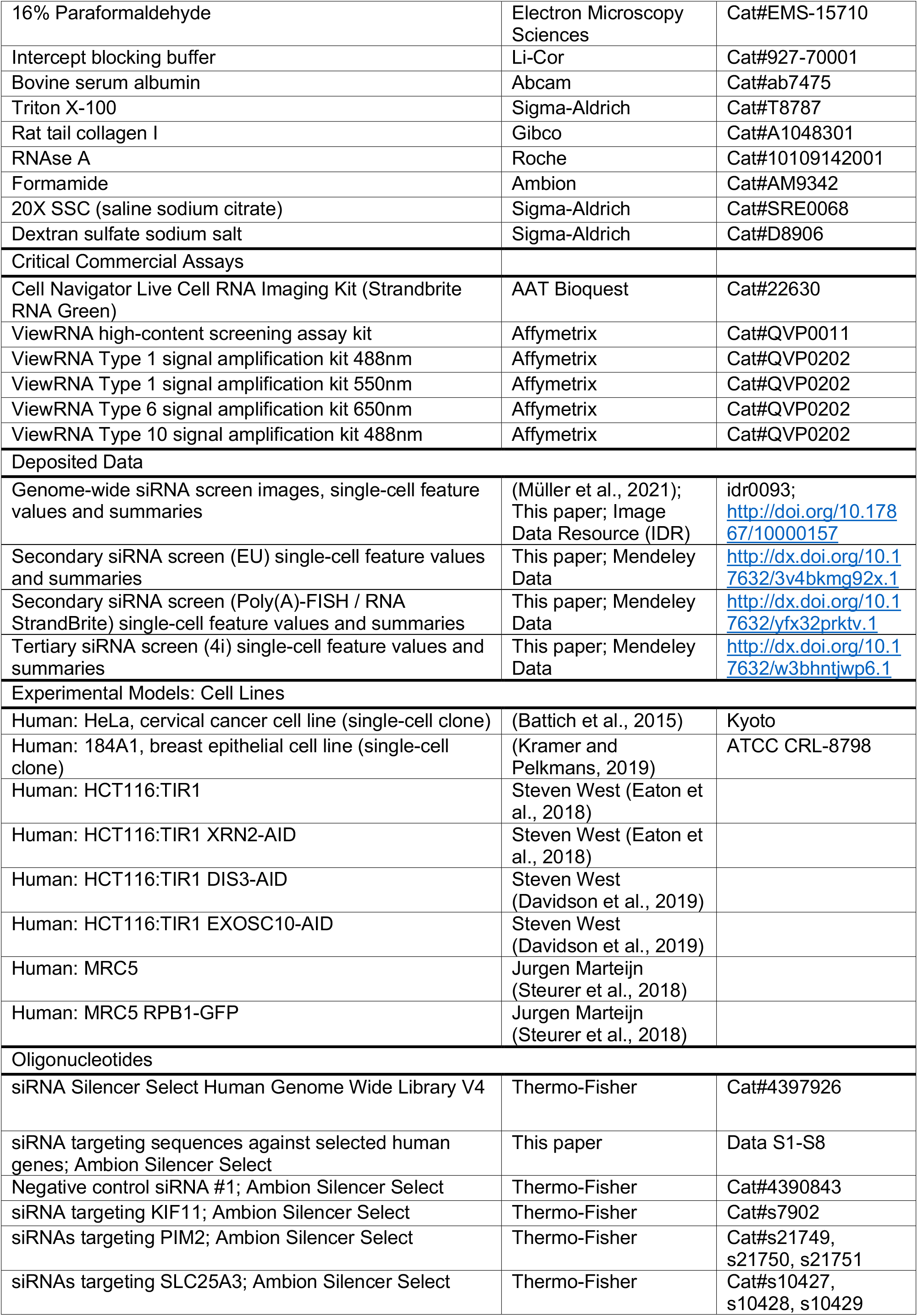
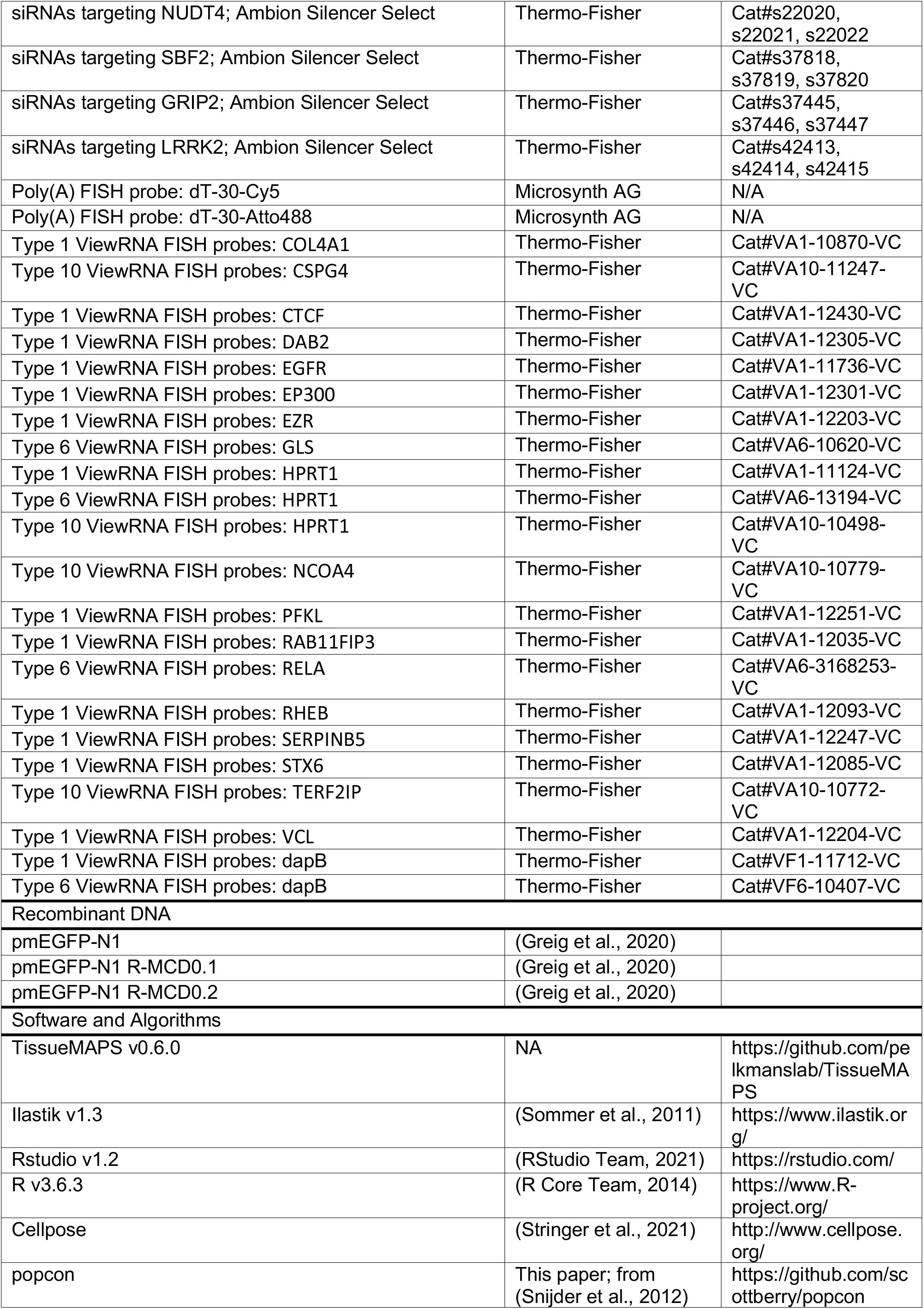
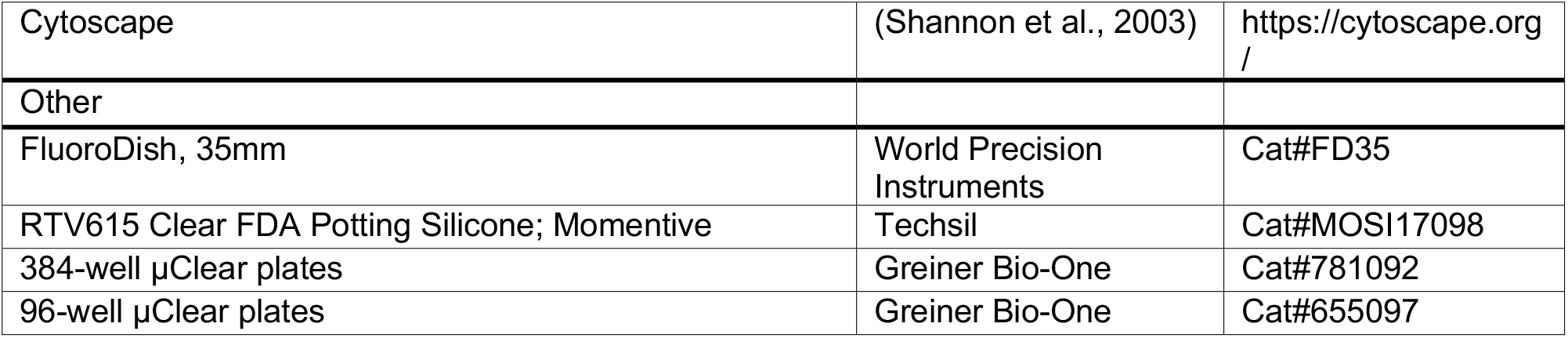
KEY RESOURCES.

