## Supplemental Figures and Tables for "Nuclear RNA concentration coordinates RNA production with cell size in human cells"

### Figure S1

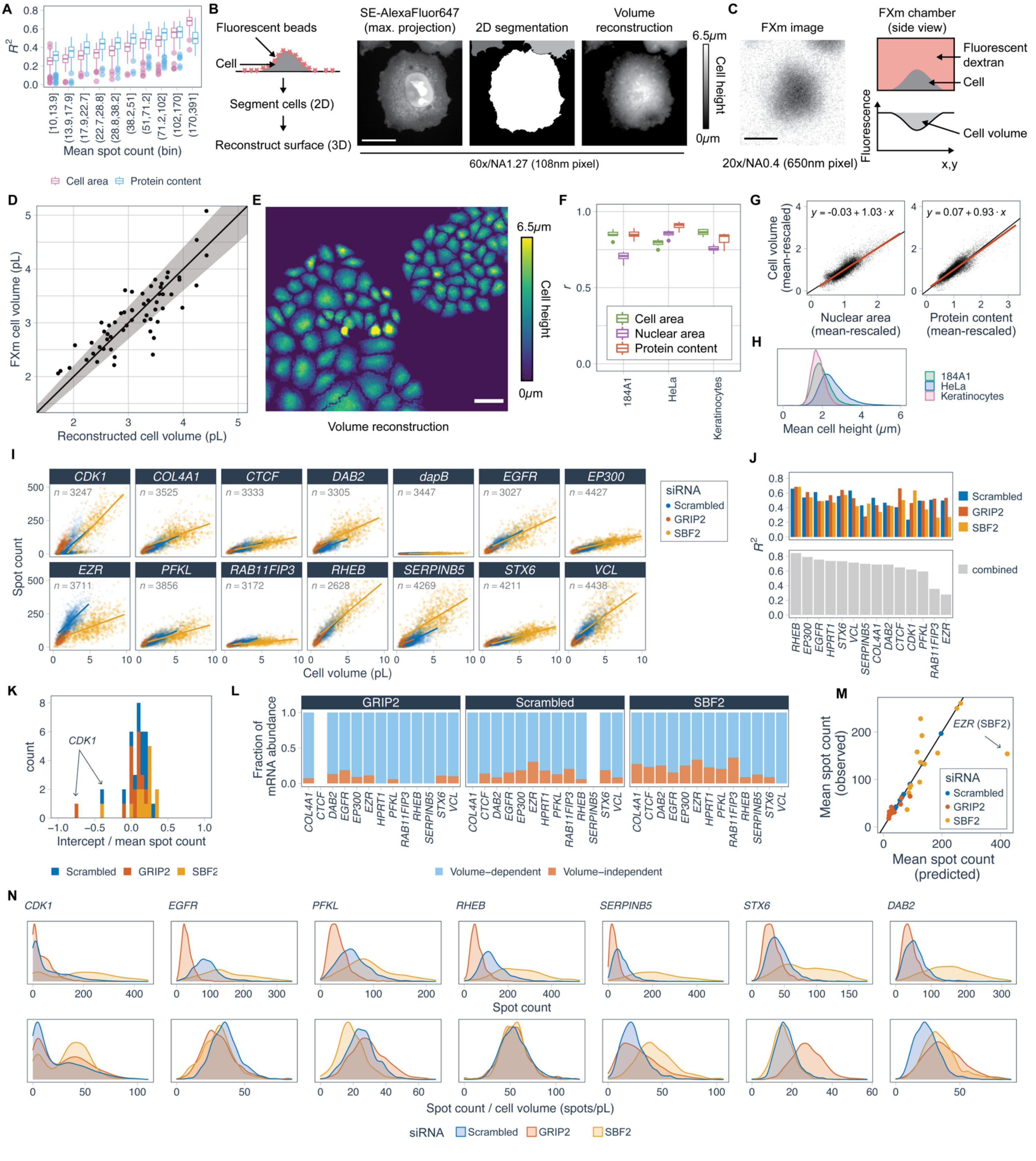

A
Mean spot count (bin)
Cell area
Protein content
B
Fluorescent beads
Cell
Segment cells (2D)
Reconstruct surface (3D)
SE-AlexaFluor647 (max. projection)
2D segmentation
Volume reconstruction
6.5μm
Cell height
0μm
60x/NA1.27 (108nm pixel)
C
FXm image
20x/NA0.4 (650nm pixel)
FXm chamber (side view)
Fluorescent dextran
Cell
Fluorescence
Cell volume
x,y
D
FXm cell volume (pL)
Reconstructed cell volume (pL)
E
Volume reconstruction
F
Cell area
Nuclear area
Protein content
184A1
HeLa
Keratinocytes
G
y = -0.03 + 1.03 · x
y = 0.07 + 0.93 · x
Cell volume (mean-rescaled)
Nuclear area (mean-rescaled)
Protein content (mean-rescaled)
H
Mean cell height (μm)
184A1
HeLa
Keratinocytes
I
CDK1
COL4A1
CTCF
DAB2
dapB
EGFR
EP300
EZR
PFKL
RAB11FIP3
RHEB
SERPINB5
STX6
VCL
n = 3247
n = 3525
n = 3333
n = 3305
n = 3447
n = 3027
n = 4427
n = 3711
n = 3856
n = 3172
n = 2628
n = 4269
n = 4211
n = 4438
Spot count
Cell volume (pL)
siRNA
Scrambled
GRIP2
SBF2
J
Scrambled
GRIP2
SBF2
combined
K
CDK1
count
Intercept / mean spot count
Scrambled
GRIP2
SBF2
L
GRIP2
Scrambled
SBF2
Fraction of mRNA abundance
Volume−dependent
Volume−independent
M
EZR (SBF2)
Mean spot count (observed)
Mean spot count (predicted)
siRNA
Scrambled
GRIP2
SBF2
N
CDK1
EGFR
PFKL
RHEB
SERPINB5
STX6
DAB2
Spot count
Spot count / cell volume (spots/pL)
siRNA
Scrambled
GRIP2
SBF2

**Figure S1, related to Figure 1: Cell volume measurement and smFISH. (A)** Coefficient of determination ($R^2$) for linear regression predicting mRNA spot count at the single-cell level from protein content or cell area in HeLa cells for 505 genes (all genes with mean spot count > 10 from Battich et al., 2015). **(B)** Cell volume reconstruction is achieved by attaching fluorescent beads to the cell and slide surface. Cells are then imaged as confocal z-stacks and segmented in x-y from maximum projected images of total protein stain (succinimidyl ester). The position of beads relative to the slide surface in three dimensions is then determined from confocal stacks. Beads are then interpolated to reconstruct the cell volume. **(C)** The same cell shown in B, imaged using the fluorescence exclusion method (FXm), as depicted alongside (Cadart et al., 2017). **(D)** Comparison of cell volume in HeLa cells measured by bead-based reconstruction and FXm (n=58). Line shows 1:1 correspondence. Shaded region represents the ±10% error estimated from FXm. **(E)** Cell height above the slide surface measured in high-throughput for a population of HeLa cells. Scale bar 50μm. **(F)** Correlations between high-throughput cell volume measurement and other 'cell size' features obtained from maximum-projected confocal images. Boxplots summarise correlations calculated in different wells (n=13,12,14 wells for 184A1, HeLa, and keratinocytes, respectively). **(G)** Relationship between cell volume and nuclear area or total cellular protein content in HeLa cells. Points are single cells, with all axes rescaled by means. Black line shows 1:1 correspondence. Red line shows linear fit, as specified by the inset equation (n=17,000 cells). **(H)** Mean cell height distributions obtained from 3D reconstructions. (n=5,951; 17,000; or 10,635 cells for 184A1, HeLa, and keratinocytes, respectively). **(I)** Spot count: number of cytoplasmic transcripts detected by bDNA smFISH for the genes indicated, as a function of cell volume, in cells transfected with scrambled siRNA or siRNA targeting GRIP2 or SBF2. *dapB* is bacterial gene used here as a negative control for smFISH. Fit lines show the regression $n_{\text{spots}} = a + bV$. Total cell number inset. **(J)** $R^2$ for linear regression predicting spot count from cell volume, for the genes indicated. Either considering genetic perturbations separately or combined. **(K)** Intercept values from regression for the 14 genes measured, normalised by mean spot count. **(L)** Fraction of predicted mRNA abundance that is volume independent ($a/(a + bV)$) or volume-dependent ($bV/(a + bV)$) for all genes measured (Padovan-Merhar et al., 2015). Because these comparisons are meaningful only for $a, b > 0$, we replaced 6 non-significant (p > 0.05) negative intercept values with zero, and omitted 5 gene/siRNA combinations with significant negative intercepts: CDK1 (all conditions), CTCF (GRIP2 siRNA) and SERPINB5 (Scrambled siRNA). **(M)** Mean spot count predicted from the change in cell volume in perturbations ($\bar{n}_{\text{perturbed}} = \bar{n}_{\text{scrambled}} \bar{V}_{\text{perturbed}} / \bar{V}_{\text{scrambled}}$). Line shows the fit through all scrambled siRNA points. **(N)** Examples of single-cell spot count distribution for genes indicated above, normalised by cell volume in lower panels.

#### Figure S2

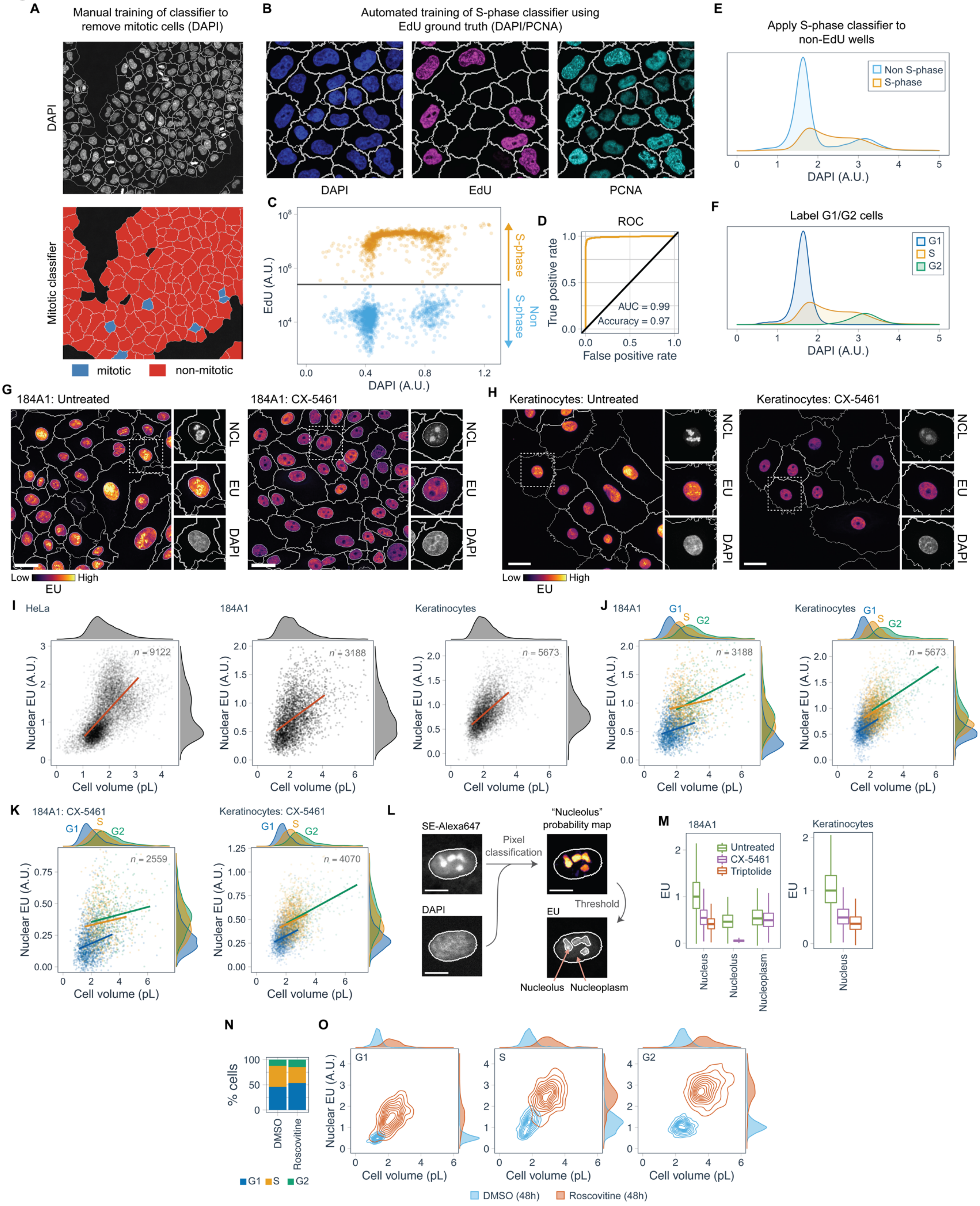
A
Manual training of classifier to remove mitotic cells (DAPI)
DAPI
Mitotic classifier
mitotic
non-mitotic
B
Automated training of S-phase classifier using EdU ground truth (DAPI/PCNA)
DAPI
EdU
PCNA
C
EdU (A.U.)
DAPI (A.U.)
S-phase
Non S-phase
D
ROC
True positive rate
False positive rate
AUC = 0.99
Accuracy = 0.97
E
Apply S-phase classifier to non-EdU wells
Non S-phase
S-phase
DAPI (A.U.)
F
Label G1/G2 cells
G1
S
G2
DAPI (A.U.)
G
184A1: Untreated
184A1: CX-5461
NCL
EU
DAPI
Low
High
EU
H
Keratinocytes: Untreated
Keratinocytes: CX-5461
NCL
EU
DAPI
Low
High
EU
I
HeLa
184A1
Keratinocytes
n = 9122
n = 3188
n = 5673
Nuclear EU (A.U.)
Cell volume (pL)
J
184A1
Keratinocytes
G1
S
G2
n = 3188
n = 5673
Nuclear EU (A.U.)
Cell volume (pL)
K
184A1: CX-5461
Keratinocytes: CX-5461
G1
S
G2
n = 2559
n = 4070
Nuclear EU (A.U.)
Cell volume (pL)
L
SE-Alexa647
DAPI
Pixel classification
"Nucleolus" probability map
Threshold
EU
Nucleolus
Nucleoplasm
M
184A1
Keratinocytes
Untreated
CX-5461
Triptolide
EU
Nucleus
Nucleolus
Nucleoplasm
N
% cells
DMSO
Roscovitine
G1
S
G2
O
G1
S
G2
Nuclear EU (A.U.)
Cell volume (pL)
DMSO (48h)
Roscovitine (48h)

**Figure S2, related to Figure 1: Cell-cycle classification and nascent RNA measurements. (A)** Manual supervised classification of mitotic cells in TissueMAPS, using texture features derived from DAPI staining (STAR Methods). **(B)** For cells treated with 30 min 5-ethynyl-2'-deoxyuridine (EdU) pulse, incorporation of label indicates active DNA replication. PCNA texture features also show similar characteristic staining in S-phase cells. **(C)** A random-forest classifier was trained using features derived from DAPI and PCNA to predict S-phase cells in EdU-treated wells, using EdU incorporation as a ground truth. **(D)** Receiver operating characteristic (ROC) curve for the classifier, derived from non-training data. Inset values show area under the curve (AUC) and accuracy of the final classifier. **(E)** The trained classifier is applied to non-EdU treated wells to identify S-phase cells using PCNA and DAPI measurements. **(F)** G1 and G2 cells are distinguished using the central local minimum of sum nuclear DAPI intensity of non S-phase cells **(G,H)** EU incorporated into nascent RNA during 30 min incubation of untreated and CX-5461-treated 184A1 cells, and primary human keratinocytes. EU was visualised using click chemistry with fluorescent azide (STAR Methods). Inset panels show NCL immunofluorescence together with EU and DAPI staining. Scale bars 25μm. **(I)** Sum nuclear EU intensity as a function of cell volume for cell types indicated. EU and cell volume distributions are shown to the right and top, respectively. Total number of cells inset. **(J)** Sum nuclear EU intensity as a function of cell volume, separating cells by cell-cycle stage, for cell types indicated. EU and cell volume distributions are shown to the right and top, respectively. Linear fit lines truncated at the 5-95th percentile of the cell volume distribution in each cell cycle phase. Total number of cells inset. Same data as I. **(K)** As in J, for cells pre-incubated with CX-5461 for 2h. **(L)** Pixel classification strategy for segmentation of the nucleolus from succinimidyl ester (SE)-AlexaFluor647 and DAPI images (Müller et al., 2021). **(M)** Sum EU intensity measured in the nucleus, nucleolus and nucleoplasm (non-nucleolus), with cell type as indicated. Boxplots summarise single-cell values with outliers omitted for clarity. Values relative to median nuclear EU intensity of untreated cells. **(N)** Cell-cycle distribution and **(O)** sum nuclear EU and cell volume bivariate distributions in HeLa cells treated with roscovitine for 48h before EU metabolic labelling. EU and cell volume distributions are shown to the right and top, respectively.

### Figure S3

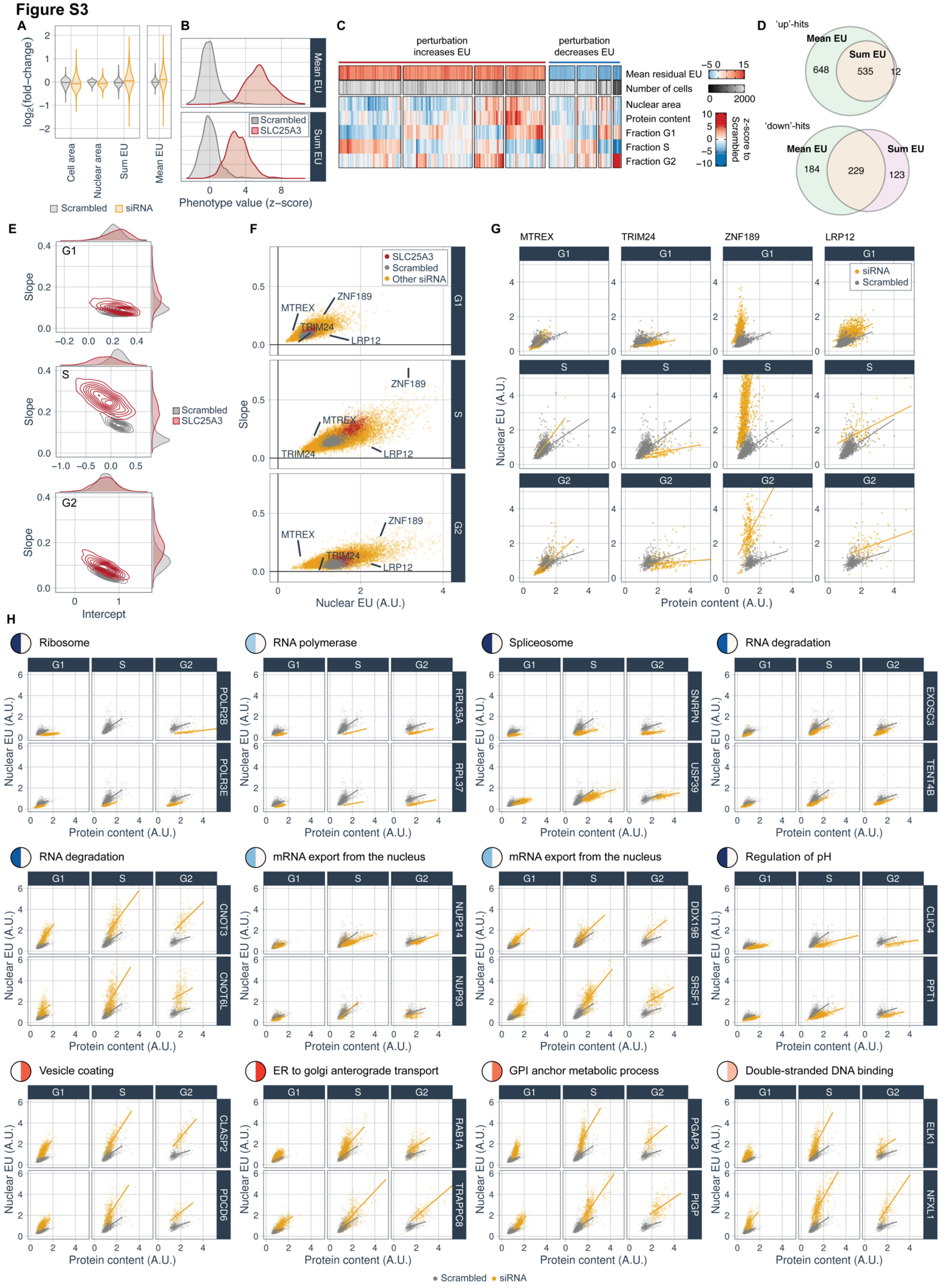

A
log2(fold-change)
Cell area
Nuclear area
Sum EU
Mean EU
Scrambled
siRNA
B
Mean EU
Sum EU
Scrambled
SLC25A3
Phenotype value (z-score)
C
perturbation increases EU
perturbation decreases EU
Mean residual EU
Number of cells
Nuclear area
Protein content
Fraction G1
Fraction S
Fraction G2
z-score to Scrambled
D
'up'-hits
Mean EU
Sum EU
648
535
12
'down'-hits
Mean EU
Sum EU
184
229
123
E
G1
S
G2
Slope
Intercept
Scrambled
SLC25A3
F
SLC25A3
Scrambled
Other siRNA
MTREX
TRIM24
ZNF189
LRP12
Slope
Nuclear EU (A.U.)
G
MTREX
TRIM24
ZNF189
LRP12
siRNA
Scrambled
Nuclear EU (A.U.)
Protein content (A.U.)
H
Ribosome
POLR2B
POLR3E
RNA polymerase
RPL35A
RPL37
Spliceosome
SNRPN
USP39
RNA degradation
EXOSC3
TENT4B
RNA degradation
CNOT3
CNOT6L
mRNA export from the nucleus
NUP214
NUP93
mRNA export from the nucleus
DDX19B
SRSF1
Regulation of pH
CLIC4
PPT1
Vesicle coating
CLASP2
PDCD6
ER to golgi anterograde transport
RAB1A
TRAPPC8
GPI anchor metabolic process
PGAP3
PIGP
Double-stranded DNA binding
ELK1
NFXL1
Nuclear EU (A.U.)
Protein content (A.U.)
Scrambled
siRNA

**Figure S3, related to Figure 2: Genome-wide screen. (A)** Variability in perturbation-averaged cellular feature values in the genome-wide screen, quantified as the $\log_2$(fold-change) compared to scrambled siRNA controls. Violin plots summarise conditions with at least 500 cells. **(B)** Distributions of residual mean EU and residual sum EU across all scrambled siRNA and SLC25A3 control wells of the genome-wide screen. **(C)** Hierarchical clustering of perturbation-average cell size features and cell cycle fractions for mean residual EU hits. Number of cells and mean residual EU are annotations only and were not used for clustering. Values z-scored to scrambled siRNA controls. **(D)** Euler plots for numbers of genes annotated as hits ($p_{\text{posterior}} > 0.85$) using residual mean EU and residual sum EU (STAR Methods). **(E)** Bivariate distributions of slope and intercept values obtained from cell-cycle-specific regression of sum nuclear EU from total protein content in control wells of the genome-wide screen (STAR Methods). Panels show different cell cycle stages. **(F)** Slope and intercept values obtained from cell-cycle-specific regression of sum nuclear EU from total protein content, for all wells in the genome-wide screen. Panels show different cell cycle stages. Selected perturbations highlighted in G. **(G)** Examples of the relationship between sum nuclear EU and total protein content for example genetic perturbations highlighted in F. siRNA shown above. A single exemplary scrambled siRNA well included on each panel for comparison. **(H)** As in G, for example siRNA perturbations corresponding to enriched annotations in Figure 2F. siRNA shown on right. Enriched annotation shown above, with node colours as in Figure 2F.

**Figure S4**

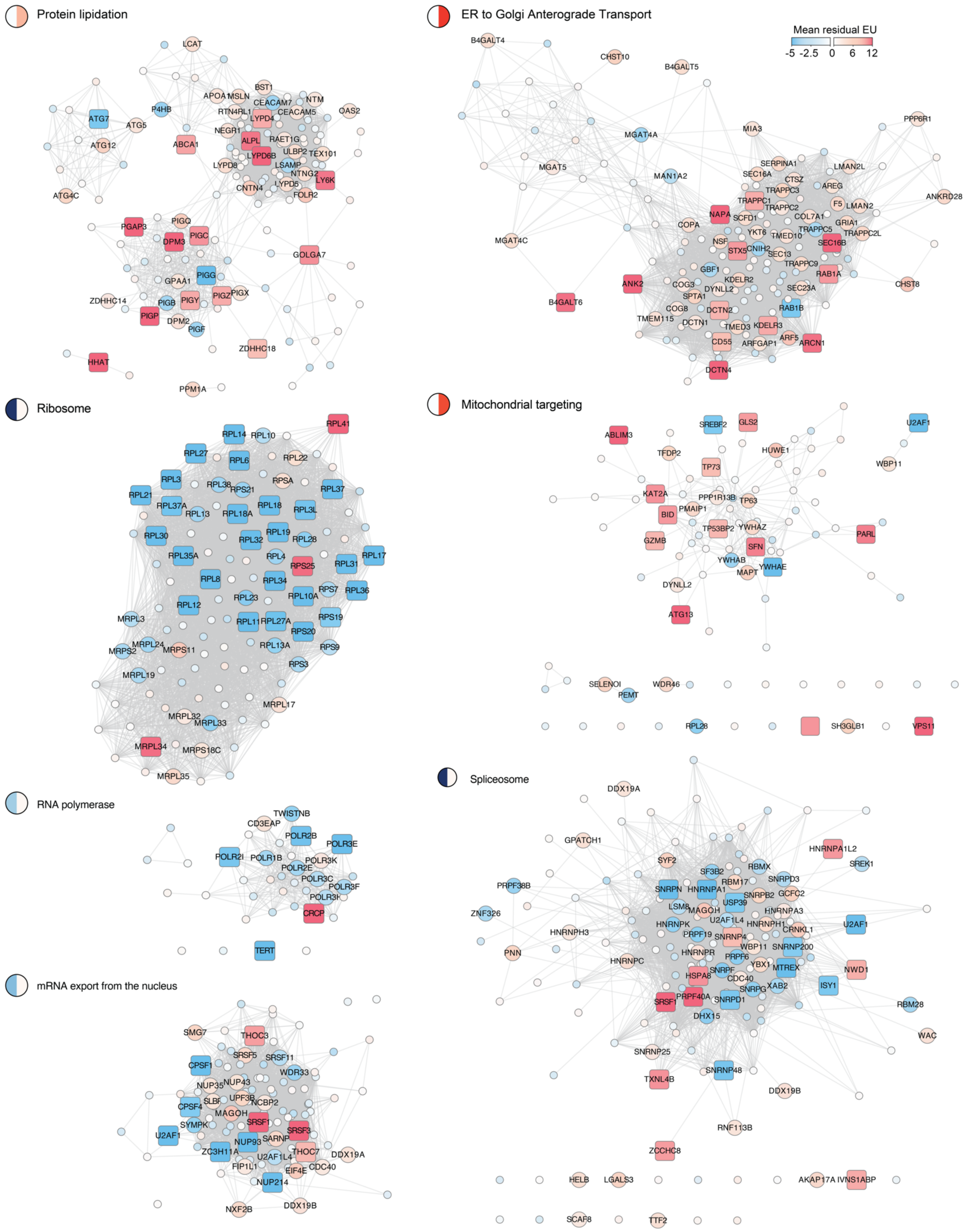

Protein lipidation
LCAT
BST1
APOA1
MSLN
CEACAM7
NTM
P4HB
RTN4RL1
LYPD4
CEACAM5
OAS2
ATG7
ATG5
NEGR1
RAET1G
ATG12
ABCA1
ALPL
ULBP2
TEX101
LYPD6B
LYPD8
LSAMP
NTNG2
LY6K
CNTN4
LYPD5
FOLR2
ATG4C
PGAP3
PIGQ
PIGC
DPM3
GOLGA7
PIGG
GPAA1
PIGZ
PIGX
ZDHHC14
PIGB
PIGY
PIGP
DPM2
PIGF
ZDHHC18
HHAT
PPM1A
ER to Golgi Anterograde Transport
Mean residual EU
-5 -2.5 0 6 12
B4GALT4
CHST10
B4GALT5
MIA3
PPP6R1
MGAT4A
SERPINA1
LMAN2L
MGAT5
SEC16A
MAN1A2
CTSZ
TRAPPC3
AREG
ANKRD28
TRAPPC1
F5
LMAN2
TRAPPC2
NAPA
SCFD1
COL7A1
COPA
GRIA1
YKT6
TRAPPC5
TMED10
TRAPPC2L
MGAT4C
NSF
SEC16B
STX5
CNIH2
SEC13
TRAPPC9
GBF1
RAB1A
CHST8
ANK2
KDELR2
COG3
DYNLL2
SEC23A
SPTA1
B4GALT6
COG8
DCTN2
RAB1B
TMEM115
DCTN1
TMED3
KDELR3
CD55
ARF5
ARFGAP1
ARCN1
DCTN4
Ribosome
RPL41
RPL14
RPL10
RPL27
RPL6
RPL22
RPL3
RPL38
RPS21
RPSA
RPL37
RPL21
RPL37A
RPL18
RPL13
RPL18A
RPL3L
RPL19
RPL30
RPL32
RPL28
RPL17
RPL35A
RPL4
RPS25
RPL31
RPL8
RPL34
RPS7
RPL36
RPL23
RPL10A
RPL12
MRPL3
RPS19
RPL11
RPL27A
RPS20
MRPL24
RPL13A
MRPS2
MRPS11
RPS9
RPS3
MRPL19
MRPL17
MRPL32
MRPL33
MRPL34
MRPS18C
MRPL35
Mitochondrial targeting
SREBF2
GLS2
U2AF1
ABLIM3
TFDP2
HUWE1
TP73
WBP11
KAT2A
PPP1R13B
TP63
BID
PMAIP1
TP53BP2
YWHAZ
PARL
GZMB
SFN
YWHAB
YWHAE
MAPT
DYNLL2
ATG13
SELENOI
WDR46
PEMT
RPL28
SH3GLB1
VPS11
RNA polymerase
TWISTNB
CD3EAP
POLR2B
POLR3E
POLR2I
POLR1B
POLR3K
POLR2E
POLR3C
POLR3F
POLR3H
CRCP
TERT
mRNA export from the nucleus
SMG7
THOC3
SRSF5
SRSF11
CPSF1
WDR33
NUP35
NUP43
SLBP
UPF3B
NCBP2
CPSF4
MAGOH
SYMPK
SRSF1
SRSF3
U2AF1
NUP93
SARNP
ZC3H11A
U2AF1L4
THOC7
DDX19A
FIP1L1
EIF4E
CDC40
NUP214
NXF2B
DDX19B
Spliceosome
DDX19A
GPATCH1
HNRNPA1L2
SYF2
SREK1
SF3B2
RBMX
PRPF38B
RBM17
SNRPD3
SNRPN
HNRNPA1
SNRPB2
GCFC2
USP39
LSM8
MAGOH
HNRNPA3
ZNF326
U2AF1L4
HNRNPK
HNRNPH1
CRNKL1
U2AF1
HNRNPH3
PRPF19
SNRNP4
WBP11
SNRNP200
PNN
HNRNPR
PRPF6
HNRNPC
YBX1
MTREX
NWD1
HSPA8
SNRPF
CDC40
SNRPG
XAB2
ISY1
SRSF1
PRPF40A
SNRPD1
RBM28
DHX15
WAC
SNRNP25
SNRNP48
TXNL4B
DDX19B
RNF113B
ZCCHC8
HELB
LGALS3
AKAP17A
IVNS1ABP
SCAF8
TTF2

**Figure S4, related to Figure 2: Genome-wide screen.** STRING protein-protein association network of genes with selected enriched annotations, omitting perturbations with less than 500 cells. Labelled circles for $p_{\text{posterior}} > 0.5$ and squares for $p_{\text{posterior}} > 0.85$. Network node colours represent mean residual EU, as shown in legend (top-left). Two-colour nodes in titles as in Figure 2F.

**Figure S5**

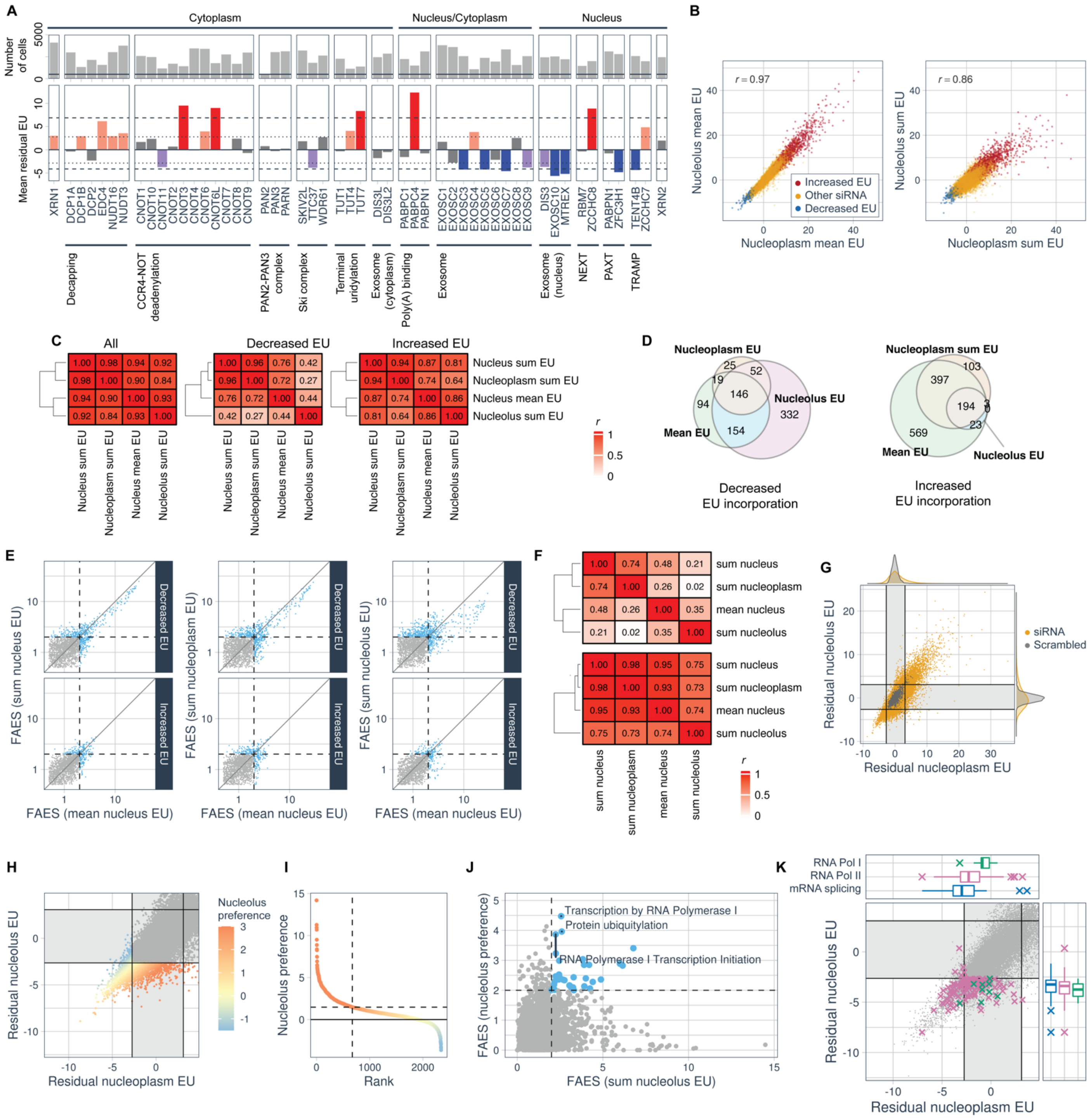
A
Cytoplasm
Nucleus/Cytoplasm
Nucleus
Number of cells
Mean residual EU
XRN1
DCP1A
DCP1B
DCP2
EDC4
NUDT16
NUDT3
CNOT1
CNOT10
CNOT11
CNOT2
CNOT3
CNOT4
CNOT6
CNOT6L
CNOT7
CNOT8
CNOT9
PAN2
PAN3
PARN
SKIV2L
TTC37
WDR61
TUT1
TUT4
TUT7
DIS3L
DIS3L2
PABPC1
PABPC4
PABPN1
EXOSC1
EXOSC2
EXOSC3
EXOSC4
EXOSC5
EXOSC6
EXOSC7
EXOSC8
EXOSC9
DIS3
EXOSC10
MTREX
RBM7
ZCCHC8
PABPN1
ZFC3H1
TENT4B
ZCCHC7
XRN2
Decapping
CCR4-NOT deadenylation
PAN2-PAN3 complex
Ski complex
Terminal uridylation
Exosome (cytoplasm)
Poly(A) binding
Exosome
Exosome (nucleus)
NEXT
PAXT
TRAMP
B
r = 0.97
r = 0.86
Increased EU
Other siRNA
Decreased EU
Nucleolus mean EU
Nucleoplasm mean EU
Nucleolus sum EU
Nucleoplasm sum EU
C
All
Decreased EU
Increased EU
Nucleus sum EU
Nucleoplasm sum EU
Nucleus mean EU
Nucleolus sum EU
r
D
Nucleoplasm EU
Nucleolus EU
Mean EU
Decreased EU incorporation
Nucleoplasm sum EU
Nucleolus EU
Mean EU
Increased EU incorporation
E
FAES (sum nucleus EU)
FAES (sum nucleoplasm EU)
FAES (sum nucleolus EU)
FAES (mean nucleus EU)
Decreased EU
Increased EU
F
sum nucleus
sum nucleoplasm
mean nucleus
sum nucleolus
G
siRNA
Scrambled
Residual nucleolus EU
Residual nucleoplasm EU
H
Nucleolus preference
Residual nucleolus EU
Residual nucleoplasm EU
I
Nucleolus preference
Rank
J
Transcription by RNA Polymerase I
Protein ubiquitylation
RNA Polymerase I Transcription Initiation
FAES (nucleolus preference)
FAES (sum nucleolus EU)
K
RNA Pol I
RNA Pol II
mRNA splicing
Residual nucleolus EU
Residual nucleoplasm EU

**Figure S5, related to Figure 2: Genome-wide screen supplements. (A)** Mean residual EU and cell number for a manually-selected group of genes with known roles in RNA degradation. Genes were assigned to a single complex or pathway and then to the nucleus, cytoplasm, or both based on recent literature reviews (Grudzien-Nogalska and Kiledjian, 2017; Łabno et al., 2016; Schmid and Jensen, 2018; Siwaszek et al., 2014). Dotted and dashed lines, as well as bar colours, indicate lower and upper hit thresholds ($p_{\text{posterior}} = 0.5, 0.85$), respectively. **(B)** Correlations of perturbation-averaged nucleoplasm and nucleolar EU measurements (mean or sum indicates whether pixel values in each subnuclear compartment are averaged or summed). Colours indicate perturbations identified as 'mean EU' hits ($p_{\text{posterior}} > 0.85$). Perturbations with low cell number omitted. Values z-scored to scrambled siRNA control wells. **(C)** Pearson's correlations between residual EU phenotypes. Three panels represent correlations calculated over different sets of perturbations, either all, or only those with $p_{\text{posterior (decreased EU)}} > 0.5$ or $p_{\text{posterior (increased EU)}} > 0.5$, for at least one of the four residual EU phenotypes. **(D)** Euler plots showing the number of genes annotated as hits ($p_{\text{posterior}} > 0.85$) using residual sum nucleolar EU, sum nucleoplasmic EU or mean nuclear EU. **(E)** Scatter plots comparing functional annotation enrichment scores (FAES) derived from residual sum EU phenotypes, compared to those for residual mean EU. Dashed lines show FAES = 2 **(F)** Correlations between FAES scores of annotations, calculated for annotations that have FAES > 2 for at least one of the phenotypes being compared. Only annotations with 30-3500 genes considered, with groups of similar annotations combined when Cohen's $\kappa > 0.85$ (STAR Methods). Top/bottom panels for increased/decreased EU. **(G)** Scatter plot comparing residual nucleoplasmic and nucleolar EU for all perturbations. Shaded grey boxes indicate $p_{\text{posterior}} < 0.5$. **(H)** Zoom in of G, showing perturbations with nucleolar-specific EU reduction, with points coloured by the 'nucleolar preference' (residual of regression of y from x). **(I)** Genetic perturbations with $p_{\text{posterior}} > 0.5$ for either reduced residual sum nucleolus EU or reduced residual sum nucleoplasm EU. Ranked and coloured by nucleolus preference. Dashed line indicates the 1st percentile of scrambled siRNA control wells, and the corresponding rank. **(J)** FAES for 'increased nucleolar preference' versus FAES for 'reduced residual sum nucleolus EU'. Blue points indicate annotations enriched in both analyses. Top 3 enriched annotations highlighted. **(K)** Residual nucleoplasm sum EU versus residual nucleolar sum EU, omitting perturbations with low cell number. siRNA perturbations of genes with annotations related to RNA Pol I, RNA Pol II, or mRNA splicing. Shaded grey rectangles indicate $p_{\text{posterior}} < 0.5$, for each phenotype.

Figure S6

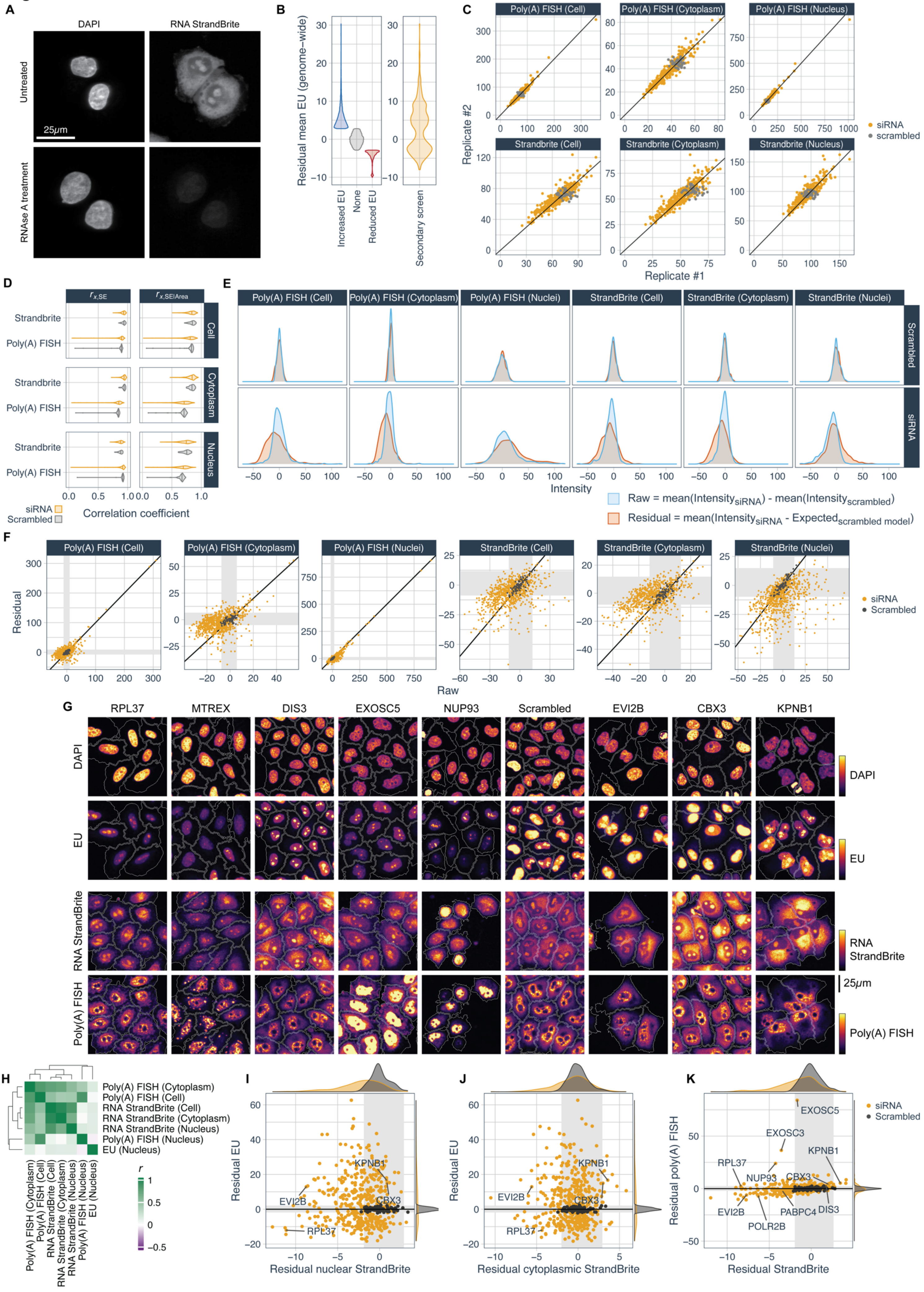
A
DAPI
RNA StrandBrite
Untreated
RNAse A treatment
25μm
B
Residual mean EU (genome-wide)
Increased EU
None
Reduced EU
Secondary screen
C
Poly(A) FISH (Cell)
Poly(A) FISH (Cytoplasm)
Poly(A) FISH (Nucleus)
Strandbrite (Cell)
Strandbrite (Cytoplasm)
Strandbrite (Nucleus)
Replicate #2
Replicate #1
siRNA
scrambled
D
Strandbrite
Poly(A) FISH
Cell
Cytoplasm
Nucleus
siRNA
Scrambled
Correlation coefficient
E
Poly(A) FISH (Cell)
Poly(A) FISH (Cytoplasm)
Poly(A) FISH (Nuclei)
StrandBrite (Cell)
StrandBrite (Cytoplasm)
StrandBrite (Nuclei)
Scrambled
siRNA
Intensity
Raw = mean(Intensity siRNA) - mean(Intensity scrambled)
Residual = mean(Intensity siRNA - Expected scrambled model)
F
Poly(A) FISH (Cell)
Poly(A) FISH (Cytoplasm)
Poly(A) FISH (Nuclei)
StrandBrite (Cell)
StrandBrite (Cytoplasm)
StrandBrite (Nuclei)
Residual
Raw
siRNA
Scrambled
G
RPL37
MTREX
DIS3
EXOSC5
NUP93
Scrambled
EVI2B
CBX3
KPNB1
DAPI
EU
RNA StrandBrite
Poly(A) FISH
25μm
H
Poly(A) FISH (Cytoplasm)
Poly(A) FISH (Cell)
RNA StrandBrite (Cell)
RNA StrandBrite (Cytoplasm)
RNA StrandBrite (Nucleus)
Poly(A) FISH (Nucleus)
EU (Nucleus)
r
I
Residual EU
Residual nuclear StrandBrite
KPNB1
EVI2B
CBX3
RPL37
J
Residual EU
Residual cytoplasmic StrandBrite
KPNB1
EVI2B
CBX3
RPL37
K
Residual poly(A) FISH
Residual StrandBrite
EXOSC5
EXOSC3
KPNB1
RPL37
NUP93
CBX3
EVI2B
PABPC4
DIS3
POLR2B
siRNA
Scrambled

**Figure S6, related to Figure 3: RNA abundance measurements. (A)** HeLa cells stained with DAPI and RNA StrandBrite with or without incubation with 0.5mg/mL RNAse A at 37°C for 30 min, together with untreated controls. Identical image rescaling for both samples. **(B)** Violin plots of mean residual EU from genome-wide screen, separated into increased/reduced EU incorporation classes according to $p_{\text{posterior}} > 0.50$ (STAR Methods) compared to violin plot of mean residual EU from the genome-wide screen for the 436 perturbations selected for the secondary screen. **(C)** Reproducibility of well-mean of mean intensity measurements for poly(A) FISH and RNA Strandbrite. Samples show 423 pairs of siRNA replicates together with 80 pairs of scrambled siRNA replicates (randomly paired). **(D)** Correlations of RNA Strandbrite and poly(A) FISH with protein content (succinimidyl ester, SE). $r_{x,\text{SE}}$ indicates (full) correlation of RNA Strandbrite or poly(A) FISH intensity for the nucleus, cytoplasm, or whole cell. $r_{x,\text{SE}|\text{Area}}$indicates a partial correlation, accounting for area of the object (nucleus, cytoplasm or whole cell). Violin plots summarise values calculated for each well. **(E)** Distributions of 'raw' and 'residual' mean RNA abundance measurements. 'Residual' is the well-mean of corrected single-cell intensity values, obtained by subtracting the intensity predicted from a model trained on scrambled siRNA controls (STAR Methods). 'Raw' is the well-mean of single-cell intensity values, with the distribution centred by subtracting the mean of scrambled wells. **(F)** Comparison of well-mean 'raw' and 'residual' measurements of RNA abundance, as defined in E. Grey boxes show the 1st/99th percentiles of scrambled siRNA controls wells. **(G)** Images of DAPI, EU (from secondary EU-metabolic labelling screen) and for RNA StrandBrite and poly(A) FISH (from secondary RNA abundance screen), for perturbations shown above, and labelled in I-K. **(H)** Pairwise correlations of 'raw' mean RNA abundance measurements and 'raw' mean EU across perturbations. See Figure 3D for similar plot for 'residual' measurements. **(I)** Mean residual EU versus mean nuclear RNA Strandbrite. Mean of two replicates shown for each siRNA. Grey boxes indicate 1st/99th percentiles of scrambled siRNA controls. **(J)** As in I, for cytoplasmic RNA Strandbrite. **(K)** As in I, for mean residual cellular poly(A) FISH versus mean residual cellular RNA Strandbrite.

### Figure S7

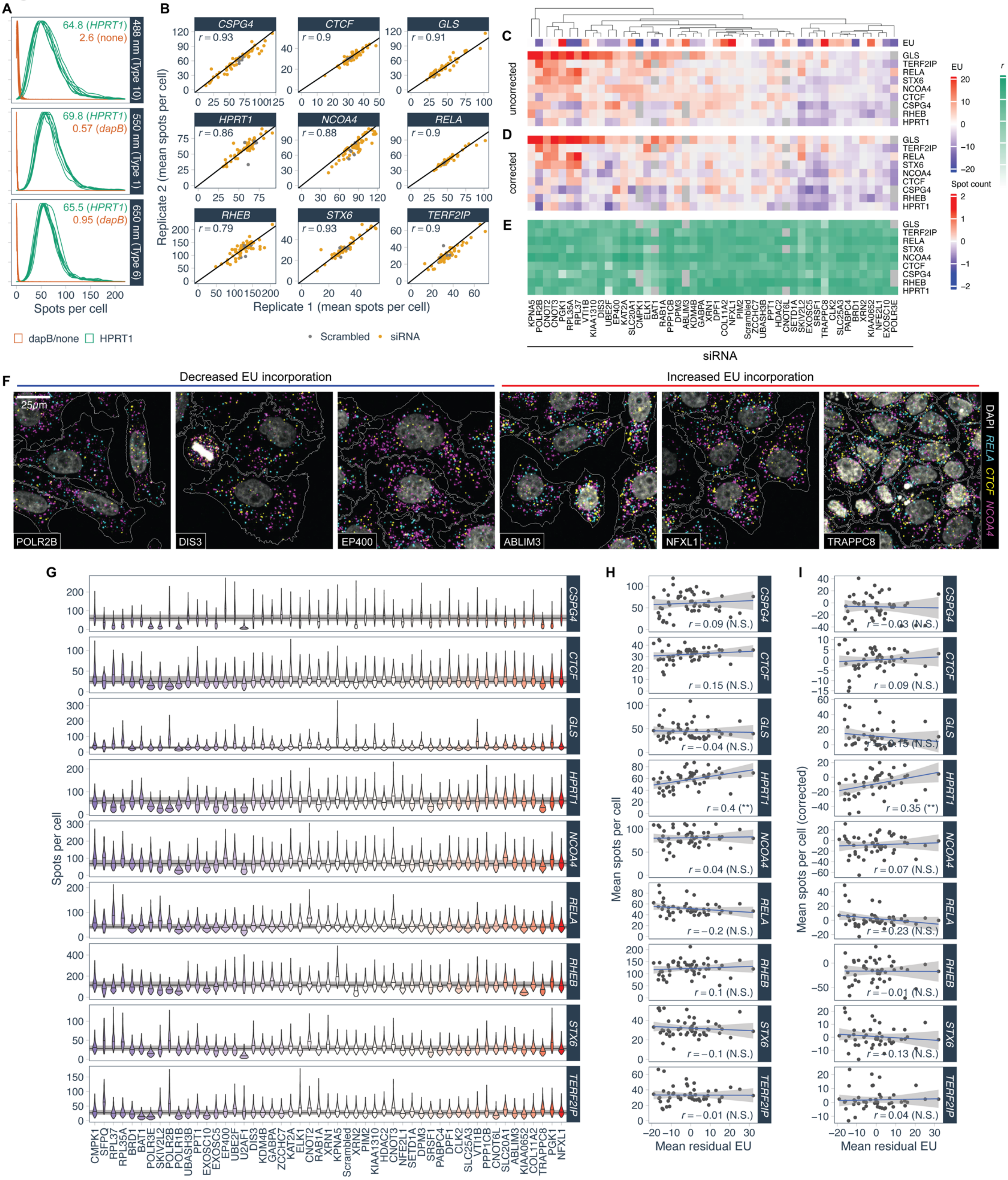

A
64.8 (HPRT1)
2.6 (none)
488 nm (Type 10)
69.8 (HPRT1)
0.57 (dapB)
550 nm (Type 1)
65.5 (HPRT1)
0.95 (dapB)
650 nm (Type 6)
Spots per cell
dapB/none
HPRT1
B
CSPG4
CTCF
GLS
HPRT1
NCOA4
RELA
RHEB
STX6
TERF2IP
r = 0.93
r = 0.9
r = 0.91
r = 0.86
r = 0.88
r = 0.9
r = 0.79
r = 0.93
r = 0.9
Replicate 2 (mean spots per cell)
Replicate 1 (mean spots per cell)
Scrambled
siRNA
C
D
E
EU
uncorrected
corrected
GLS
TERF2IP
RELA
STX6
NCOA4
CTCF
CSPG4
RHEB
HPRT1
Spot count
r
siRNA
F
Decreased EU incorporation
Increased EU incorporation
25μm
POLR2B
DIS3
EP400
ABLIM3
NFXL1
TRAPPC8
DAPI
RELA
CTCF
NCOA4
G
Spots per cell
H
Mean spots per cell
Mean residual EU
r = 0.09 (N.S.)
r = 0.15 (N.S.)
r = −0.04 (N.S.)
r = 0.4 (**)
r = 0.04 (N.S.)
r = −0.2 (N.S.)
r = 0.1 (N.S.)
r = −0.1 (N.S.)
r = −0.01 (N.S.)
I
Mean spots per cell (corrected)
Mean residual EU
r = −0.03 (N.S.)
r = 0.09 (N.S.)
r = −0.15 (N.S.)
r = 0.35 (**)
r = 0.07 (N.S.)
r = −0.23 (N.S.)
r = −0.01 (N.S.)
r = −0.13 (N.S.)
r = 0.04 (N.S.)

**Figure S7, related to Figure 3: single-molecule RNA FISH. (A)** Distributions of cytoplasmic bDNA smFISH spot counts for unperturbed cells for positive control (*HPRT1*) and negative control (*dapB* or none) obtained using bDNA FISH with different amplifier probe set types and excitation wavelengths, as shown to the right. Individual overlaid curves show the 4 replicate wells from each of the 2 plates. Inset values show mean spot counts, after pooling replicates. **(B)** Comparison of mean spot count per well between replicates, across all perturbations (n=49-51), and scrambled siRNA control wells (n=7). Correlations inset. Lines show 1:1 equality. **(C)** Hierarchical clustering of mean spot counts relative to scrambled siRNA controls $(\bar{n}_{\text{spots}} - \bar{n}_{\text{spots (scrambled)}})/\bar{n}_{\text{spots (scrambled)}}$ for all gene/perturbation combinations. Rows are transcripts measured. Columns are genes targeted by siRNA. EU annotation above is mean residual EU phenotype (mean of 3 replicates, Z-scored to scrambled). **(D)** As in C, except using transcript abundance corrected for cell size and cell cycle changes induced by perturbations (STAR methods). **(E)** Correlation coefficients of smFISH spots per cell with cellular protein content. **(F)** 3-plex smFISH images for three perturbations with reduced, or increased EU incorporation, respectively. Transcripts shown to the right. Gene targeted by siRNA inset. **(G)** Single-cell spot count distributions for each of the genes listed, across the panel of library siRNA perturbations shown underneath. Perturbations ordered and coloured according to residual mean EU. **(H)** Mean spot count per cell compared to mean residual EU. Pearson's correlation inset, with N.S. indicating not significant ($p > 0.05$), and ** indicating $p < 0.01$. **(I)** As in H, after correcting spot count measurements for cell size and cell cycle changes induced by perturbations (STAR methods).

#### Figure S8

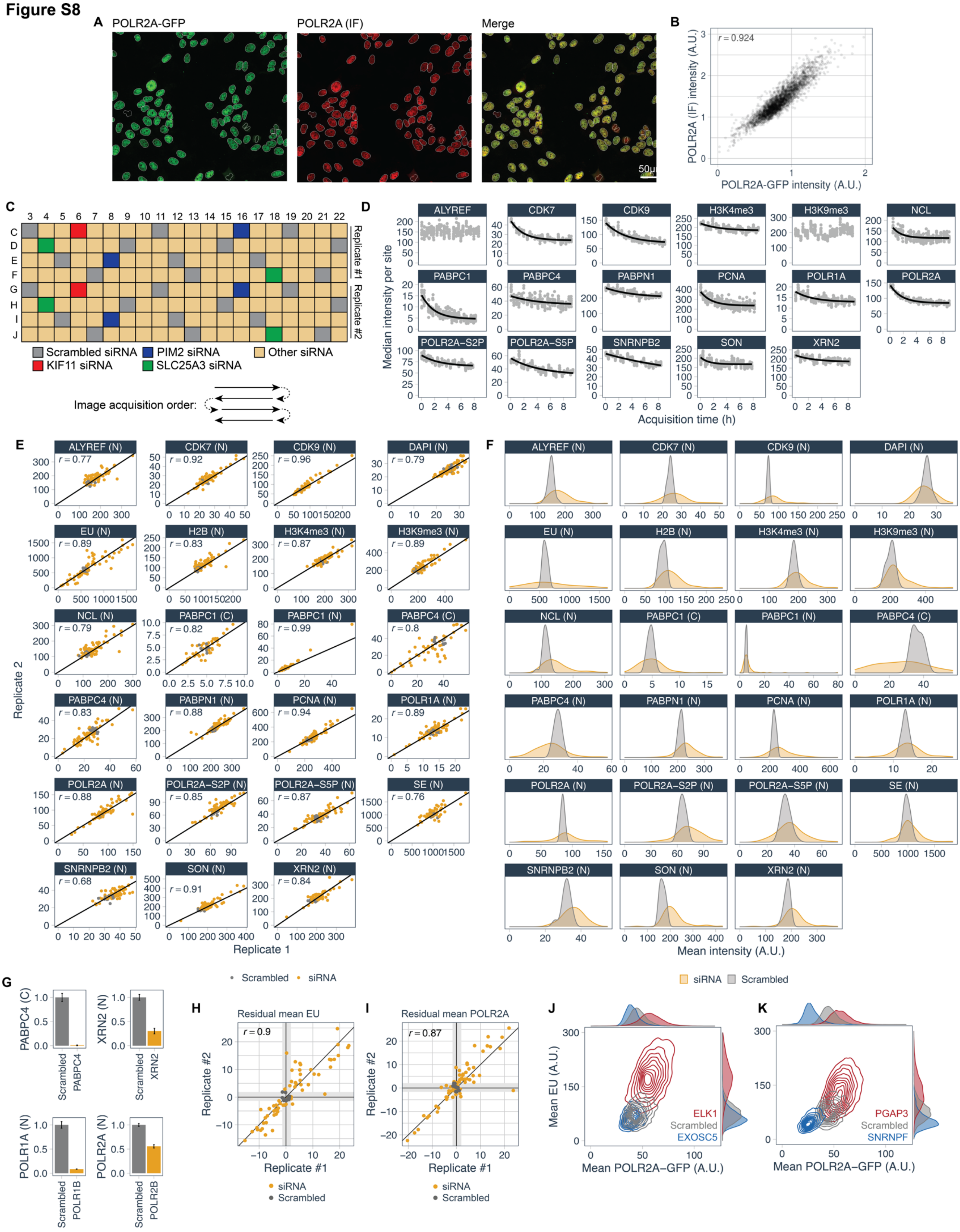

A
POLR2A-GFP
POLR2A (IF)
Merge
50µ
B
r = 0.924
POLR2A (IF) intensity (A.U.)
POLR2A-GFP intensity (A.U.)
C
Replicate #1
Replicate #2
Scrambled siRNA
PIM2 siRNA
Other siRNA
KIF11 siRNA
SLC25A3 siRNA
Image acquisition order:
D
Median intensity per site
Acquisition time (h)
ALYREF
CDK7
CDK9
H3K4me3
H3K9me3
NCL
PABPC1
PABPC4
PABPN1
PCNA
POLR1A
POLR2A
POLR2A-S2P
POLR2A-S5P
SNRNPB2
SON
XRN2
E
Replicate 2
Replicate 1
ALYREF (N) r = 0.77
CDK7 (N) r = 0.92
CDK9 (N) r = 0.96
DAPI (N) r = 0.79
EU (N) r = 0.89
H2B (N) r = 0.83
H3K4me3 (N) r = 0.87
H3K9me3 (N) r = 0.89
NCL (N) r = 0.79
PABPC1 (C) r = 0.82
PABPC1 (N) r = 0.99
PABPC4 (C) r = 0.8
PABPC4 (N) r = 0.83
PABPN1 (N) r = 0.88
PCNA (N) r = 0.94
POLR1A (N) r = 0.89
POLR2A (N) r = 0.88
POLR2A-S2P (N) r = 0.85
POLR2A-S5P (N) r = 0.87
SE (N) r = 0.76
SNRNPB2 (N) r = 0.68
SON (N) r = 0.91
XRN2 (N) r = 0.84
Scrambled
siRNA
F
Mean intensity (A.U.)
siRNA
Scrambled
G
PABPC4 (C)
XRN2 (N)
POLR1A (N)
POLR2A (N)
Scrambled
PABPC4
XRN2
POLR1B
POLR2B
H
Residual mean EU
r = 0.9
Replicate #2
Replicate #1
I
Residual mean POLR2A
r = 0.87
J
Mean EU (A.U.)
Mean POLR2A-GFP (A.U.)
ELK1
Scrambled
EXOSC5
K
PGAP3
Scrambled
SNRNPF

**Figure S8, related to Figure 4. (A)** Example images showing heterogeneity in POLR2A abundance across a population of MRC5 POLR2A-GFP cells, measured by either direct imaging of GFP, or immunofluorescence (IF) of POLR2A, after EU metabolic RNA labelling. **(B)** Scatter plot showing correspondence of single-cell sum nuclear intensity of GFP and POLR2A IF. Pearson's correlation inset (n=2,707). **(C)** Plate layout showing positions of control and siRNA wells in 4i experiment performed on a 384-well plate. Wells were imaged row-wise zig-zag, in the direction indicated. **(D)** Median nuclear (or cytoplasmic for PABPC1 and PABPC4) mean intensity of each imaging site in scrambled siRNA control wells, plotted as a function of image acquisition time. Asymptotic regression model fit shown is the line used to correct for intensity reduction seen during image acquisition (STAR methods). ALYREF and H3K9me3 were left uncorrected. **(E)** Comparison of well-median intensities for the two experimental replicates in 4i experiment, after data correction in B. Line shows 1:1 correspondence. Pearson's correlations inset. **(F)** Distribution of well-median intensities for all 4i markers across 63 siRNA transfections compared to equivalent distributions of scrambled siRNA controls. 4i marker shown above. N indicates nuclear mean intensity. C indicates cytoplasmic mean intensity. **(G)** Cytoplasmic, 'C' or nuclear, 'N' intensities for conditions in which the siRNAs directly target a gene encoding a protein in the antibody panel (PABPC4, XRN2), or a gene encoding a protein in the same protein complex (POLR1A, POLR2A). Mean ± s.d. for n=24 (Scrambled siRNA) or n=2 (target siRNA) wells, normalised to mean intensity in scrambled siRNA control wells. **(H)** Reproducibility of residual mean EU between replicates in the 4i experiment. Z-scored to scrambled siRNA controls. Line shows 1:1 correspondence. **(I)** As in H for residual mean POLR2A. **(J,K)** Example bivariate distributions between sum nuclear EU and POLRA-GFP in MRC5 POLR2A-GFP cells for selected example perturbations. EU and POLR2A distributions shown to the right and top, respectively.

#### Figure S9

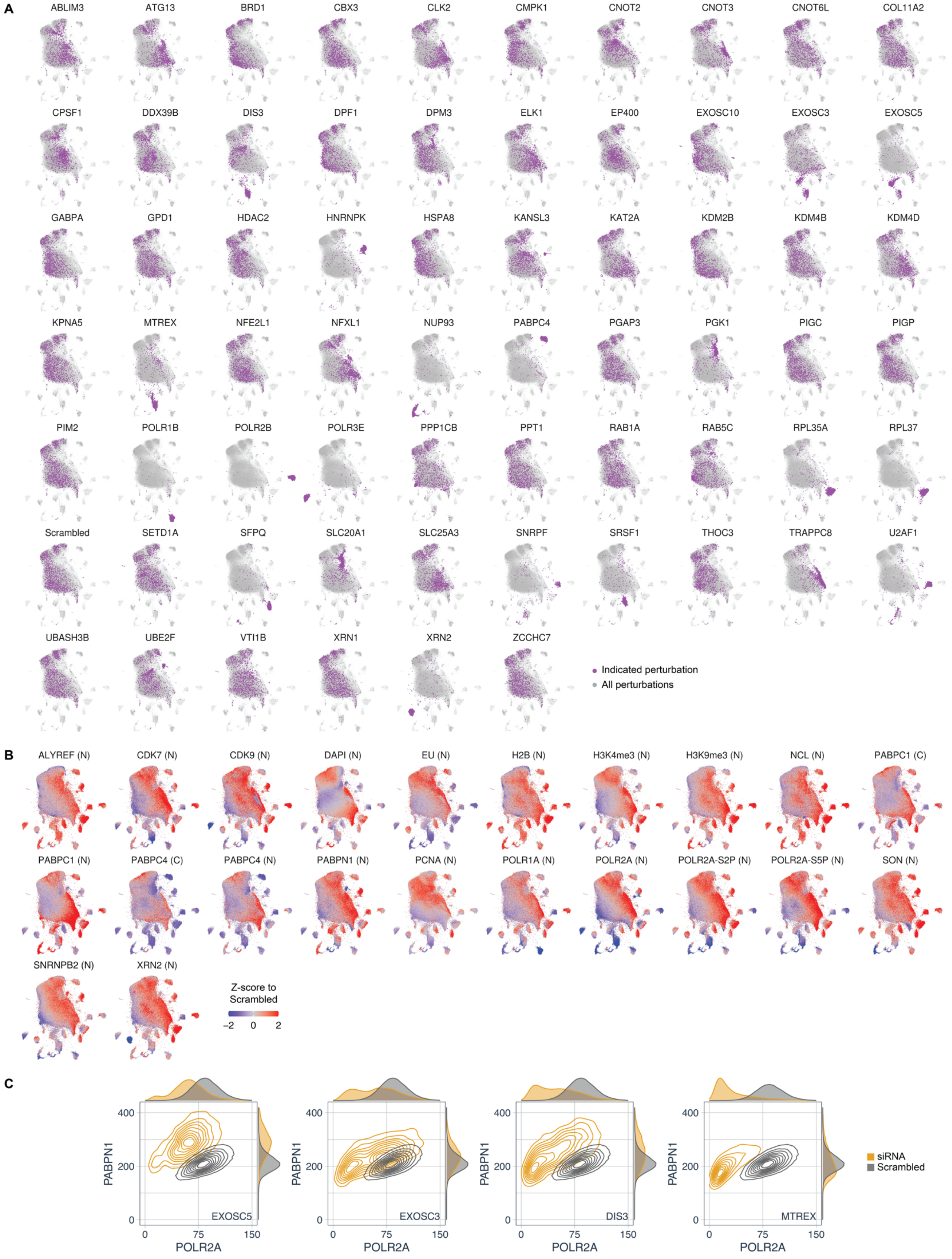
A
ABLIM3
ATG13
BRD1
CBX3
CLK2
CMPK1
CNOT2
CNOT3
CNOT6L
COL11A2
CPSF1
DDX39B
DIS3
DPF1
DPM3
ELK1
EP400
EXOSC10
EXOSC3
EXOSC5
GABPA
GPD1
HDAC2
HNRNPK
HSPA8
KANSL3
KAT2A
KDM2B
KDM4B
KDM4D
KPNA5
MTREX
NFE2L1
NFXL1
NUP93
PABPC4
PGAP3
PGK1
PIGC
PIGP
PIM2
POLR1B
POLR2B
POLR3E
PPP1CB
PPT1
RAB1A
RAB5C
RPL35A
RPL37
Scrambled
SETD1A
SFPQ
SLC20A1
SLC25A3
SNRPF
SRSF1
THOC3
TRAPPC8
U2AF1
UBASH3B
UBE2F
VTI1B
XRN1
XRN2
ZCCHC7
Indicated perturbation
All perturbations
B
ALYREF (N)
CDK7 (N)
CDK9 (N)
DAPI (N)
EU (N)
H2B (N)
H3K4me3 (N)
H3K9me3 (N)
NCL (N)
PABPC1 (C)
PABPC1 (N)
PABPC4 (C)
PABPC4 (N)
PABPN1 (N)
PCNA (N)
POLR1A (N)
POLR2A (N)
POLR2A-S2P (N)
POLR2A-S5P (N)
SON (N)
SNRNPB2 (N)
XRN2 (N)
Z-score to Scrambled
−2
0
2
C
PABPN1
POLR2A
0
200
400
0
75
150
EXOSC5
EXOSC3
DIS3
MTREX
siRNA
Scrambled

**Figure S9, related to Figure 5: UMAP of 4i experiment in genetically perturbed cells**. **(A)** Distribution of 1400 randomly chosen cells from each perturbation highlighted in purple on the UMAP, all other cells shown in grey. **(B)** Single cells in UMAP coloured by their mean intensity values, ‘N’ indicates nuclear intensity, ‘C’ indicates cytoplasmic intensity, each marker z-scored to scrambled siRNA control cells. **(C)** Bivariate distributions of single-cell mean nuclear PABPN1 and POLR2A intensity upon RNA exosome knockdown, compared to scrambled siRNA controls. Gene targeted by siRNA inset.

**Figure S10**

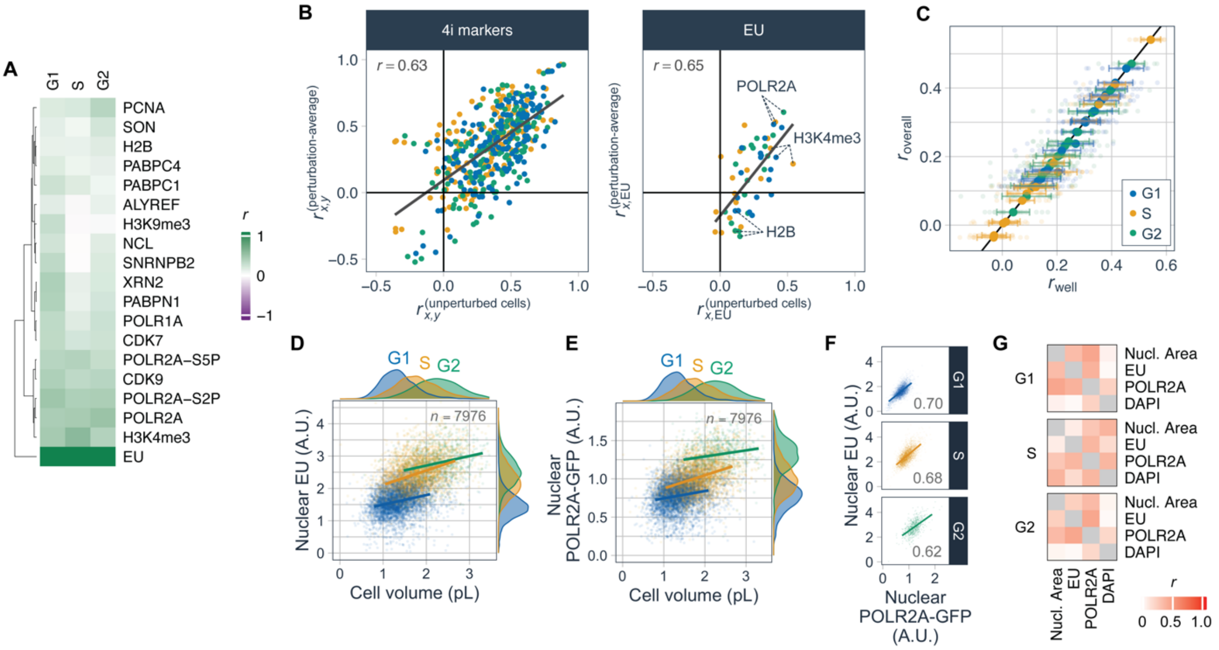

A
G1
S
G2
PCNA
SON
H2B
PABPC4
PABPC1
ALYREF
H3K9me3
NCL
SNRNPB2
XRN2
PABPN1
POLR1A
CDK7
POLR2A–S5P
CDK9
POLR2A–S2P
POLR2A
H3K4me3
EU
r
1
0
−1
B
4i markers
EU
r = 0.63
r = 0.65
POLR2A
H3K4me3
H2B
C
G1
S
G2
D
E
n = 7976
Nuclear EU (A.U.)
Nuclear POLR2A-GFP (A.U.)
Cell volume (pL)
F
0.70
0.68
0.62
Nuclear POLR2A-GFP (A.U.)
G
Nucl. Area
EU
POLR2A
DAPI

**Figure S10, related to Figure 5: (A)** Correlations of 4i marker intensities with mean nuclear EU intensity at the single-cell level in unperturbed HeLa cells, calculated for each cell-cycle phase. **(B)** Scatter plot showing the relationship between correlations of 4i marker intensities, either calculated across cells for unperturbed cells transfected with scrambled siRNA ($r_{x,y}^{\text{(unperturbed cells)}}$) or across all perturbations, after taking the mean per perturbation ($r_{x,y}^{\text{(perturbation-average)}}$). Selected markers highlighted. Points coloured by cell-cycle stage, according to legend in D. Pearson's correlation inset. **(C)** Technical validation that pooling cells from different wells does not bias correlations calculated at the single-cell level. $r_{\text{overall}}$ is the correlation calculated by pooling cells from all wells after multiplicative centring of well-medians, as shown in A. $r_{\text{well}}$ shows the mean ± s.d. of values calculated within each well before correcting and pooling across wells. Points coloured by cell cycle stage. Line shows 1:1 correspondence. **(D)** Sum nuclear EU incorporation as a function of cell volume, for different cell cycle stages in MRC5 POLR2A-GFP cells. EU and cell volume distributions are shown to the right and top, respectively. **(E)** As in D for sum nuclear POLR2A-GFP intensity. **(F)** Correlation of sum nuclear POLR2A-GFP and EU at the single-cell level in each cell-cycle stage. Pearson's correlations inset. **(G)** Partial correlations between EU, POLR2A-GFP, DNA content (DAPI) and nuclear area, in MRC5 POLR2A-GFP cells, for each cell cycle stage.

**Figure S11**

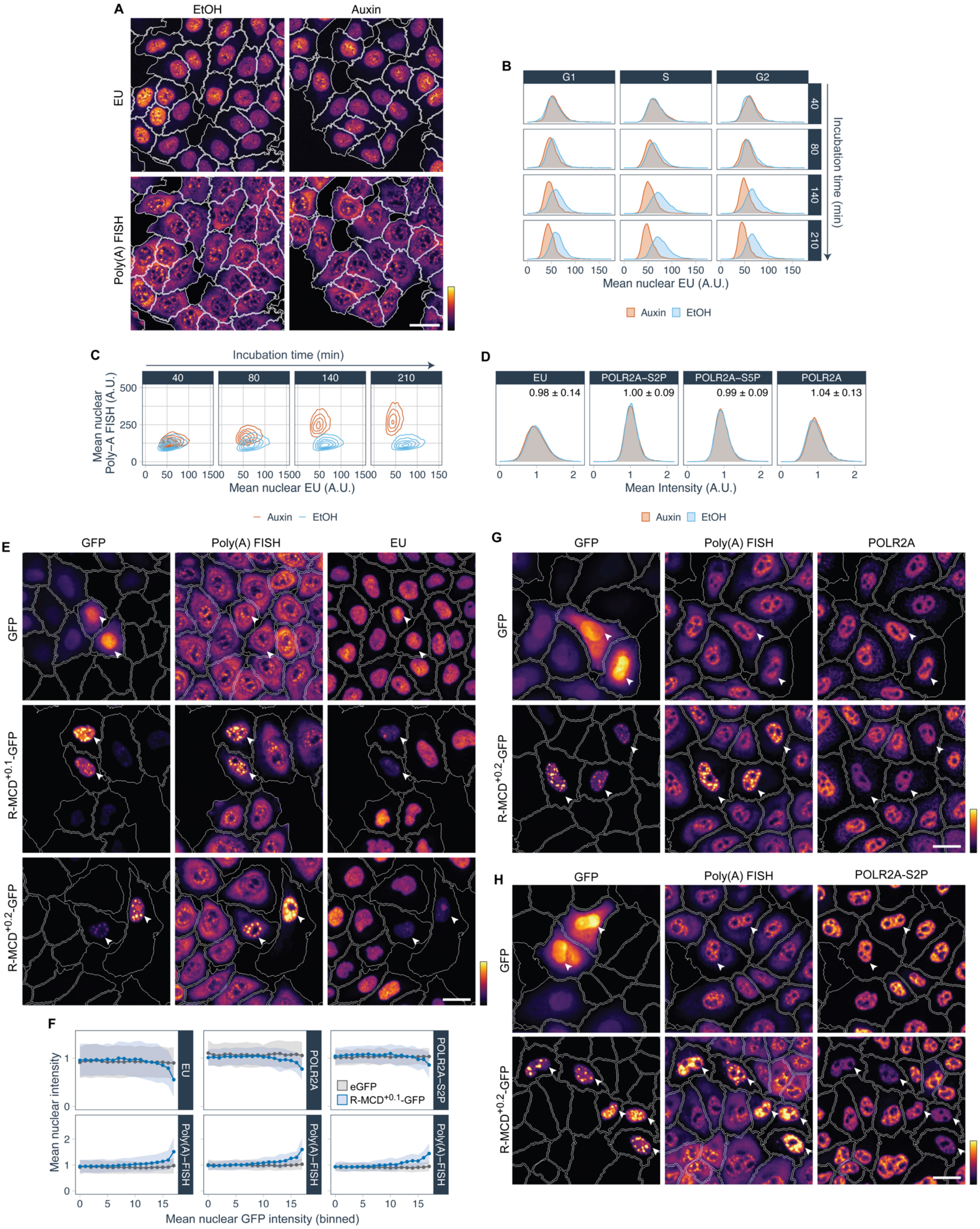

A
EtOH
Auxin
EU
Poly(A) FISH
B
G1
S
G2
40
80
140
210
Incubation time (min)
0 50 100 150
Mean nuclear EU (A.U.)
Auxin EtOH
C
Incubation time (min)
40
80
140
210
Mean nuclear Poly–A FISH (A.U.)
500
250
0
Mean nuclear EU (A.U.)
Auxin EtOH
D
EU
POLR2A–S2P
POLR2A–S5P
POLR2A
0.98 ± 0.14
1.00 ± 0.09
0.99 ± 0.09
1.04 ± 0.13
Mean Intensity (A.U.)
Auxin EtOH
E
GFP
Poly(A) FISH
EU
GFP
R-MCD+0.1-GFP
R-MCD+0.2-GFP
F
EU
POLR2A
POLR2A–S2P
Poly(A)–FISH
eGFP
R-MCD+0.1-GFP
Mean nuclear intensity
Mean nuclear GFP intensity (binned)
G
GFP
Poly(A) FISH
POLR2A
R-MCD+0.2-GFP
H
GFP
Poly(A) FISH
POLR2A-S2P
R-MCD+0.2-GFP

**Figure S11, related to Figure 6: Repression of transcription upon increased nuclear mRNA abundance. (A)** EU metabolic labelling and poly(A) FISH in HCT116:TIR1 cells (parental cells for AID cell lines) after 140 min incubation with Auxin or EtOH. **(B)** Single-cell mean nuclear EU intensity distributions for DIS3-AID cells in Auxin time-course experiment, separated into G1, S, G2 cell cycle stages. **(C)** Bivariate distributions of single-cell mean nuclear EU intensity and mean nuclear poly(A) FISH intensity for DIS3-AID cells in Auxin time-course experiment. **(D)** As in Figure 6F, but for parental HCT116:TIR1 cells. Mean nuclear intensity distributions of EU and POLR2A (immunofluorescence), for cells treated with Auxin/EtOH for 3.5h. Inset: population median for Auxin relative to EtOH controls (mean +/- s.d., n=6 (IF) or n=18 (EU) wells). **(E)** Poly(A) FISH and EU-metabolic labelling in cells expressing R-MCD$^{+0.1}$-GFP, R-MCD$^{+0.2}$-GFP or GFP control. GFP expressing cells indicated with arrows. **(F)** As in Figure 6I-K, for R-MCD$^{+0.1}$-GFP. Mean nuclear EU, or POLR2A or POLR2A-S2P (immunofluorescence), as a function of binned nuclear GFP intensity for cells expressing GFP or R-MCD$^{+0.1}$-GFP. Lower panels show mean nuclear poly(A) FISH intensity in the same cells. **(G,H)** Example images of cells transfected with R-MCD$^{+0.2}$-GFP or GFP control: poly(A)-FISH and POLR2A or POLR2A-S2P immunofluorescence, quantification in Figure 6J-K.

#### Figure S12

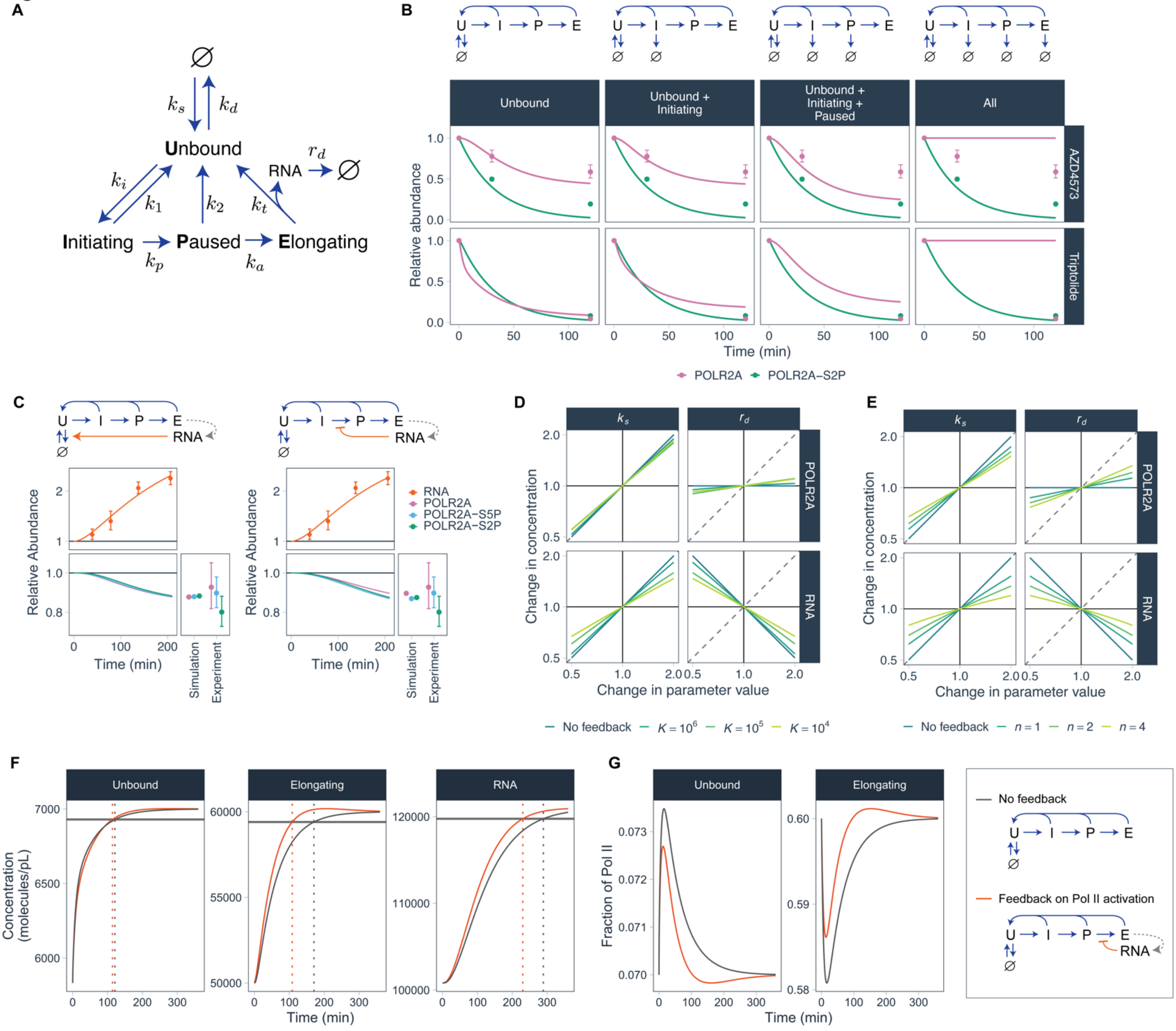

A
Unbound
Initiating
Paused
Elongating
RNA
B
Unbound
Unbound + Initiating
Unbound + Initiating + Paused
All
AZD4573
Triptolide
Relative abundance
Time (min)
POLR2A
POLR2A−S2P
C
RNA
POLR2A
POLR2A−S5P
POLR2A−S2P
Relative Abundance
Time (min)
Simulation
Experiment
D
Change in concentration
Change in parameter value
POLR2A
RNA
No feedback
E
No feedback
F
Unbound
Elongating
RNA
Concentration (molecules/pL)
Time (min)
G
Unbound
Elongating
Fraction of Pol II
Time (min)
No feedback
Feedback on Pol II activation

**Figure S12, related to Figure 7: Mathematical model of transcription with mRNA-based negative feedback. (A)** Schematic showing Pol II states in the model with arrows showing the transitions allowed and their rate constants, as defined in Table S1. Arrows to/from ∅ indicate production/degradation. **(B)** Best-fit model prediction for relative abundance of POLR2A and POLR2A-S2P upon AZD4573 or Triptolide treatment, after fitting the Pol II synthesis time-scale $k_s$ to the data shown. Four models are compared, in which the species listed above are targeted for degradation, as shown in model schematics. **(C)** Simulation of DIS3-AID depletion experiment for models including negative feedback from mRNA, as indicated in schematics above. Best-fit models shown with poly(A) FISH data from Figure 6E, and Pol II immunofluorescence data from Figure 6F. All error bars represent mean ± s.d. for all pairwise comparisons of treated and control wells. **(D)** Parameter sensitivity analysis for model with mRNA-based feedback on "paused" to "elongating" transitions: $k_a \to k_a(K/(K+R))$, for various values of $K$. x-axis indicates a multiplicative constant applied to the parameter value $k_s$ (left) or $r_d$ (right). Y-axis indicates the relative steady-state value of RNA or POLR2A which results from the parameter change. **(E)** As in D, with non-linear feedback ($k_a \to k_a(K^n/(K^n+R^n))$), with Hill coefficient $n \geq 1$ ($K = 10^5$). **(F)** Simulation of abrupt 20% cell volume increase, for models indicated in legend in F. Horizontal line indicates 99% of steady-state concentration, with vertical dashed lines indicating the time taken for models to restore Pol II and RNA concentrations to 99% of steady-state levels. Model parameters as fitted to experimental data (Table S1). **(G)** As in F, showing the fraction of Pol II in the unbound and elongating states.

**Table S1: Mathematical model parameter definitions and values.** Values are not shown if they are the same as those for the "No feedback" model.

| | | Model | | | | |
|---|---|---|---|---|---|---|
| | **Parameter** | **No feedback** | **mRNA activates Pol II decay** | **mRNA represses initiation** | **mRNA represses initiation to pausing** | **mRNA represses pausing to elongation** |
| $k_s$ | Pol II synthesis rate (molecules pL$^{-1}$ s$^{-1}$) | 3162† | 3312 | | | |
| $k_d$ | Pol II degradation rate (s$^{-1}$) | 0.452 ($k_s$/([Pol II] $\times$ 0.07))* | 0.628 | | | |
| $k_i$ | Pol II initiation rate (s$^{-1}$) | 26.96* | | 31.3 | 31.7 | |
| $k_p$ | Pol II pause rate (s$^{-1}$) | 2.397* | | | 2.88 | |
| $k_1$ | Pol II initiation-abort rate (s$^{-1}$) | 16.48 (6.87 $\times$ $k_p$)* | | | 19.8 | |
| $k_a$ | Pol II activation rate (s$^{-1}$) | 0.0792* | | | | 0.103 |
| $k_2$ | Pol II pause-abort rate (s$^{-1}$) | 0.9629 (12.16 $\times$ $k_a$)* | | | | |
| $k_t$ | Pol II termination rate (s$^{-1}$) | ($\log 2$ /1370)* | | | | |
| $r_d$ | RNA degradation/export rate (s$^{-1}$) | N/A | 0.015 | 0.015 | 0.015 | 0.015 |
| $K$ | Half-maximal repression / activation RNA concentration (molecules pL$^{-1}$) | N/A | $4 \times 10^4$ | $8 \times 10^5$ | $6 \times 10^5$ | $4 \times 10^5$ |
| [Pol II] | Total Pol II concentration (molecules pL$^{-1}$) | $10^5$* | | | | |

*Directly constrained by (Steurer et al., 2018).

†Optimised to fit the dynamics of POLR2A and POLR2A-S2P depletion upon Triptolide or AZD4573 treatment.
